# The genetic architecture of the human bZIP interaction network

**DOI:** 10.1101/2025.08.21.671354

**Authors:** A.M. Bendel, B. Eichenberger, G. Kempf, D. Klein, K. Shimada, M. Franco, S. Jaaks-Kraatz, S. Durdu, V. Herrmann, G. Roth, C. Soneson, S. Smallwood, N.H. Thomä, D. Schübeler, M.B. Stadler, F. Zenke, G. Diss

## Abstract

Generative biology holds the promise to transform our ability to design and understand living systems by creating novel proteins, pathways, and organisms with tailored functions that address challenges in medicine, sustainability, and technology. However, training generative models requires large quantities of data that captures the genetic architecture of protein function in all its complexity, but these are currently scarce. Here, we systematically mutagenized all 54 human basic-leucine zipper (bZIP) domains and quantified their interactions with each other using bindingPCA, a quantitative deep mutational scanning assay. This resulted in ∼2 million interaction measurements, capturing the effect of all single amino acid substitutions at each of the 35 interfacial positions. We found that mutation effects are largely additive in the vicinity of each wild-type bZIPs, but diverge across the family, indicating strong context dependency. A global additive thermodynamic model provided moderate prediction of mutation effects, while individual models per bZIP achieved higher performance, supporting a model of local simplicity and global complexity. Our results therefore suggest that the genetic architecture of protein function is more complex than previously anticipated, which could hinder predictability. However, a convolutional neural network trained on this dataset could accurately predict binding scores from sequence alone. Furthermore, the model enabled the design of synthetic bZIPs with high target specificity, demonstrating practical applicability for bioengineering purposes. Our study shows that capturing family-wide diversity is essential to reveal context dependencies and train accurate quantitative models of protein-protein interactions.

## Introduction

Understanding the complexity of the genetic architecture underlying protein function has been a long-standing objective, as it determines our ability to predict function from sequence and defines the scale and type of data needed to train accurate predictive models (*1–3*). At one extreme, a simple genetic architecture implies that each residue contributes independently and additively to molecular function. In this case, measuring the effects of all single mutations across the sequence would suffice to predict protein function using a simple additive model. At the other extreme, a complex architecture implies extensive context dependencies, otherwise known as epistasis, where the effect of a mutation varies depending on the sequence background. This is due to direct or long-range (allosteric) couplings between residues, such that a mutation’s effect depends on the amino acids present at other positions. A complex genetic architecture with pervasive pairwise or higher-order epistatic interactions would require large datasets that fully capture context dependencies to enable models to learn and accurately predict function from sequence.

The extent of epistasis—and thus the complexity of the genetic architecture—has been widely studied but still remains a matter of debate. Early studies measuring the effects of all pairwise combinations of mutations at up to five or six positions suggested pervasive epistasis, including higher-order interactions (*4*, *5*). However, these studies often focused on residues within enzyme active sites or other suspected interaction clusters, biasing the observed prevalence of epistasis and limiting insight into its generality across protein functions. The development of deep mutational scanning has enabled broader exploration of sequence-function landscapes using larger combinatorial libraries (*6*). For example, studies have measured all double mutants across pairs of positions in small domains (*7–9*), all 20 amino acids at four positions (*10*, *11*) or two amino acids at 16 positions (*12*, *13*). These studies found that most functional variance can be explained by additive effects and a small number of pairwise interactions, with little evidence for higher-order terms, suggesting a relatively simple genetic architecture (*1*). However, these findings are based on single proteins or a few close homologs and thus sample only narrow regions of sequence space near specific wild-types (Fig. 1A).

**Fig. 1.**
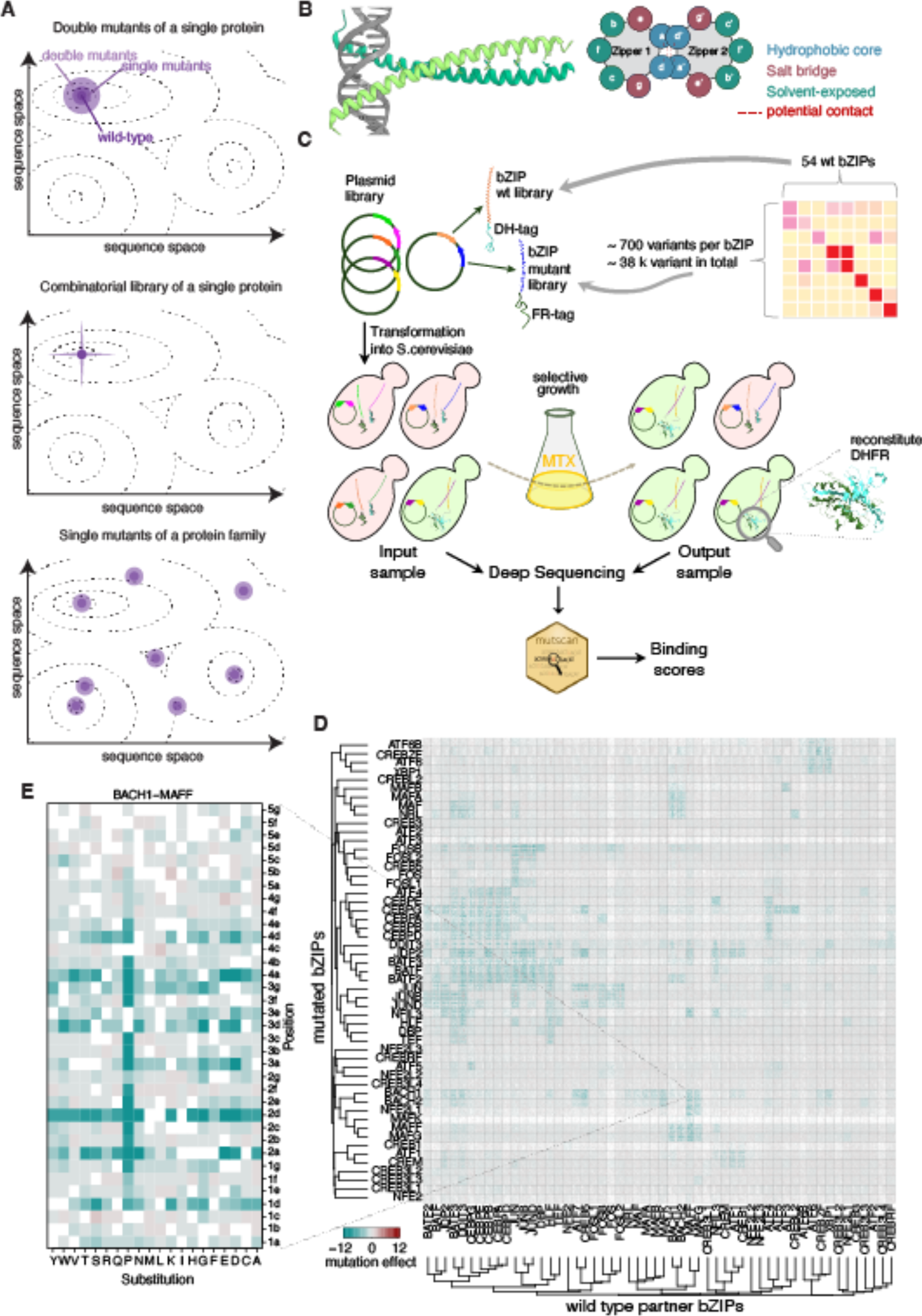
Deep mutational scanning of the entire human bZIP family. (**A**) Schematic representation of sequence space showing how deep mutational scanning in protein families (bottom) covers a wider and more diverse region of sequence space than an equivalent number of all double mutants (top) or a few multiple mutants (middle) of a single protein. x and y axes represent the sequence dimensions, dashed contours represent isochores of the quantitative function. (**B**) Members of the basic leucine zipper (bZIP) domain family dimerize by forming coiled coils (left). Repeats of seven amino acids (heptads) that form exactly two helcial turns place each of the seven amino acids lables *a* to *g* at the same position relative to the inteface within each heptads (right). (**C**) bPCA screen overview. All possible single amino acid mutants of each of 54 human bZIPs are paired with all 54 wild-type bZIP sequences and tagged with DHFR moities (DH and FR, respectively). The plasmid library containing all possible combinations of mutant and wild-type (WT) are transformed into *S. cerevisiae* and grown under methotrexate (MTX) selection. Plasmids before and after selection are extracted and barcodes are amplified for deep sequencing. Count data is processed by *mutscan* to generate a binding score for each variant-partner pair. (**D**) Heatmap of heatmaps. Each individual heatmap represents the mutation effect for each pair of mutated bZIP - wild-type partner pair (variant binding score relative to the corresponding wild-type interaction binding score). The dendrograms represent hierarchical clustering based on the Pearson correlation coefficient between each bZIP’s mutation effect profile. All variants of ATF7 were filtered out because 90% of them were missing. (**E**) Zoom on the deep mutational scanning heatmap for the interaction between mutated BACH1 and wild-type MAFF. Rows represent the substituted amino acid and columns represent the position within BACH1 bZIP. For position nomenclature, numbers represent the heptad and letters represent the position within the heptad (cf. **B**). White indicates missing data.

In contrast, protein families can contain hundreds of homologs with diverse sequences that adopt similar structures and perform the same function. For instance, the basic-leucine zipper (bZIP) domain is present in 54 human transcription factors that vary in sequence, but all form an alpha-helix and dimerize as a coiled-coil with other bZIPs (Fig. 1B) (*14–18*). Each member of a protein family therefore represents a distinct point in the broader genetic landscape (Fig. 1A). Whether mutation effects are conserved across the family-wide landscape or depend on the different contexts in distant parts of the landscape remains unknown (*19*). Comparing mutation effects across distant homologs can provide a more diverse and representative sampling of sequence space and clarify the complexity of the genetic architecture underlying protein function.

Here, we address these questions by systematically mutagenizing all 54 human bZIP domains using a deep mutational scanning assay with a quantitative protein-protein interaction read-out, bindingPCA (bPCA; Fig. 1C) (*8*). In bPCA, two proteins of interest are fused to either half of a methotrexate (MTX)-resistant dihydrofolate reductase (DHFR). Interaction between the two proteins reconstitutes DHFR activity, enabling yeast cells to grow in the presence of methotrexate. Cells expressing variants that promote dimerization grow faster and increase in relative frequency over time. Barcode sequencing before and after selection quantifies the interaction strength between each pair of bZIP variant and wild-type partner. bPCA has previously been used to study epistasis between the bZIP domains of FOS and JUN (*8*), to study the determinants of specificity of the JUN bZIP domain for the 54 human wild-type copies (*20*) and to map the energetic and allosteric landscapes of various proteins (*19*, *21–27*). Results from bPCA measurements in yeast have previously been shown to correlate strongly with measurements by splitFAST in human cells (*20*), a split fluorescent protein reporter of protein-protein interaction (*28*), and with *in vitro* measurements of binding affinity (*20–23*, *29*).

Using bPCA, we quantified the effect of all single amino acid substitutions at each of the 35 positions of the dimerization domain of each of the 54 human bZIPs, measuring binding to all 54 wild-type bZIPs for a total of ∼2 million quantitative interaction measurements (Fig. 1C). In addition to characterizing specificity and evolvability across the family, we found that mutation effects are largely additive in the vicinity of each wild-type protein. However, the same mutation can have different effects in distant homologs, indicating that sequence context becomes increasingly important across broader regions of sequence space. As a result, a simple additive model describes the family-wide landscape with only moderate accuracy. Nonetheless, a convolutional neural network accurately predicts bZIP interactions from sequence alone, but decreases in performance when holding out entire subsets of the family for training. Measuring mutation effects across an entire protein family is therefore essential for capturing the diverse context dependencies required to train accurate predictive models of protein function.

## Results

### Deep mutational scanning of the entire human bZIP interaction network

To quantify the effects of amino acid substitutions on binding profiles for all 54 human bZIPs, we mutated each of the 35 positions (or 34 for ATF1 and CREB1) in every bZIP domain to all other 19 amino acids and a premature stop codon. For each of the ∼38,000 resulting bZIP variants, we measured binding to each of the 54 wild-type bZIPs in triplicate, yielding 2,370,890 quantitative binding scores (precision-weighted log fold change between input and output samples (*30*, *31*) including 1,604,024 of the 2,041,956 expected pairs after quality filtering (all variants of ATF7 were excluded due to poor library representation, resulting in mutation data for 53 bZIPs; Table S1; see Methods). Input and output read counts were respectively highly correlated across replicates, confirming the high reproducibility of the assay (Fig. S1). Interactions among the 2,916 wild-type pairs recapitulated the known modular organization of the bZIP network (*29*), with subfamilies clustering together and preferentially interacting with partners of the same subfamily (Fig. S2).

Mutation effects relative to each of these 2,916 wild-type interactions can be represented as one heatmap of as many variant effect heatmaps, which can be clustered based on the profile of mutation effects across partners (Fig. 1D, E). Detectable interactions display characteristic mutation patterns, with high sensitivity at key interfacial residues, such as hydrophobic positions, and uniformly deleterious effects of proline across positions (Fig. 1D, E). In contrast, many wild-type interactions show little response to mutation, indicating that their dimer concentration fall below our dynamic range. This is reflected in the distribution of mutation effects, which is tightly centered around zero for non-interacting pairs (Fig. S3). Out of 1,601,098 variants for which a mutation effect relative to wild-type could be calculated, 82,485 showed a significant difference to the wild-type interaction (5.2%; FDR=5%; absolute mutation effect > 2), with predominantly negative effects (5,578 vs. 76,907 for positive and negative effects, respectively, or 6.8% vs. 93.2% of all significant variants, respectively). There is a strong relationship between the binding score of a wild-type interaction and the number of mutations that significantly decrease it (Fig. S4), suggesting that the latter could be used as a more robust metric to determine whether two proteins do or do not interact. Interestingly, the majority of mutations with positive effects enable new interactions with partners that the wild-type does not bind (partners for which none of the mutations significantly decrease the wild-type binding score). We identified 3,230 such cases (57.9% of all variants with positive effects), corresponding to 1,418 new bZIP interaction pairs (59.3% of all pairs for which no mutations significantly decrease the wild-type binding score). Therefore, the majority of bZIP pairs are on the edge of binding, which can be promoted by a single mutation.

### Specificity profile rewired by single mutants

To identify specificity-rewiring mutations, we compared the interaction profiles of each mutant to that of its corresponding wild-type bZIP. We found 169 mutants whose binding profile is entirely distinct from the wild-type (see Methods). For example, wild-type CREB5 binds JUN, JUND and CEBPG, but substituting an asparagine with leucine in the hydrophobic core of the third heptad abolishes these interactions while enabling new interactions with BACH1 and BACH2 (Fig. S5). Such neofunctionalizing mutations account for 0.46% of all mutations and are found in 28 out of the 53 bZIPs with an average of 6 per bZIP. This indicates that bZIPs can frequently acquire new interaction specificities through a single mutation, which likely contributes to the high evolvability of the family and supports its expansion by gene duplication followed by neofunctionalization (*32*).

### A global additive model does not predict mutation effects accurately

In the absence of context dependencies, mutations are expected to act additively on the free energy of binding. However, bPCA binding scores are proportional to dimer concentration, which is a nonlinear function of binding energy (Fig. 2A). To compare effects across different sequence contexts, we first need to convert binding scores to the latent free energy dimension. We previously showed that, since bZIPs are unfolded as monomers and fold upon binding, a two-state thermodynamic model can accurately predict binding changes due to JUN mutations across partners (*20*). We therefore fitted a global additive model (*21*, *33*) to the whole dataset (excluding stops) that considers only the first order, context-independent effect of mutations (Fig. 2B). The model fits a pseudo Gibbs free energy of binding for a baseline wild-type interaction (ΔG_ref; JDP2-DDIT3), a pseudo change in Gibbs free energy of binding for each sequence differences relative to JDP2 (ΔΔG_mut), and a pseudo change in Gibbs free energy of binding for replacing DDIT3 by each of the 53 remaining wild-type partners (ΔΔG_bZIP). The total pseudo ΔG for any variant-partner pair is computed by summing the relevant terms, which is then transformed to fraction bound using a two-state equation. A global linear transformation is used to predict the observed binding score from the fraction bound.

**Fig. 2.**
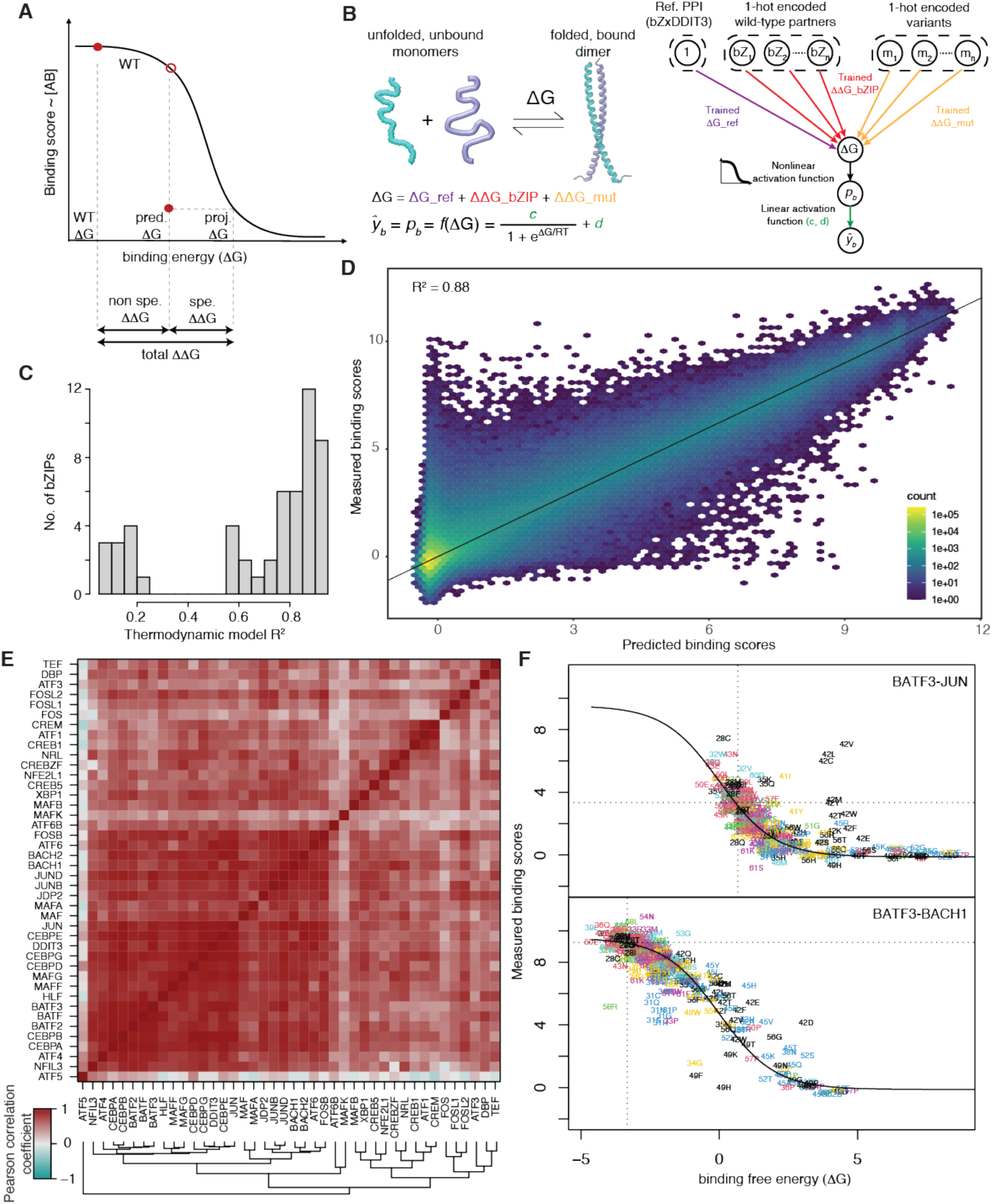
A simple additive thermodynamic model fitted on each mutated bZIP independently shows that mutation have mostly additive effects in the vicinity of each wild-type bZIP. (**A**) Binding scores are proportional to dimer concentration but non-linearly related to binding energy. The total change in binding free energy (ΔΔG) of a mutation is composed of a non-specific ΔΔG representing the partner-independent mutation effect (the ΔΔG_mut parameter fitted by the thermodynamic model), and a component representing the ΔΔG specific to a given partner. (**B**) bZIPs are present at an equilibrium of unfolded, unbound monomers and folded, bound dimers. ΔΔGs for each mutant can thus be inferred using a simple two-state thermodynamic model. (**C**) Distribution of thermodynamic models performance (R^2^ between predicted and measured binding scores) when fitted separately for each mutated bZIP. (**D**) Correlation between predicted and measured binding scores when pooling all individual thermodynamic models. (**E**) Clustered heatmap of Pearson correlation coefficients between fitted ΔΔG_mut values for the same substitutions across all bZIP pairs. (**F**) Measured binding scores as a function of binding free energy ΔG for BATF3 mutants vs. wild-type JUN and wild-type BACH1 as examples. The sigmoidal fit represents predicted binding scores from fitted ΔG (ΔG_ref + ΔΔG_bZIP + ΔΔG_mut). Residuals from the fit represent mutation effects specific to a given partner. Variants are colored by their position within a heptad to highlight that high residuals correspond to mutations at the same positions.

This global additive model performed moderately well (R² = 0.47; Fig. S6; Table S1), suggesting that non-additive effects, i.e. context dependencies, are prevalent across the family-wide landscape.

### The interaction profile of individual bZIPs is well predicted by a simple additive model

The performance of the global model suggests that the same mutation can have different effects in different bZIP backgrounds. To test this, we fitted a separate two-state model for each bZIP, using the dataset of all 681 variants of that bZIP (excluding stops) paired with the 54 wild-type partners. This yielded individual ΔΔG_mut values for each mutation in each of the 54 bZIPs, allowing for direct comparison of mutation effects across the family. Each model also includes ΔΔG_bZIP values for the 53 non-reference wild-type partners (using DDIT3 as the reference partner in all cases).

Model performance varied across bZIPs, with R² values ranging from 0.02 to 0.94 (Fig. 2C; Table S1). However, this variation largely reflects differences in data quality, driven by differences in binding score distributions. Since weak binding scores have inherently lower accuracies due to reduced read depth, datasets from bZIPs with a lower number of binding partners are typically noisier. For instance, the model for FOS yielded a low overall R² of 0.56, but restricting evaluation to strongly interacting partners (JUN, JUNB and JUND) increased performance to R² = 0.82. There is indeed a strong relationship between the average binding score of each mutated bZIP and their thermodynamic model performance (Spearman rho=0.93; Fig. S7). Pooling predictions from all 54 bZIP-specific models gave a higher overall R² (0.88; Fig. 2D) than the global model. Altogether, these results support our hypothesis that mutation effects are mostly additive in the vicinity of the wild-type proteins but differ across bZIPs.

### Fitted additive mutation effects are similar across distant bZIPs

To further compare mutation effects across bZIPs, we computed Pearson correlation coefficients between ΔΔG_mut values for the same substitution across all bZIP pairs. These correlations ranged from –0.58 to 0.93, with an average of 0.42 (Fig. S8A). However, low correlations are mostly due to bZIPs with bad MoCHI fits, i.e. bZIPs with low average binding scores (Fig. S7), as removing bZIPs with R^2^ below 0.4 increases the minimum correlation coefficient to –0.23 and the average correlation to 0.62 (Fig. S8B).

Hierarchical clustering based on the Pearson correlation across ΔΔG_mut profiles does not recapitulate well sub-family relationships (Fig. 2E). However, since each ΔΔG_mut term is fitted on the interaction with all 54 partners, it captures only the non-specific, average effect of a mutation across all partners. This non-specific component of mutation effects reflects intrinsic biophysical properties, such as α-helical stability and hydrophobicity, which we previously termed “zipper propensity” (*20*). Deviations from the model (residuals) reflect partner-specific effects and reveal specificity-determining residues (Fig. 2A; Fig. 2F).

### Patterns of specificity differ across distant bZIPs

Our data allows studying the sequence determinants of specificity at the level of the whole family. To capture these partner-specific effects, we generated specificity maps for each bZIP, showing the mutation residuals across all partners (Fig. 3A, with full dataset in Fig. S9). Specificity scores are narrowly centered around 0 with long extending tails on both sides (Fig. 3B). Mutations having strong positive effects on specificity are more numerous than thos having strong negative effects (272 and 48, respectively, at thresholds of specificity scores of 6 and –6 respectively). bZIPs from the same subfamily tend to use similar specificity determinants. For example, MAFF and MAFG share specificity patterns, as do BATF2 and BATF3 (Fig. S9). Interestingly, distant bZIPs can share some specificity determinants despite overall divergence. This indicates that different bZIPs reuse different elements from a common pool of specificity-determining residues in distinct combinations to generate unique interaction profiles (Fig. 3C).

**Fig. 3.**
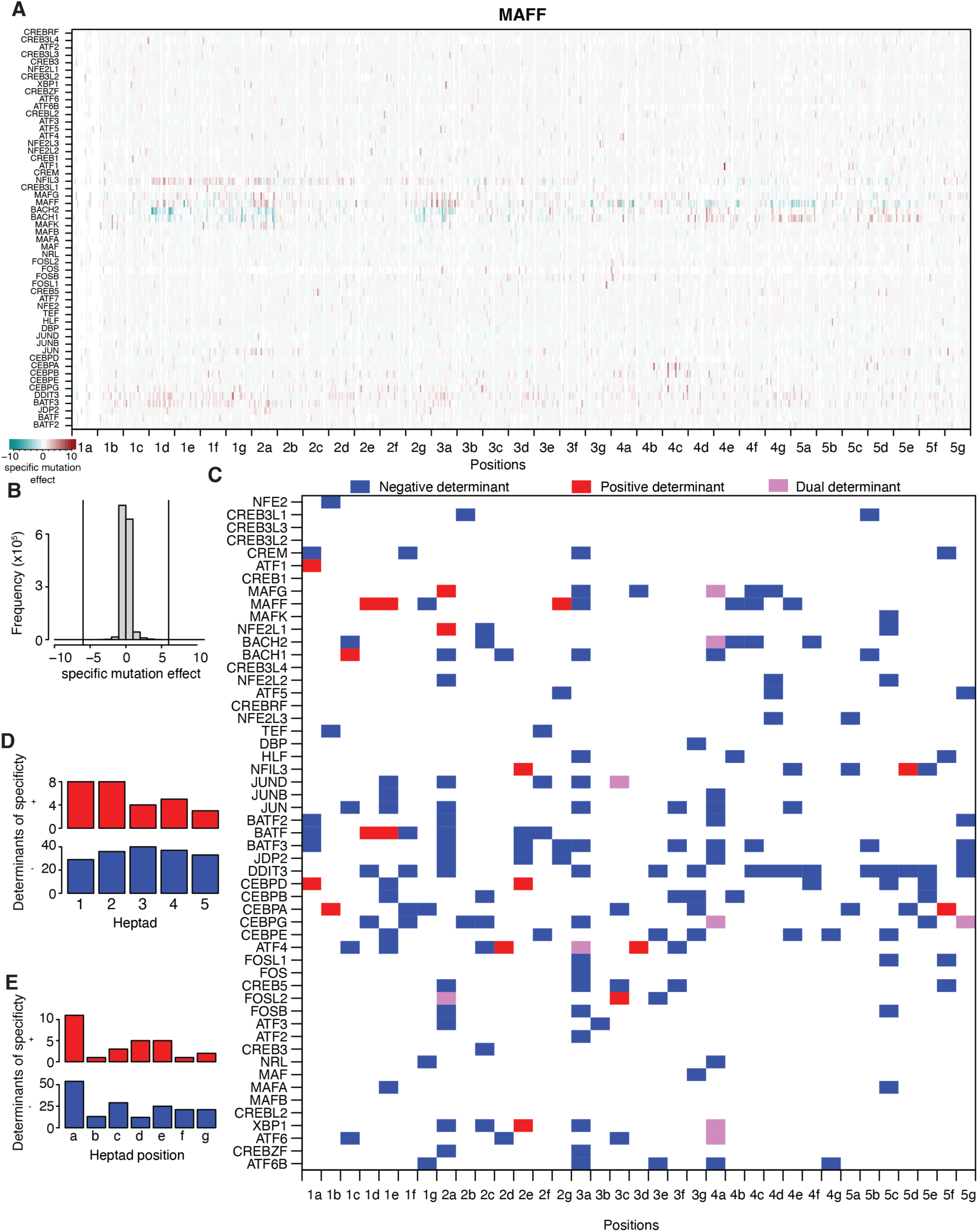
Family-wide mapping of bZIPs determinants of specificity. (**A**) Heatmap of specific mutation effects (i.e. residuals from thermodynamic model) for MAFF. Row represent the 54 wild-type partners. Columns represent each of the 20 amino acids at each of the 35 positions. Negative specific effects (turquoise) indicate that the wild-type residue serves as a negative determinant of specificity. Positive specific effects (red) indicates that the wild-type residue serves as a negative determinant of specificity. (**B**) Distribution of specific mutation effects across all bZIPs. (**C**) Determinants of specificity for all bZIPs. Positions where at least one substitution has a strong specific effect with one partner were considered determinants of specificity. Dual determinants (pink) correspond to position that can serve as both a negative or positive detemrinant of specificity. (**D**) Number of determinants of specificity per heptad across all bZIPs. Blue indicates negative determinants of specificity, red indicates positive ones. (**E**) Number of determinants of specificity per heptad position across all bZIPs. Blue indicates negative determinants of specificity, red indicates positive ones.

The central heptad contains more negative determinants of specificity than the ones at the extremity of the domain, while positive determinants are more frequently found in the first two heptads (Fig. 3D). Position *a* at the hydrophobic core of heptads (Fig. 1B) is the position most frequently involved in determining specificity, both positively and negatively (Fig. 3E).

### The genetic architecture of bZIP interaction is complex

To assess the overall complexity of the landscape, we compared the full mutation effect profiles (total ΔΔG; Fig. 2A) across all informative bZIPs (with thermodynamic model R^2^ > 0.4, Fig. S7). High correlation between profiles indicates that mutation effects are conserved, whereas low correlation implies strong context dependencies. Related bZIPs such as MAFF and MAFG or BATF2 and BATF3 have similar profiles (R = 0.69 and 0.73, respectively; hamming distance 13 and 34, respectively; Fig. 4AB). In contrast, distant bZIPs such as MAFF and BATF2 show weaker correlations (R = -0.25; hamming distance 48; Fig. 4C). The correlation coefficients range from –0.91 to 0.80, with an average of 0.17, lower than for the fitted ΔΔG_mut. Clustering the full correlation matrix reveals a modular structure, with bZIPs from the same subfamily forming highly correlated clusters and unrelated subfamilies showing little correlation (Fig. 4D). Altogether, these results support a view where the genetic architecture is locally simple, with mutations having similar effects in similar bZIPs, but globally complex, with mutation effects differing between distant bZIPss.

**Fig. 4.**
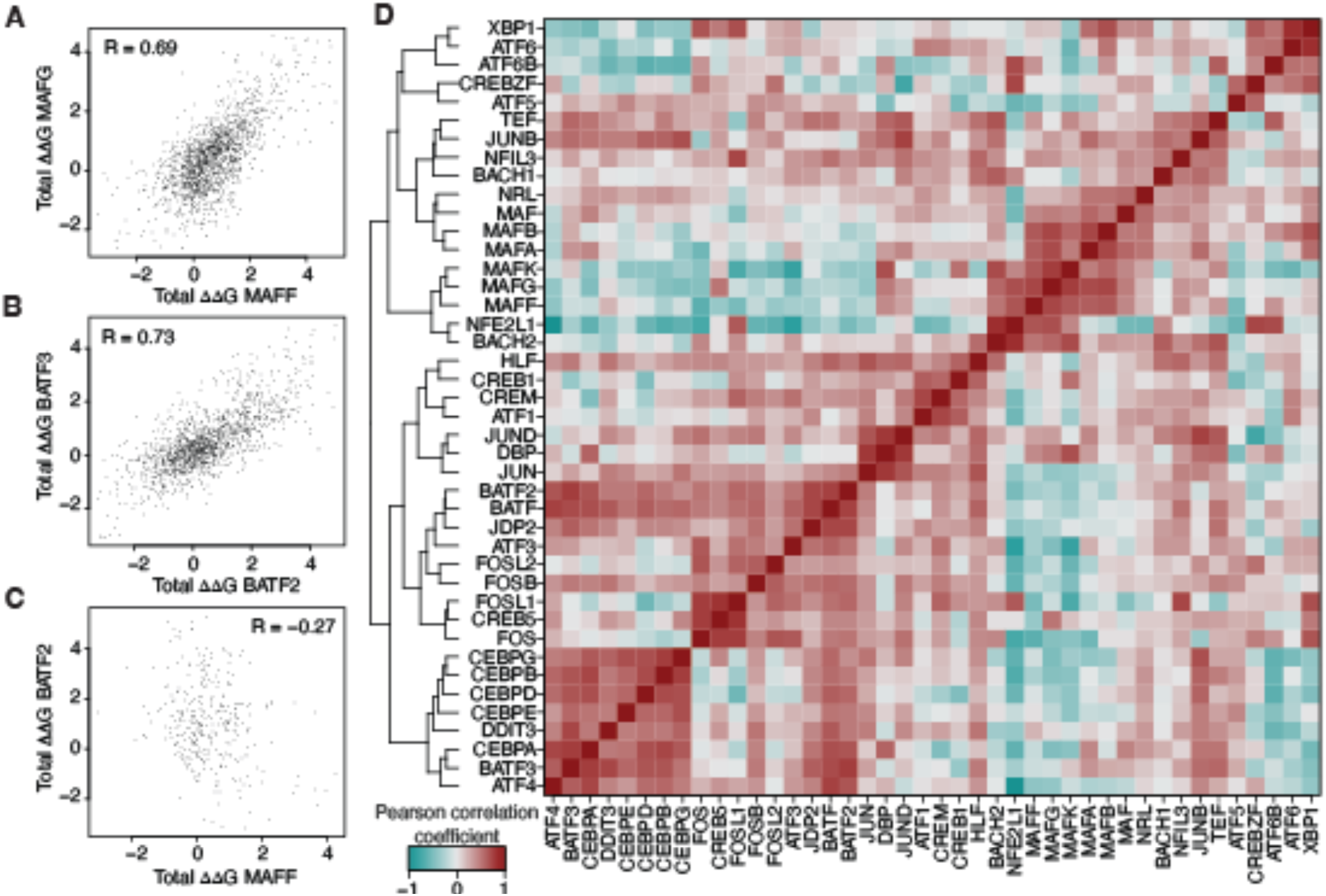
The same mutation has similar effects in closely related bZIPs but different effects in distant bZIPs. (**A-C**). Comparison of mutation effects (total ΔΔG) for the same mutations between related (**A, B**) or distant bZIPs (**C**) bZIPs. Pearson correlation coefficients (R) are indicated. (**D**) Clustered heatmap of Pearson correlation coefficients between projected total ΔΔG values for the same substitutions across informative bZIP pairs (i.e. for which the R^2^ of the thermodynamic model was above 0.4, since the fitted value for low performance bZIPs are unreliable). Only variants for which the projected ΔG as well as the one from the corresponding wild-type variant are between –3 and 3 were considered to avoid very deep projection at the plateaux of the sigmoid.

### A simple convolutional neural network predicts binding scores accurately

To evaluate the predictability of this complex landscape, we trained a simple convolutional neural network with four 1D convolutional layers (see Methods and Fig. 5A for details) to predict binding scores from the one-hot encoded sequences of bZIP variant-partner pairs. The data was randomly split into training (80%), test (10%), and validation (10%) sets. The model reached high accuracy (test R² = 0.88, Fig. 5B; Table S2), demonstrating that despite global complexity, the bZIP sequence-function landscape is highly predictable with appropriate models.

**Fig. 5.**
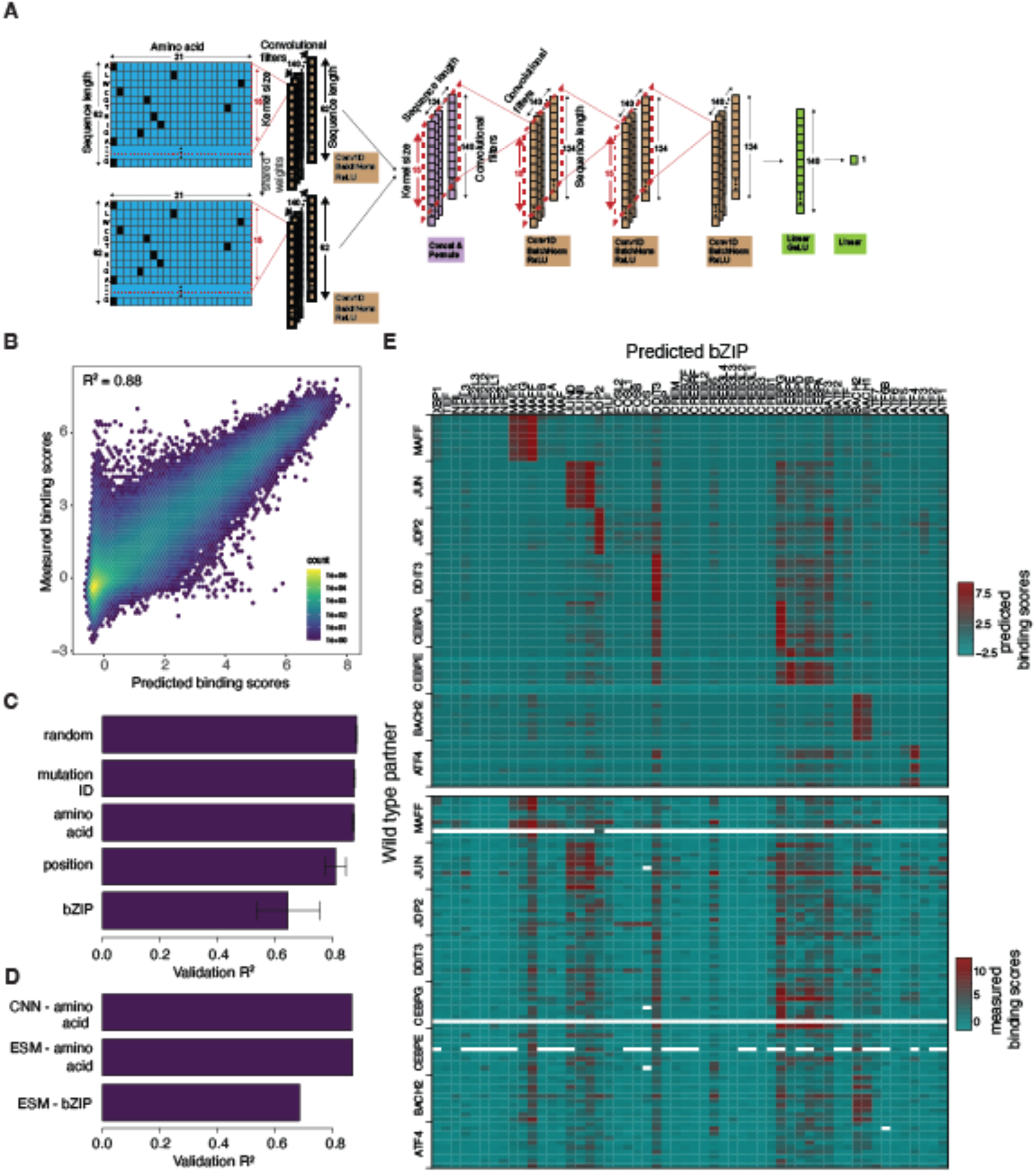
A sequence-function model for the bZIP family. (**A**) Architecture of the convolutional neural network (CNN) trained to predict binding scores from sequence. Dimensions of each layer are indicated (see Methods for more details). (**B**) Correlation between predicted and measured binding scores shows high predictive accuracy (R^2^ = 0.88). (**C**) Comparison of model performance on the test set for different splitting strategies between train, test and validation sets (80%, 10% and 10% of the data, respectively). Error bars represent the 95% confidence interval 10-fold cross validation. (**D**) Model performance after using an ESM2 model (15B parameters) for exclusion of entire bZIPs or specific amino acids. Due to high computational resources requirements, a single representative fold was performed. Performance of the CNN was also evaluated on the single fold for the amino acid split for comparison purpose. (**E**) Predicted (top) versus measured (bottom) binding scores for synthetic bZIPs generated by a preliminary (see Method) CNN model trained on the amino acid split using a genetic algorithm.

To guide future experimental design, we tested how different training strategies affect model performance. First, we excluded mutations toward randomly chosen amino acids at specific positions in specific bZIPs. For instance, the model never saw mutations to arginine at position 3 in JUN during training and was evaluated on these held-out mutations. We also tested stricter exclusions, such as removing all mutations to arginine at position 3 in any bZIP, or removing all mutations at a specific position across bZIPs. In all cases, model performance remained similar to the fully random split, with a small drop in performance in the latter case (position split, test R² = 0.81; Fig. 5C); Table S2. However, when we excluded all data for a randomly chosen subset of bZIPs (e.g. the model never saw any mutations in JUN or CEBPB), performance decreased (test R² = 0.63 on the held-out bZIPs; Fig. 5C; Table S2). This suggests that when entire regions of sequence space are missing from training, the model cannot learn their specific context dependencies. Nonetheless, using a fine-tuned, pre-trained large language model (ESM2, 15B parameters, (*34*); see Methods) improved predictions for held-out bZIPs (test R² = 0.75 vs. 0.88 for amino acid split; Fig. 5D; Table S2), likely because the model had already learned general protein features from large-scale training.

### Prediction of new synthetic sequences that target specific bZIPs

To demonstrate the practical utility of our model, we used a genetic algorithm to design synthetic bZIPs that bind specific targets with high specificity. For each of eight targets, we ran ten parallel optimization trajectories, starting from the wild-type sequence with the best initial specificity and penalizing off-target interactions. We then experimentally tested the interaction of these 80 synthetic bZIPs with all 54 wild-type-partners using bPCA in triplicate. The experimental results matched the predicted binding scores closely (Fig. 5E; Table S3). Notably, these synthetic bZIPs differ by 16 until up to 59 amino acids from all natural human bZIPs, showing that the model can extrapolate to novel regions of sequence space when trained on a sufficiently broad and diverse dataset.

## Conclusions

By systematically mutagenizing all 54 human bZIPs, we generated a comprehensive dataset that captures both conserved and divergent context-dependencies of mutation effects across the family. The resulting 54 specificity maps show that different bZIPs rely on distinct combinations of shared specificity-determining positions to encode their interaction profiles (Fig. 3C).

A single global additive model, where a single term is fitted for each substitution, explains only 47% of the variance in binding scores. Fitting an individual model on each mutated bZIP, allowing a different effect to be estimated for the same substitution in different bZIPs, explains 88% of the variance in binding scores across the whole dataset. The good performance of the individual models compared to the low performance of the single model supports a view where local simplicity coexists with global complexity. The difference in variance explained by these two modelling approaches suggests that first-order additive effects and context-dependencies respectively represent 47% and 41% of the variance in binding scores. The remaining 12% of unexplained variance consists in specific mutation effects that change binding to one or a few partners specifically (Fig. 2F and Fig. 3A,C), as well as measurement error.

Our convolutional neural network trained on this dataset accurately predicts binding scores from sequence alone, demonstrating the utility of sampling a wide and diverse genetic landscape to learn regional context-dependencies. While the performance does not exceed the one of the ensemble of the individual thermodynamic models, it allows extrapolating to mutations not seen during training (Fig 5C). The thermodynamic modelling approach, despite having lower number of free parameters, requires data on each mutation during training to be able to fit its free energy parameter. These results also suggest that accurate prediction does not require sampling all amino acid substitutions at all positions, potentially enabling smaller and more targeted libraries.

However, this may rely on the repetitive heptad structure of bZIPs, where the same substitution at equivalent positions across heptads provides redundant information for learning. Whether this strategy applies to protein domains without such a convolutional structure remains to be tested.

Moreover, as large language models improve, they may further enhance prediction accuracy in previously unsampled regions of sequence space. Our dataset will therefore provide a useful resource for the development of new models that aim at predictive mutation effects. Extending this approach to other protein interaction domain families, i.e. sampling the global genetic landscape of protein-protein interactions, will enable training more general sequence-function models applicable to diverse protein-protein interactions.

## Acknowledgments

We thank Ben Lehner, Albert Escobado, Christian Landry and Luca Giorgetti for comments on the manuscript. ChatGPT4 was used only to correct language.

## Author contributions

Conceptualization: GD, AMB

Methodology: GD, AMB, FZ, BE, GK, MBS, CS, GR, KS, SJK

Investigation: AMB, GD, FZ, BE, GK, MBS, CS, GR, SJK, KD, KS, SD, VF, MEF, SS

Visualization: GD, AMB, GK

Funding acquisition: GD, FZ, DS

Project administration: GD

Supervision: GD, FZ, MBS, DS, NT

Writing – original draft: GD, AMB

Writing – review & editing: GD, AMB, GK, CS, MBS, FZ

## Data and materials availability

Raw fastq files have been deposited in GEO with accession code XXX. Table S1, S2 and S8 have been deposited at Zenodo (https://doi.org/10.5281/zenodo.16913737). All code to analyze data and generated all figures is available on GitHub (https://github.com/gdiss/bZIP). All code related to deep learning model training and synthetic bZIP desig is avilable on GitHub (https://github.com/fmi-basel/Deep-learning-bZIPs)

## Material and Methods

### Yeast Strain

All experiments were performed in BY4742 (MATα *his3*Δ1 *leu2*Δ0 *lys2*Δ0 *ura3*Δ0).

### Media and buffers used

- LB: 25 g/L Luria-Broth-Base (Invitrogen, Waltham, MA, USA), autoclaved 20 min at 121 °C.
- LB-agar with 2X Ampicillin (100 μg/mL): 25 g/L Luria-Broth-Base (Invitrogen, Waltham, MA, USA), 7.5 g/L Agar, 1.2 g/L MgSO_4_ · H_2_O. Autoclaved 20 min at 121 °C. Cool-down to 45 °C. Addition of 100 mg/L Ampicillin.
- YPAD: 20 g/L Bacto-Peptone, 20 g/L Dextrose, 10 g/L Yeast extract, 25 mg/L Adenine. Filter-sterilized (Millipore Express ®PLUS 0.22 μm PES, Merck, Darmstadt, Germany).
- SC-ura: 6.7 g/L Yeast nitrogen base without amino acids, 20 g/L glucose, 0.77 g/L complete supplement mixture drop-out without uracil. Filter-sterilized (Millipore Express ®PLUS 0.22 μm PES, Merck, Darmstadt, Germany).
- SC-ura/ade/met: 6.7 g/L Yeast nitrogen base without amino acids and folic acid, 20 g/L glucose, 0.74 g/L complete supplement mixture drop-out without uracil, adenine and methionine. Filter-sterilized (Millipore Express ®PLUS 0.22 μm PES, Merck, Darmstadt, Germany).
- SORB: 1 M sorbitol, 100 mM LiOAc, 10 mM Tris-HCl pH 8.0, 1 mM EDTA pH 8.0. Filter-sterilized (Millipore Express ®PLUS 0.22 μm PES, Merck, Darmstadt, Germany).
- Plate mixture: 40 % PEG3350, 100 mM LiOAc, 10 mM Tris-HCL pH 8.0, 1 mM EDTA pH 8.0. Filter-sterilized (Millipore Express ®PLUS 0.22 μm PES, Merck, Darmstadt, Germany).
- Recovery medium: YPAD + 0.5 M sorbitol. Filter-sterilized (Millipore Express ®PLUS 0.22 μm PES, Merck, Darmstadt, Germany).
- Competition medium: SC-ura/ade/met + 200 μg/mL methotrexate (BioShop Canada Inc., Canada), 2 % DMSO.
- DTT buffer: 0.1 M EDTA-KOH pH7.5, 10 mM DTT
- Zymolyase buffer: 20 mM K-phoshpate pH 7.2, 1.2 M sorbitol, 0.4 mg/mL Zymolyase 20T (amsbio, USbiological), 100 μg/mL RNAse A
- Home-made Buffer P1: 50 mM Tric-HCl pH 8, 10 mM EDTA, 0.1 mg/mL RNAse A
- Home-made Buffer P2: 200 mM NaOH, 1 % SDS
- TAE buffer: 0.4 M Tris, 0.01 M EDTA, 0.2 M Acetic Acid

### Library cloning

#### Overview cloning strategy

In order to allow reusing the library for other projects, each of the 37,814 bZIP variant had to be paired with itself as a DH and FR fusion (henceforth referred to as bait and prey, respectively) to form an intermediate library. To construct the final library, the preys are removed from the plasmid and replaced with an insert coming from a library consisting of the 54 wild-type partners. The intermediate library is constructed by cloning oligonucleotides ordered form Twist Bioscience (San Francisco, CA, USA; Table S4). However, because the oligonucleotide synthesis is limited to 300 nt, only the zipper domains were encoded, plus the four last nt of the DNA binding domain (DBD), which is directly upstream of the zipper domain. These four nucleotides are left as overhangs after digesting by type IIs restriction enzymes, which allows cloning the oligonucleotide library in a pre-constructed library containing only the wild-type DBD of each of the 54 bZIPs on both the bait and prey side in tail-to-tail orientation, separated by the two type IIs recognition sites.

#### Plasmid backbone construction

Since the plasmid library construction required many cloning steps and different combinations of restriction enzymes, our standard deepPCA plasmid (*20*) could not be used as a backbone, and some positions had to be altered by silent mutations. The backbone was optimized to remove certain restriction sites that would be required for subsequent cloning steps. It carries the two DHFR-cassettes with CYC promoters and a barcode landing pad. The resulting plasmid was named pDL00260.

The backbone was assembled from five individual fragments in yeast. Two of the fragments were synthetic DNA molecules (gBlocks) ordered from Twist Bioscience (San Francisco, CA, USA). gBlock 1 (gBlock_URA3) for the bait side, a 3x Myc tag and directly adjacent the first 34 bases of the DH-fragment open reading frame (ORF). gBlock 2 (gBlock_DHFR) carried the rest of the DH cassette including the entire DH ORF and 6 bases of the 3x Myc tag to create a 40 bp homologous overhang between the two gBlocks. Next to the DH ORF, there was a 60 bp linker sequence called Linker L3 (GSAGASAGGSGSAGSASGGS), a BamHI restriction site, 12 random base pairs as placeholders, and an NheI restriction site. Adjacent began the FR cassette encoded in opposite sense of the DH cassette. First, there was another 60 bp linker sequence called Linker L4 (GASGSAAGGSGSAGSGASAS), followed by the FR fragment ORF, a 3x Flag tag and the CYC promoter for the prey side. Next to the CYC promoter was an SP2 primer binding site for binding of the Illumina reverse standard sequencing primer, followed by a 259 bp random spacer sequence. This spacer was designed to create enough distance between the two Illumina sequencing primer binding sites, so that the resulting PCR amplicon would have the ideal size for efficient clustering on Illumina flow cells. Next to the spacer was a wild-type prey placeholder barcode of 7 bp length (CTGATGT), then an NcoI restriction site and a variant bait placeholder barcode of 22 bp length (ACTAGCTAGTACTATCTAACCG), a KpnI restriction site, the SP1 primer binding site for the binding of the Illumina forward standard sequencing primer, an annealing site for the M13-rev primer for Sanger sequencing and 81 more bases of the backbone. Both gBlocks were digested with BsaI (New England Biolabs, Ipswich, MA, USA) to remove the adapters added by the manufacturer. 1.5 μg of each gBlock were digested in a 60 μl reaction setup with 1X Cut Smart Buffer (New England Biolabs, Ipswich, MA, USA) and 1 μl of restriction enzyme for 1 hour at 37 °C. The digested fragments were gel-purified from a gel made from 1 % Agarose in 1 X TAE Buffer using the QIAquick Gel Extraction Kit (QIAGEN, Hilden, Germany).

The remaining three fragments were PCR-generated by performing PCRs on a lab-created plasmid called pDL00005 that was derived from pGD009(*8*). The first amplicon was amplified using primers oDL00743 and oDL00744 (Table S5). It carried a 37 bp overlap with the second gBlock (gBlock_DHFR), the ColE1 origin and the last 165 bp of the Ampicillin resistance cassette. The second amplicon was amplified using primers oDL00741 and oDL00742 (Table S5) and carried a 41 bp overlap with the first amplicon, the remaining Ampicillin resistance cassette, and the CEN-ARS. The final PCR fragment which was amplified with primers oDL00745 and oDL00746 (Table S5) consisted of a 40 bp overhang with the second amplicon, the first half of the URA3 gene and a 40 bp overhang with the second half of the URA3 gene present on the first gBlock.

The five fragments were co-transformed into yeast BY4741 cells for *in vivo* recombination. A 5 mL culture of BY4741 in YPAD was grown overnight at 30 °C and 200 rpm shaking. From this culture, a 10 mL pre-culture of YPAD was inoculated at OD_600_ 0.3 and grown for 4 hours until an OD_600_ of 1.2 – 1.6 was reached. The cells were harvested for 2 min at 3,000 g and first washed with 15 mL of sterile H_2_O and then with 15 mL of SORB before they were resuspended in 400 μl of SORB. 10 μl of 10 mg/mL freshly boiled salmon sperm DNA (Agilent Technologies, Santa Clara, CA) were added, and the mixture was incubated on a wheel for 30 min at room temperature (RT). Afterwards, 500 μl of Plate Mixture were added to 100 μl of the competent cells together with an equimolar mixture of 0.1 pmol of each of the five prepared fragments. This was once more incubated on a wheel for 30 min at RT. 75 μl of DMSO were added to the mixture, and the cells were heat-shocked for 20 min in a 42 °C water bath and then harvested for 3 min at 800 g. The pelleted cells were resuspended in 100 μl of YPAD medium and recovered for 1 hour at 37 °C. Finally, 10 μl and 90 μl of the recovered culture were plated on SC-ura agar plates and incubated at 30 °C. After two days of growth, four colonies were picked and grown to saturation in 20 mL of liquid SC-ura overnight. The cells were harvested, and DNA was extracted using the MasterPure™ Yeast DNA Purification Kit (LGC Biosearch Technologies, Hoddesdon, UK) according to the manufacturer’s protocol. The DNA extracts were then transformed into NEB 10-beta Electrocompetent *E.coli* (New England Biolabs, Ipswich, MA, USA) according to the manufacturer’s protocol. The cells were recovered for 1 hour at 37 °C after which 1/10 and 9/10 of the culture were spread onto LB + 2X Ampicillin agar plates and incubated over night at 37 °C. Eight colonies were picked for verification. A final clone was Sanger-sequenced using multiple primers to cover the entire plasmid and ensure that there were no point mutations. The sequence-verified plasmid was named pDL00260.

#### Variant library design

The sequences (DBD + Zipper) of all 54 human bZIPs were obtained from Uniprot according to the bZIP annotations (Table S6). Only amino acids 1 to 62 were kept (61 for ATF1 and CREB1 since they are at the C-terminus of the full-length proteins), corresponding to the 27 amino acids DNA binding domain and the 35 amino acids (five heptads) of the zipper domain. They were sequence-optimized to remove any restriction sites required for cloning. Additionally, the versions that were to be cloned in the bait site were altered with synonymous mutations to make them as different from the wildtype sequence as possible, while considering known codon usage from *S.cerevisiae.* This way, issues with PCR and cloning of inverted repeat sequences were avoided. For each bZIP, every possible single amino acid substitution plus a stop at each position in the 35 amino acid-long zipper domain was designed. The resulting library consisted in 701 variants per bZIP (35 position x 20 substitutions + one wild-type sequence; 681 for ATF1 and cREB1) for a total 37’814 variants (701 x 52 + 682 x 2). Oligonucleotides encoding these variants were ordered from Twist as matrixed 300 nt-long oligonucleotide pools with a capacity of 121 variant per well (Twist Bioscience, San Francisco, CA, USA; Table S4). For each bZIP, the 701 (or 681) variants were distributed over 6 wells of a 385-well plate. Two zipper sequences were encoded in a tail-to-tail orientation so that each oligonucleotide carried the same variant twice, the one on the prey side with the reference sequence (only optimized to remove restriction site), and the one on the bait side with the alternative sequence. Only the zipper domain and the four last nucleotide of the DBD (directly upstream of the zipper domain) were encoded. PCR-handles and the corresponding restriction sites were added up- and downstream of the paired zippers to enable subsequent amplification and cloning.

#### Intermediate DNA-binding domain library

To clone the plasmids carrying the bait and prey DBD into which the oligonulcoetide library will be cloned, 54 gBlocks carrying tail-to-tail DBDs of the same bZIP, one with the reference sequence (prey side), one with the alternative sequence (bait side), separated by a 13 bp spacer and flanked by NheI and BamHI restriction sites, were ordered from Twist Bioscience (San Francisco, CA, USA; Table S7).

To clone the intermediate DBD library, plasmid pDL00260 was digested with NheI-HF and BamHI-HF (New England Biolabs, Ipswich, MA, USA) in 1X Cut Smart Buffer (New England Biolabs, Ipswich, MA, USA) and de-phosphorylating with Quick CIP alkaline phosphatase (New England Biolabs, Ipswich, MA, USA) at 37 °C for one hour to create the backbone. All 54 gBlocks were digested with these same enzymes without de-phosphorylation. The ligations were performed in the same way as for the intermediate wild-type prey library (see above), and the final clones were sequence-verified with Sanger sequencing.

#### Intermediate bZIP variants libraries

For the intermediate bZIP variants libraries, an individual variants library was cloned for each of the 54 bZIPs. The oligonucleotides carrying the mutated zipper domains were ordered lyophilized in a 384-well plate, each variants of one bZIP were distributed over 6 individual wells (Table S4), and each well contained 2.244 ng or 24.37 fmol of oligonucleotides. The oligonucleotides were resuspended in 5 μl of Buffer EB (QIAGEN, Hilden, Germany) and incubated at RT for 30 min to completely solubilize. Then, all 6 wells per bZIP were pooled, and 15 μl per pool corresponding to approximately 73 fmol of oligonucleotides were used as templates for PCRs to make the oligonucleotides double stranded and add random 22 bp molecular barcodes at the 5’ end. PCRs were setup in 25 μl reactions using the KAPA HiFi HotStart DNA Polymerase Kit (Roche Sequencing Solutions Inc, Pleasanton, CA) and primers oDL00789 and oDL00763 (Table S5). The PCR was run for 20 cycles with an annealing step of 63 °C for 15 sec and an extension step of 72 °C for 15 sec. Afterwards, the reactions were treated with 10 μl of ExoSAP-IT (Thermo Fisher Scientific, Waltham, MA) at 37 °C for 15 min to remove residual single stranded oligonucleotides, followed by heat inactivation at 80 °C for 15 min. The samples were then purified using the MinElute PCR Purification Kit (QIAGEN, Hilden, Germany), eluted in 16 μl of Buffer EB and immediately digested with KpnI-HF and BsaI-HFv2 (New England Biolabs, Ipswich, MA, USA) in 1X Cut Smart Buffer (New England Biolabs, Ipswich, MA, USA) at 37 °C overnight and heat-inactivated for 20 min at 80 °C in the morning. The digestion setups were again purified with the MinElute PCR Purification Kit (QIAGEN, Hilden, Germany) and eluted in 12 μl of Buffer EB. The yield was measured with a Qubit system (Invitrogen, Waltham, MA, USA), and 2 μl per sample were loaded on an analytical gel to verify successful PCR and subsequent digest.

The backbones of the libraries were prepared by individually digesting the 54 intermediate DBD plasmids from the previous step with KpnI-HF and BsaI-HFv2 (New England Biolabs, Ipswich, MA, USA) in 1X Cut Smart Buffer (New England Biolabs, Ipswich, MA, USA) and de-phosphorylating with Quick CIP alkaline phosphatase (New England Biolabs, Ipswich, MA, USA) at 37 °C overnight. They were gel-purified using the QIAquick Gel Extraction Kit (QIAGEN,

Hilden, Germany), and the yield was measured with Qubit (Invitrogen, Waltham, MA, USA). Backbones and their respective insert library were then ligated with NEB T4 ligase (New England Biolabs, Ipswich, MA, USA) according to the manufacturer’s protocol using 70 fmol of backbone and 141 fmol of insert. The reaction was incubated overnight in a PCR cycler using the temperature-cycle ligation (TCL) protocol to increase ligation efficiency (4 cycles of 10 min at 22 °C and 10 min at 16 °C, 49 cycles of 30 sec each at 4 °C, 27 °C, and 13 °C, 10 cycles of 30 sec at 4 °C, 1 hour at 16 °C, 10 min at 22 °C, 10 min at 16 °C, 49 cycles of 30 sec each at 4 °C, 27 °C, and 13 °C, finally 30 sec at 4 °C, 1 hour at 16 °C, 1 hour at 22 °C and 3 h at 18 °C). In the morning, the samples were dialyzed against approximately 50 mL of MilliQ H_2_O (Merck Millipore, Darmstadt, Germany) to reduce the buffer salt concentration using nitrocellulose membranes with 0.025 μm-sized pores (Merck Millipore, Darmstadt, Germany). The dialyzed setups were transformed into NEB 10-beta Electrocompetent *E.coli* (New England Biolabs, Ipswich, MA, USA) according to the manufacturer’s protocol. The cells were recovered for 30 min to prevent an overestimation of transformants due to cell division. For quantification of the transformants, the recovered culture was diluted 100-fold and 1 μl, 10 μl and 100 μl of this dilution were plated on LB + 2X Ampicillin. To harvest between 10,000 and 20,000 transformants (corresponding to that same number of barcodes) per bZIP, the remaining recovery culture was plated in different amounts, namely two times each 10 μl, 20 μl, 50 μl, and 100 μl in LB + 2X Ampicillin. All plates were incubated at 37 °C overnight, and the number of transformants was counted in the morning. According to that number, plates with the estimated number of colonies corresponding to 10 – 20,000 clones were harvested off the plates by adding sterile H_2_O to the plates and then scraping off the cells with a plastic spatula. This would correspond to an average of around 20 barcodes per variant. The harvested cells were spun down at 3,000 g for 15 min, washed once with sterile H_2_O and re-suspended in 50 mL of liquid LB + 4X Ampicillin. They were grown for approximately 6 more hours, then harvested at 5,000 g for 15 min, and the library was extracted using the QIAGEN Plasmid Plus Midi Kit (QIAGEN, Hilden, Germany).

An initial quality control for the barcode-variant association (see next section) revealed that all but eight (ATF1, ATF2, ATF3, ATF6B, BACH1, NFE2, NFE2L1, NRL) of the cloned libraries showed a strong sequence bias in their barcode sequence towards Thymine bases. This resulted in low barcode complexity that could eventually lead to a strong reduction in our data quality. Therefore, the affected 46 libraries were re-cloned. The sequence bias originated from the primer oDL00789 (Table S5) which was initially ordered as PAGE-purified. When re-ordered as simply desalted, there was no strong sequence bias in the barcodes.

To re-clone the biased libraries, the backbones (intermediate DBD library) were digested with KpnI-HF and BamHI-HF (New England Biolabs, Ipswich, MA, USA) in 1X Cut Smart Buffer (New England Biolabs, Ipswich, MA, USA) and de-phosphorylating with Quick CIP alkaline phosphatase (New England Biolabs, Ipswich, MA, USA) at 37 °C overnight and gel-purified with the QIAquick Gel Extraction Kit (QIAGEN, Hilden, Germany). To create the re-barcoded inserts, 2 ng from the biased libraries of the first cloning attempt were used as template for PCRs using the KAPA HiFi HotStart DNA Polymerase Kit (Roche Sequencing Solutions Inc, Pleasanton, CA) and primers oDL00789 (new order) and oDL00331 (Table S5). The PCR was run for 15 cycles, annealing at 60 °C for 15 sec and extension at 72 °C for 30 sec. The product was purified with the MinElute PCR Purification Kit (QIAGEN, Hilden, Germany) and eluted in 16 μl of Buffer EB. Afterwards, all amplicons were digested with KpnI-HF, BamHI-HF and additionally DpnI in order to remove remaining template plasmid (New England Biolabs, Ipswich, MA, USA) in 1X Cut Smart Buffer (New England Biolabs, Ipswich, MA, USA) and de-phosphorylating with Quick CIP alkaline phosphatase (New England Biolabs, Ipswich, MA, USA) at 37 °C overnight and gel-purified with the QIAquick Gel Extraction Kit (QIAGEN, Hilden, Germany). Backbones and inserts were ligated as described above.

#### Barcode-variant association

For each of the 54 sub-libraries, two rounds of PCR were performed to add the Illumina adapters to the amplicons. A region covering the barcodes and the bZIP variants was amplified with the KAPA HiFi HotStart DNA Polymerase Kit (Roche Sequencing Solutions Inc, Pleasanton, CA) in 25 μl reaction setups. In the first round, primers oDL00345 and oDL00518 (Table S5) and 1 ng of plasmid template corresponding to almost 200 Mio molecules were used. The PCR was run for 10 cycles, annealing at 60 °C for 15 sec and extension at 72 °C for 30 sec. Afterwards, the samples were treated with 2 μl per 5 μl reaction of ExoSAP-IT (Thermo Fisher Scientific, Waltham, MA) as described above and an analytical gel was run to check if the PCR was successful. The samples were then purified using the QIAEX II Gel Extraction Kit (QIAGEN, Hilden, Germany). 90 μl Buffer QX1, 1.5 μl of beads and 10 μl of 3M NaOAc were added to 30 μl of sample. The samples were incubated at RT for 10 min and then spun down for 2 min at 4,000 rpm. The beads were washed twice with 100 μl of Buffer PE and then dried and re-suspended in 20 μl of Buffer EB. They were incubated for 45 min at RT to ensure complete elution and finally spun down again. 17 μl of eluate were recovered for each sample and used as template for the second round of PCR that was performed in 100 μl setups again using the KAPA HiFi HotStart DNA Polymerase Kit (Roche Sequencing Solutions Inc, Pleasanton, CA) and primers oDL00345 and oDL00346 (Table S5). The PCRs were run for 12 cycles, annealing at 72 °C for 15 sec and extension at 72 °C for 30 sec. All samples were gel-purified with the MinElute Gel Extraction Kit (QIAGEN, Hilden, Germany). The columns were washed three times and eluted with 20 μl of Buffer EB. The concentration was determined by Qubit (Invitrogen, Waltham, MA, USA) and all 54 samples were pooled at equimolar ratios. The final pool was once more purified using AMPure XP beads (Beckman Coulter, Brea, CA, USA). The beads were added to the pool at a 1:1 volume ratio, mixed and incubated for 5 min, transferred onto a magnetic rack, and incubated for another 5 min. They were then washed twice with 200 μl of freshly prepared 70 % Ethanol and each time incubated for 30 sec before discarding the Ethanol. Then, the pellet was left to dry completely and resuspended in Buffer EB. The elution volume was slightly lower than the original sample volume to concentrate the pool. The resuspended beads were incubated at 37 °C for 10 min, placed back on the magnetic rack and left for another 5 min at RT before the eluate was removed. For a final quality control, a qPCR was run using the KAPA Library Quantification Standards with Primer (Roche Sequencing Solutions Inc, Pleasanton, CA) and then submitted for sequencing on the Illumina NovaSeq 6000 Sequencing System (Illumina, San Diego, CA) using an SP flow cell and 2 x 250 nt paired-end sequencing. 699.5 million reads were obtained.

Barcode-variant association was performed as described in (*20*). For pre-processing of the reads, *mutscan v0.2.29* (*30*) was used. In short, reads with a Phred score below 20 in either the forward or reverse read were discarded. Passing reads were the merged allowing for an overlap between 85 and 132 bp with maximum 3% mismatch. Read counts were then aggregated to identical sequences. Unique sequences that did not contain a perfect NcoI or SpeI site were discarded before splitting the sequence into the barcode and variant sequence by NcoI. This yielded barcode and variant sequences of variable lengths, with a peak in read count at the expected length (22 and 371 bp for barcode and variant, respectively). At this step, we obtained a total of 453 M reads for 22 bp-long barcodes, for 800,000 expected barcodes (estimated by the total number of transformants harvested after cloning), for an average of ∼566 reads per barcode. To distinguish true unique barcode-variant pairs present in the sample from sequencing errors or PCR template switching, the total counts for each unique barcode sequence and each unique variant sequence were determined. Pairs with barcodes shorter 20 or longer than 23 bp were filtered out. Barcode-variant pairs with less than five reads were considered too low frequency to correspond to true barcode-variant pairs given the expected 566 average number of reads per pair and thus filtered out as PCR or sequencing errors. At this step, we obtained 965,689 unique barcode sequences, some of which still likely sequencing errors of true barcodes with high frequency. To identify those, we grouped all barcodes associated with the same unique variant sequence. For each unique variant, all barcodes with a Levenstein distance (to account for varying barcode length) of 4 bp or less to the most abundant barcode were filtered out. This was then repeated to the second highest frequency barcode, and iteratively until all remaining barcode sequences were processed and all true barcodes associated to this variant were identified. This yielded a final list of 885,846 true barcode sequences. Next, to identify PCR or sequencing errors in the variant sequence, the same process was used by grouping all variants corresponding to the same unique barcode sequence and filtering out all variant sequences with a Levenstein distance of 4 or less to the most frequent variant iteratively. At this stage, the variant sequence is still composed of the two tail-to-tail bait and prey variant sequences. The sequence was then split with SpeI and BamHI to isolate the bait variant sequence. Given that only the bait variant is kept during cloning of the heterodimer library (the prey variant sequence was replaced by the 54 wild-type partners), the variant sequence was cut by SpeI and BamHI to isolate the bait sequence and by HindIII and BsmBI to isolate the prey sequence. For each unique bait and prey sequence, the closest of the 54 wild-type bZIP was identified as the one with the closest Hamming distance. We identified 116 bait variants were the DNA binding domain and zipper domain map to different wild-type bZIPs. These probably arose during cloning of the library and were flagged as chimera. Finally, for barcodes that were associated with more than one variant, the barcode-variant pairs were filtered out if the frequency was lower or equal to 1% of the frequency of the most abundant pair. These originate from template switching during PCR for the sequencing library preparation and are not present in the plasmid library. For each barcode, the Hamming distance to the closest barcode was calculated. This will allow filtering reads coming from ambiguous barcodes when analyzing bindingPCA sequencing data. The final set consisted of 900,539 barcode-variant pairs (Table S8). The substitutions in the variant sequence were then identified, and those that did not correspond to expected variants according to the oligonucleotide pool design were flagged. 752,029 pairs corresponded to expected bait variant sequences. This corresponds to a median of 25 barcodes per variant.

#### Intermediate wild-type partners library

The final library is designed to have all the 37,814 variants on the bait side (DH-tagged) and the wild-type bZIPs on the prey side (FR-tagged). An intermediate library of the 54 wild-type partners cloned in the prey site and barcoded using 7 bp-long predetermined barcode sequences wast thus constructed by cloning each of the 54 plasmids individually. 54 gBlocks were ordered carrying the 7 bp barcode, the spacer, the SP2 binding site, a CYC promoter, and the full FR cassette consisting of a 3XFlag tag, the FR fragment ORF, Linker L4 and one of the 54 wild-type bZIPs (DBD and Zipper; Table S9). The gBlocks were flanked by NcoI and HindIII restriction sites.

To clone the intermediate library, pDL00260 was digested with NcoI-HF and HindIII-HF (New England Biolabs, Ipswich, MA, USA) in 1 X Cut Smart Buffer (New England Biolabs, Ipswich, MA, USA) and de-phosphorylating with Quick CIP alkaline phosphatase (New England Biolabs, Ipswich, MA, USA) at 37 °C for one hour to create linearized backbone. All 54 gBlocks were digested with these same enzymes without de-phosphorylation. All digested DNA was gel-purified with QIAquick Gel Extraction Kit (QIAGEN, Hilden, Germany). Afterwards, 54 individual ligations were performed using NEB T4 ligase (New England Biolabs, Ipswich, MA, USA) according to the manufacturer’s protocol using 16 fmol of backbone and 31 fmol of insert. The ligation setups were incubated for 10 min at RT and then transformed into NEB 10-beta chemocompetent cells (New England Biolabs, Ipswich, MA, USA) using a small-scale plate transformation protocol. 10 μl of bacteria were pipetted in each of 54 wells of a 96 well plate that was kept on ice. 2 μl of respective ligation mixture were added to each well and the plate was transferred into a pre-cooled PCR cycler (4 °C). The cells were then incubated at 4 °C for 30 min, heat-shocked at 42 °C for 30 sec, and then incubated at 4 °C again for another 5 min. Immediately after, 100 μl of NEB 10-beta Stable Outgrowth Medium (New England Biolabs, Ipswich, MA, USA) were added to each well, and the cells were recovered at 37 °C in the PCR cycler for an hour. The recovered cells were then spread on LB + 2X Ampicillin plates using one plate per setup and incubated over night at 37 °C. All plasmids were sequence-verified using Sanger sequencing.

#### Final variant – wild-type partners library

For practical reasons, the final library was split into four batches that each consisted of 13 or 14 bZIP variant sub-libraries (Table S10). The JUN sub-library was shared across all batches for normalization. The batches were assembled so that each batch should have approximately the same number of on average strong, intermediate, and weak interactors. These estimations were made based on interaction measurements from (*20*). All bZIP variant sub-libraries per batch were pooled at equimolar ratios and 20 μg of the pools were digested in 200 μl setups with NcoI-HF and SpeI-HF (New England Biolabs, Ipswich, MA, USA) in 1X Cut Smart Buffer (New England Biolabs, Ipswich, MA, USA) and de-phosphorylating with Quick CIP alkaline phosphatase (New England Biolabs, Ipswich, MA, USA) at 37 °C overnight. To ensure proper digestion, 12 μl of each enzyme and phosphatase were used. In the morning, the reaction was heat-inactivated for 20 min at 80 °C, and the backbone fragment was gel-purified with the QIAquick Gel Extraction Kit (QIAGEN, Hilden, Germany). For the inserts, all intermediate wild-type prey plasmids were pooled at equimolar ratio and digested the same way as the backbones but without de-phosphorylation. The batches were assembled in a large-scale 100 μl ligation setup using 200 fmol of backbone, 600 fmol of insert, 10 μl of NEB T4 ligation buffer, 5 μl of NEB T4 ligase (New England Biolabs, Ipswich, MA, USA). The setups were split in two times 50 μl, incubated over night with the TCL program (see above), pooled again in the morning and dialyzed in aliquots of 33 μl as described above for approximately 2 hours. The dialyzed aliquots were combined again and concentrated to 20 μl with a Speedvac. They were then transformed into NEB 10-beta Electrocompetent *E. coli* (New England Biolabs, Ipswich, MA, USA) according to the manufacturers protocol. For each batch, four transformations were performed and pooled together. The cells were recovered for 30 min and then spread on LB + 2X Ampicillin plates with 15 cm diameter. Approximately 400 μl of recovered culture were plated per 15 cm-plate. Additionally, 1 μl and 10 μl of a 100-fold dilution of the recovered culture were spread on LB + 2X Ampicillin plates of 10 cm diameter to quantify the transformation efficiency. All plates were incubated overnight at 37 °C. In the morning, the libraries were harvested from the 15 cm-plates as described above. They were not grown further but immediately purified using the NucleoBond® PC 500 kit (Macherey-Nagel, Hilden, Germany). Between 7.6 M and 10 M clones were harvested for the four batches. As quality control, test digests were performed using restriction sites distinct from the ones used for cloning. All batches showed a clean band pattern of expected sizes. All four library batches had a size of approximately 500,000 bZIP pairs.

### bindingPCA

Each batch was individually screened in three biological replicates.

#### Large-scale yeast transformation

For each transformation, a colony of BY4742 was used to inoculate 20 mL YPAD and grown over night to saturation. In the morning, a pre-culture of 700 mL of pre-warmed YPAD was inoculated at an OD_600nm_ of 0.3 and grown for ca. 4.5 hours until an OD_600nm_ between 1.2 and 1.6 was reached. The cells were then harvested for 10 min at 3,000 g, washed with 50 mL of sterile H_2_O and afterwards with 50 mL SORB. Between the washed cells were spun down for 5 min at 3,000g. After the second wash, the cells were resuspended in 28 mL of SORB and incubated on a wheel at RT for 30 min. 700 μl of salmon sperm DNA (Agilent Technologies, Santa Clara, CA) were then added and each sample was mixed thoroughly. Afterwards, 4 μg of library DNA were added to each sample followed by thorough mixing again. Each sample was then distributed over 4 tubes in aliquots of 7 mL and 35 mL of Plate Mixture were added to each tube. The samples were once more incubated at RT on a wheel for 30 min after which 3.5 mL of DMSO were added to each tube followed by thorough mixing. The cells were then heat-shocked at 42 °C in a water batch for 20 min during which the tubes were mixed by inversion every few minutes. Afterwards, the cells were harvested for 5 min at 3,000 rpm (1,811 g), the supernatant was thoroughly removed with a vacuum pump, and each pellet was resuspended in 50 mL of Recovery Medium and incubated for one hour at 30 °C without agitation. The recovered cells were harvested for 5 min at 3,000 g and each 4 tubes per transformation were pooled by resuspending in SC-ura liquid medium and pipetting all in one tube. After spinning down again, each pellet was used to inoculate 1.4 L of SC-ura in 5 L Erlenmeyer flasks. 10 μl and 50 μl from the big culture were spread on SC-ura agar plates to quantify transformation efficiency. The cultures were incubated for 48 hours at 30 °C and 200 rpm shaking.

#### Competition screen

After 48 hours of transformant selection, a second selection cycle was started by inoculating 2 L of SC-ura/ade/met at OD_600nm_ of 0.1 and letting the cells grow for 12 hours until ca. An OD_600nm_ of 1.2. From each sample, the amount of cells required to inoculate 2 L of competition medium at an OD_600nm_ of 0.05 was harvested and used to inoculate the competition culture. The remaining cells were harvested, washed twice with sterile H2O and frozen at -20 °C until further processing. These were the Input samples. Once the competition culture had reached an OD600 of 1.6, the entire culture was harvested, washed twice and frozen. These were the Output samples.

### DNA extractions and plasmid quantification

Frozen Input and Output sample pellets were thawed at RT, resuspended in 40 mL DTT buffer, and incubated at 30 °C for 15 min and 200 rpm shaking. The cells were harvested for 5 min at 2,500 g, resuspended in 40 mL Zymolyase buffer and incubated for 2 hours at 30 °C and 200 rpm shaking. The spheroblasts were collected for 5 min at 2,500 g. 15 mL of home-made Buffer P1 were added to the cells which were resuspended thoroughly by pipetting. Next, 15 mL of home-made Buffer P2 were added, the tubes were mixed by inversion, and the mixture was incubated for 10 min at RT. Then 15 mL of pre-cooled (4 °C) commercial Buffer P3 (QIAGEN, Hilden, Germany) was added to the cells, and the samples were centrifuged for 5 min at 3,000 g and 4°C after which the supernatant was transferred to 50 mL centrifugation tubes (Nalgene, Rochester, NY, USA) and centrifuged again for 15 min at 15,000 g and 4 °C, filtered and applied to equilibrated QIAGEN Plasmid Midi Kit columns (QIAGEN, Hilden, Germany). Once all supernatant was applied, the columns were washed twice with 10 mL Buffer QC and eluted with 5 mL Buffer QF. 3.5 mL 2-propanol were added to the tubes, and the samples were spun for 15 min at 14,200 g in 2 x 5 mL reaction tubes (Eppendorf, Hamburg, Germany). The pellets were washed once with 1 mL of 70 % Ethanol thereby pooling the two pellets per sample. The pellet was then dried and resuspended in 150 μl Buffer EB. For proper solubilization, the samples were incubated over night at RT. Molar concentration of plasmid in each sample and enrichment of plasmid DNA over genomic DNA was determined by qPCR using primers

OGD241 and OGD242 (Table S5) that bind in the plasmid backbone. Samples were diluted 1/200 and a standard curve was made using a mini-prepped plasmid with its concentration determined by Qubit (Invitrogen, Waltham, MA, USA) at 0.4 ng/μl, 0.08 ng/μl, 0.016 ng/μl, 0.0032 ng/μl, 0.00064 ng/μl, 0.000128 ng/μl, and 0.0000256 ng/μl. The qPCR was performed using the 2X Sso Advanced Univeral SYBR Green Supermix (Bio-Rad Laboratories, Hercules, CA, USA) according to the manufacturers protocol.

### Sequencing library preparation

For each of the 24 samples (four batches, Input and Output sample, each in triplicate), a bit over 400 Mio reads were expected from sequencing. To prevent biases in the data, a number of plasmids of at least 10-fold higher than the number of expected reads was supposed to be used for the library preparation PCR. If the yield allowed for it, a 20-fold coverage was used. Because SP1 and SP2 Illumina primer binding sites were already present in the backbone, only one round of PCR had to be performed. To enable multiplexed sequencing, the NEBNext® Multiplex Oligos for Illumina (New England Biolabs, Ipswich, MA, USA) were used to amplify the samples (Table S11). The PCR was performed using Q5 High-Fidelity DNA Polymerase (New England Biolabs, Ipswich, MA, USA). Since more than 4 μg of genomic DNA would inhibit the PCR, the samples had to be distributed over several reactions. Five to 22 PCRs were performed for each individual sample, depending on the enrichment of plasmid over genomic DNA and the total yield. In total, 373 PCRs were performed. The PCR were run for 13 cycles, annealing at 63 °C for 30 sec and extension at 72 °C for 30 sec. Afterwards, all PCRs of one sample were pooled, mixed well, and 100 μl per sample were gel-purified with the QIAquick Gel Extraction Kit (QIAGEN, Hilden, Germany). The columns were washed three times with Buffer PE and eluted in 30 μl of Buffer EB. Sample concentrations were determined by Qubit (Invitrogen, Waltham, MA, USA), and all 24 samples were pooled at a ratio corresponding to the library size (batch 1 was bigger than batches 2, 3, and 4) and purified with AMPure XP beads (Beckman Coulter, Brea, CA, USA) using a 1:1 ratio of beads and sample as described above. For final quality control, a qPCR was performed using the KAPA Library Quantification Standards with Primer (Roche Sequencing Solutions Inc, Pleasanton, CA). The libraries were sequenced on a Illumna NovaSeq 6000 sequencer (Illumina, San Diego, CA, USA) using an entire S4 flowcell and the 35 bp single end read program pushed to 41 bp read length. 7.4 billion reads were obtained in total.

### Data processing with mutscan

Raw reads were processed using the R package *mutscan v0.2.29* (*30*). The reads consisted of a 6 bp KpnI restriction site, the 22 bp variant barcode, a 6 bp NcoI restriction site, and the 7 bp wild-type barcode. The KpnI and NcoI restriction sites were omitted, and the variant and wild-type barcodes were concatenated. All reads that did not match this structure or had an average Phred score in the barcode regions below 20 were discarded. A count table was created for the remaining reads. The reads were then split into barcode and variant barcode sequences and mapped to barcodes identified during barcode-variant association (see above). For variant barcodes, reads where the sequence did not perfectly match a 22 bp barcode with a Hamming distance of at least 2 bp to the closest barcode and were filtered out (826,286 barcode-variant pairs). Read counts were then aggregated according to the variant and wild-type partner amino acid sequences. The four batches were then normalized using the JUN mutant data present in all four batches by least square regression. To keep read counts per bZIP balanced, only the JUN reads originating from batch 2, which has more read counts, were kept in the final data table. Binding scores were calculated as Log fold changes (logFC) using the *mutscan* function calculateRelativeFC() with the method “limma” (*31*). Most of the pairs involving variants of ATF7 were absent from the dataset, including all wild-type interaction, thus preventing computing mutational effects relative to wild-type for ATF7. All variants of ATF7 were thus filtered out. Mutation effects were calculated by using a different linear model in limma in order to calculate a logFC between the variant counts and the corresponding wild-type counts. Low coverage data points, i.e. those with a read counts less or equal to 10 in any input replicate or 0 read in any output replicate, were filtered out except when specified otherwise (e.g. for thermodynamic model fitting). Mutation with significant effects relative to wild-type were identified as those with a mutation effect below -2 or above 2, for negative and positive effects respectively, at a false discovery rate of 5% (Benjamini-Hochberg).

### Specificitiy rewiring mutations

Mutations that completely change the binding profile of a bZIP were identified as those for which the wild-type variant had at least one binding partner (binding scores > 1 and at least 15 significantly detrimental mutations, i.e. mutations with a mutation effect < -2 and FDR < 5%), the mutation had a significant positive effect with at least one partner not bound by the wild-type variant (mutation effect > 2, FDR < 5% and wild-type binding score < 1) and none of the partners bound by the wild-type variant were also bound by the mutant (wild-type binding score > 1 and mutant binding score > 1).

### Global thermodynamic model

We used MoCHI (https://github.com/lehner-lab/MoCHI) (*21*, *33*) to fit a single global two-state thermodynamic model (Fig. 2B) to the whole dataset, including all variants not expected according to the experimental design and the oligo pool synthesis but identified as present in the library during the barcode-variant association. These include notably double mutants. Expected single amino acid variants with low read counts were also included because more data improves model performance even if the additional data is noisier. Variants corresponding to mutants of ATF7 were filtered out given the low number of variants present in the library. Variants with premature stop codon were also filtered out. The reference interaction chosen was JDP2-DDIT3 because it is a strong and reliable wild-type interaction. Since MoCHI takes a single amino acid sequence as input, the wild-type partner identity was dummy encoded in a 53 amino acid string where A53 represents the DDIT3 reference partner and each other partner is encoded as a substitution towards cysteine at each of the 53 positions in this sequence. The dummy sequence was then concatenated to the variant amino acid sequence. MoCHI was then ran with default parameters. All non-expected variants and low read counts variants were then filtered out to assess model performance.

### Individual thermodynamic models

For each dataset corresponding to all variants of one bZIP paired with the 54 wild-type partners, a different model was fitted using MoCHI in the exact same way as for the global thermodynamic model.

### Deep learning model training

#### Dataset preparation

Variants with a premature stop codon were removed and sequences shorter than 62 residues were padded with stop codon symbols at the end. Multiple mutants present in the dataset (not designed but originating from mutations during library construction and validated during barcode-variant association) were removed. Data points with input count reads less than 10 were removed. Ten percent of the data were held out for testing, 10% for validation and the rest was used for training. Splitting was done randomly based on different unique features, i.e. random data points (“random”), based on substitution to specific amino acid at specific positions in specific bZIPs with all wild-type partners (“mutation ID”), based on substitution to specific amino acid at specific positions in any bZIPs with all wild-type partners (“amino acid”), all substitutions at specific positions with all wild-type partners (“position”), or any substitution at any position in specific bZIPs with all wild-type partners (“bZIP”). After splitting, the binding scores were centered and normalized by subtracting the mean of the training data and division by the standard deviation.

In case of CNN training, the bait and prey sequences were one-hot encoded. For ESM training, the bait and prey sequences were concatenated with a 50 aa glycine linker in the middle. Tokenization was carried out with the AutoTokenizer class from the transformers library (4.37.2) using the ESM-specific tokenizer supplied with the pretrained model.

#### CNN training

Neural network training was carried out using the PyTorch (2.1.1) and PyTorch Lightning (2.1.1) frameworks. The MSE loss function was minimized using the Adam optimizer and a learning rate of 3.15*10-4. L1 regularization was applied to the loss function using a factor of 5*10-7. Training was performed on Nvidia V100-SXM2-32GB, A40, or A100-SXM4-80GB with 10x cross-validation (90% training and 10% validation). Training was evaluated based on validation loss over 15 epochs and the epoch checkpoint with the lowest validation loss was used for further analysis. The average R^2^ and standard deviation across the 10 folds was reported (Fig. 5C).

#### CNN hyperparameter search

The optuna libray was used to perform hyperparameter search on the following parameters: learning rate, L1 weight, batch size, convolutional kernel size, convolutional filter number, number of convolutional layers.

#### CNN architecture

The final architecture comprised a 1D convolutional layer with 140 filters and a kernel size of 15 that receives one-hot encoded bait and prey sequences separately (with bait and prey sequences sharing the same filters), followed by concatenation of the two outputs along the sequence axis and inversion of sequence and filter channel axes. The transformed features are passed to a block of three 1D convolutional layers with 140 filters and a kernel size of 15. Each of the three convolutional layers is followed by batch normalization and ReLU (Rectified Linear Unit) activation. The output is flattened to one dimension and a dropout of 20% is applied before passing it to a linear layer with the same input and output size of 140 followed by GELU (Gaussian Error Linear Unit) activation. Another dropout of 20% is applied and the features are reduced to a size of 1 using a linear layer.

#### ESM fine-tuning

Fine-tuning of the ESM 15 billion parameter model was carried out using the Trainer class from the transformers library (4.37.2) in combination with the Low-Rank Adaption of Large Language Models (LoRA) method using the PEFT (Parameter-Efficient Fine-Tuning) library (0.9.0). To fine-tune a subset of parameters from the ‘key’, ‘value’, ‘query’, ‘output.dense’, ‘intermediate.dense’ layers of the base model over one epoch, a LoRA rank of 2, LoRA alpha of 2, LoRA dropout of 0.01, maximum gradient normalization of 0.5, batch size of 32, and weight decay of 0.3 was used. Simultaneously, a classification head was trained for a regression task using the EsmForSequenceClassification class from the transformers library. The default classification head architecture was used, which consists of a dropout function (10%), a linear layer (input and output size of 5120), a tanh activation function, another dropout function (10%) and a final linear output layer (input size 5120, output size 1). The MSE loss was minimized with Adam optimizer and a cosine learning rate scheduler starting at 3.7*10-4. Quantization was performed with BitsAndBytes (0.43.0) using 4-bit precision (int4) for model weights and bfloat16 for activations. Fine-tuning was evaluated based on validation loss over 1 epoch and the epoch checkpoint with the lowest validation loss was used for further analysis. Training was performed on Nvidia A40, or A100-SXM4-80GB. Due to high resources requirements, training was performed on a single fold (80% training, 10% test, 10% validation). For comparison, training of the CNN with the amino acid split was performed on the same fold.

#### Analysis of results

Training was tracked with Weights and Biases using the coefficient of determination as the main metric.

#### Synthetic bZIPs design

A genetic algorithm was used in combination with a pretrained CNN for scoring of binding scores (model architecture and training described in detail below).

#### CNN training

A preliminary version of the CNN (compared to those described above) was used for the synthetic bZIP design. A similar model architecture was used, where the inputs (one-hot encoded sequences) are concatenated along the feature dimension followed by swapping of the sequence and feature dimensions. This input is passed to two sequential blocks each consisting of two residual 1D convolutional layers (7 kernels and 124 filter channels). All convolutional layers are followed by batch normalization and ReLU activation. The output is flattened by calculating the mean along the sequence dimension and passed to a linear layer block consisting of a dropout function (20%), a linear layer (124 neurons), a GELU activation function, another dropout function (20%), and a linear layer with a single neuron.

The network was trained for 10 epochs with a dataset split by the “amino acid” feature (with all data points available, including multiple mutants) and the following hyperparameters: learning rate: 1.11*10-3, weight decay: 0, L1 regularization factor 1.05*10-4.

#### Genetic algorithm

The genetic algorithm was run over 100 generations, each comprising a tournament step (selection, mutation, crossover) and subsequent evaluation. In the selection step, 16 “contestant” sequences were chosen from the given population (initially the 54 bZIP wild-type sequences), scored with the fitness function (described below) and the two top-ranked sequences were selected as parents. These two parent sequences were subjected to crossover at a random position within the sequence. This step was repeated, swapping the first and second parent sequence, and keeping the same crossover position. In the following mutation step, the resulting two child sequences were mutated at all positions between residue 28 and 62 (the zipper domain) with every amino acid and added to the new population. The tournament step was repeated another time, overall yielding two new sequence populations. The score from each of the two tournaments was compared to the overall best seen binding score, and if it was higher, the first parent sequence (obtained from the selection step) of the respective tournament was selected as the new best sequence. For the next iteration, the two populations were combined.

After completing all generations, the obtained best sequence was scored against the 54 bZIP sequences using the CNN model. These steps (genetic algorithm and scoring) were repeated over 10 iterations.

#### Fitness function

The given sequence is scored against the target sequence, against itself and against all other bZIP sequences using the CNN. To maximize target binding, while minimizing homodimerization and unspecific binding, the “self” score and average “other” score are subtracted from the “target” score.

#### Small-scale bindingPCA to validate the predicted strong binders

The strong binder library was ordered as oligo library from Twist Bioscience (San Francisco, CA, USA). Both zipper domain and DBD were encoded on the oligo together with a pre-designed barcode (Table S12) to identify each bZIP. The oligo library was amplified with Q5 High-Fidelity DNA Polymerase (New England Biolabs, Ipswich, MA, USA) in 15 cycles using an annealing temperature of 60 °C and primers oDL00763 and oDL01077. The amplicons were purified from a 1 % Agarose gel using the QIAquick Gel Extraction Kit (QIAGEN, Hilden, Germany). They were then digested with KpnI-HF and BamHI-HF at 37 °C for two hours and cloned into a random backbone of the intermediate wild-type prey library which had been digested with the same enzymes using NEB T4 ligase (New England Biolabs, Ipswich, MA, USA) according to the manufacturers protocol. The ligation setup was then transformed into NEB 10-beta Electrocompetent *E.coli* (New England Biolabs, Ipswich, MA, USA) according to the manufacturer’s protocol. The tranformants were harvested as described above and purified using the QIAGEN Plasmid Plus Midi Kit (QIAGEN, Hilden, Germany). In a second step, the purified library and one batch of the heterodimer library were digested with NcoI-HF and SpeI-HF. The backbone stemming from the first cloning step and the insert stemming from the heterodimer library were cloned together and transformed into electrocompetent bacteria the same way as above. A bCA of the library was then performed in the same way as described above but in four replicates and scaled down to 200 mL cultures. The samples were purified and prepared for sequencing as described above. Sequencing was done on an Illumina NovaSeq 6000 Sequencing System (Illumina, San Diego, CA). Reads were processed and transformed into binding scores with *mutscan v0.2.37* (*30*).

---

**Fig. S1.**
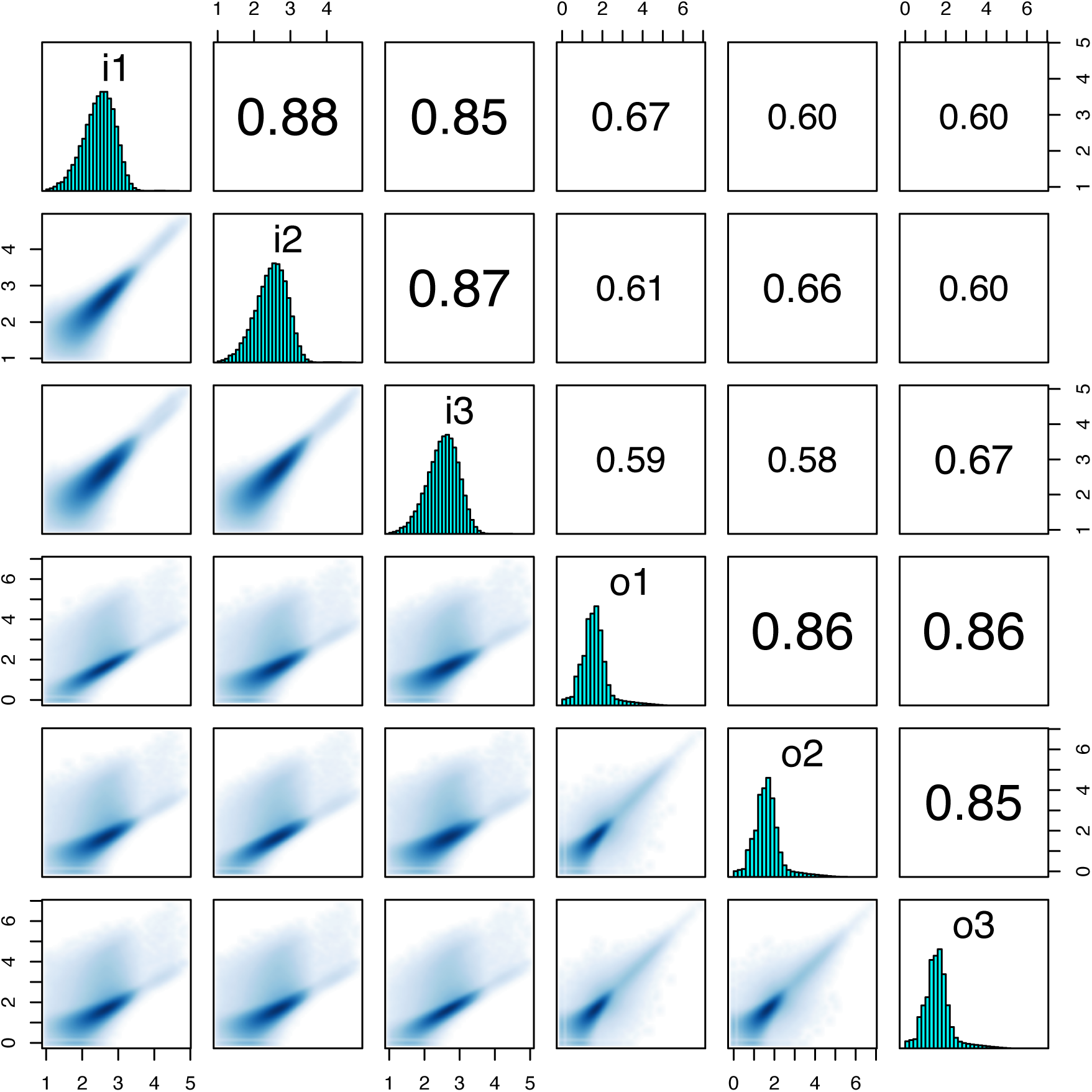
Correlation between input and output read count correlations across the three biological replicates. Pearson correlation scores are shown in the upper right boxes, count distribution are shown in the diagonal boxes, and scatter plots are shown in the lower left boxes. i1-3, input samples for replicates 1 to 3; o1-3, output samples for replicates 1 to 3.

**Fig. S2.**
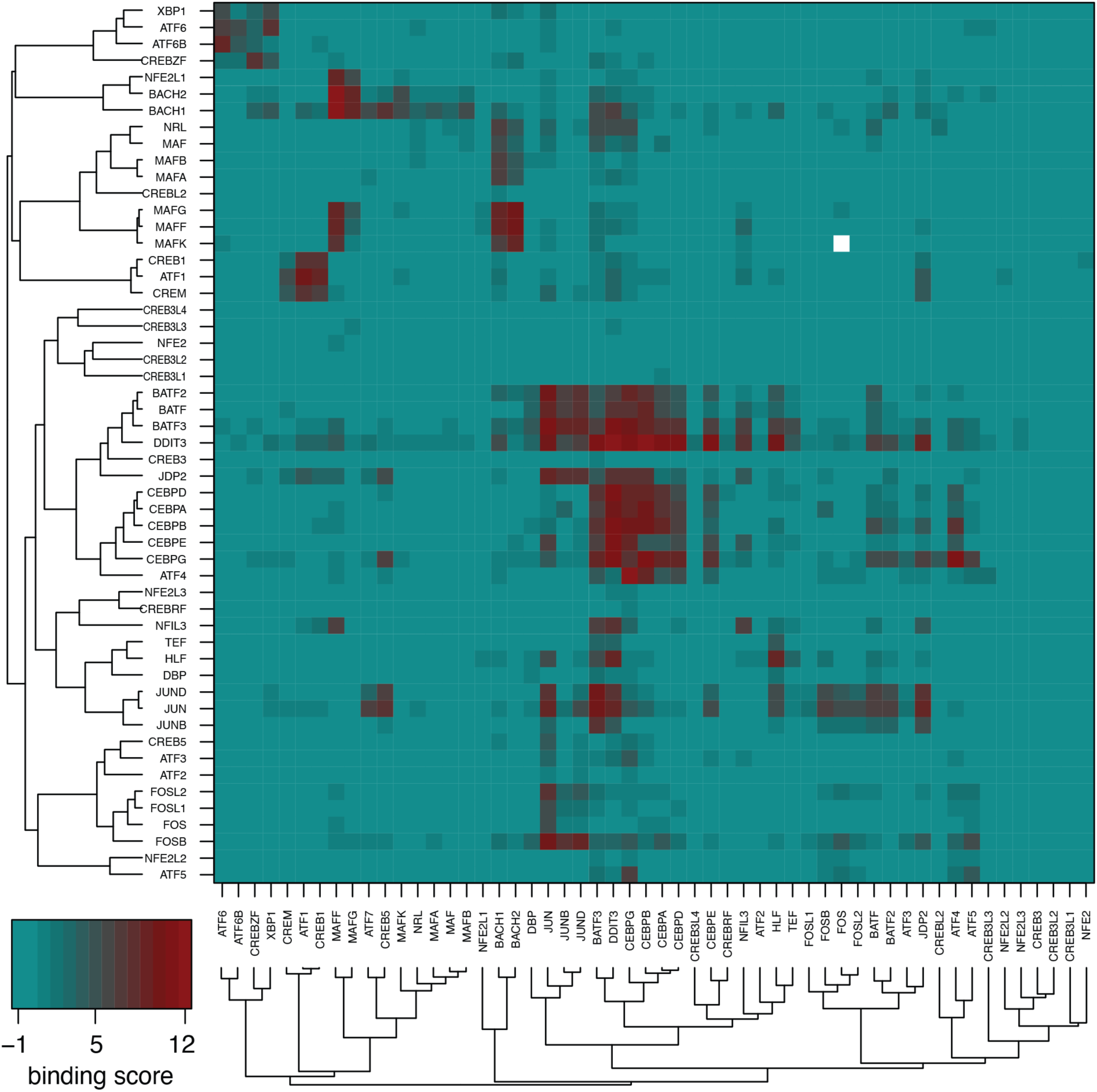
Wild-type interaction scores recapitulate the modular organization of the bZIP network. Heatmap of binding scores for each wild-type interaction. Dendrograms represent hierarchical clustering based on the Pearson correlation coefficients across interaction profiles. White represents missing data

**Fig. S3.**
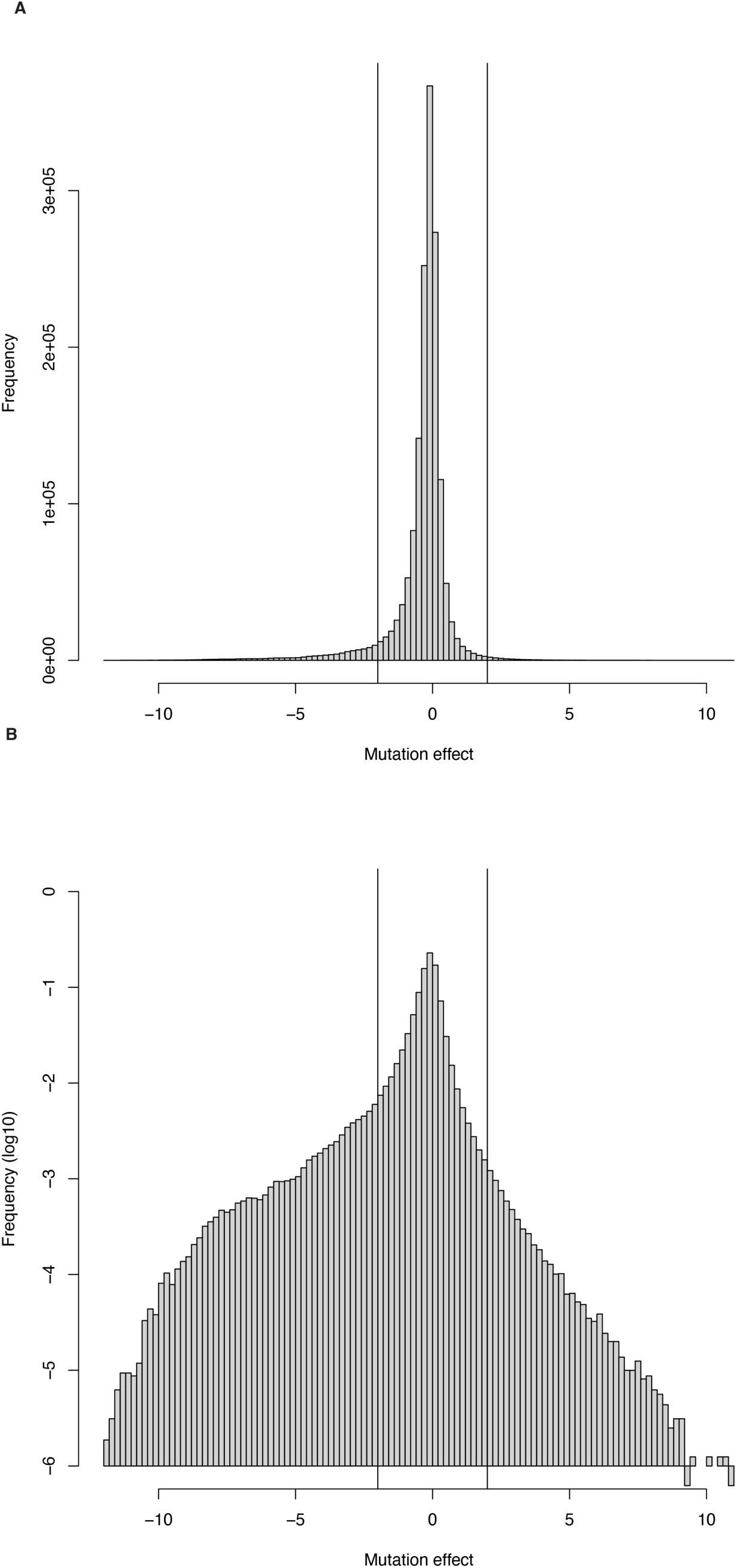
Distribution of mutation effects (i.e. binding scores relative to the corresponding wild-type interaction) on a normal scale (**A**) or a log10 scale (**B**) to highlight the difference between strongly negative and positive mutation effects.

**Fig. S4.**
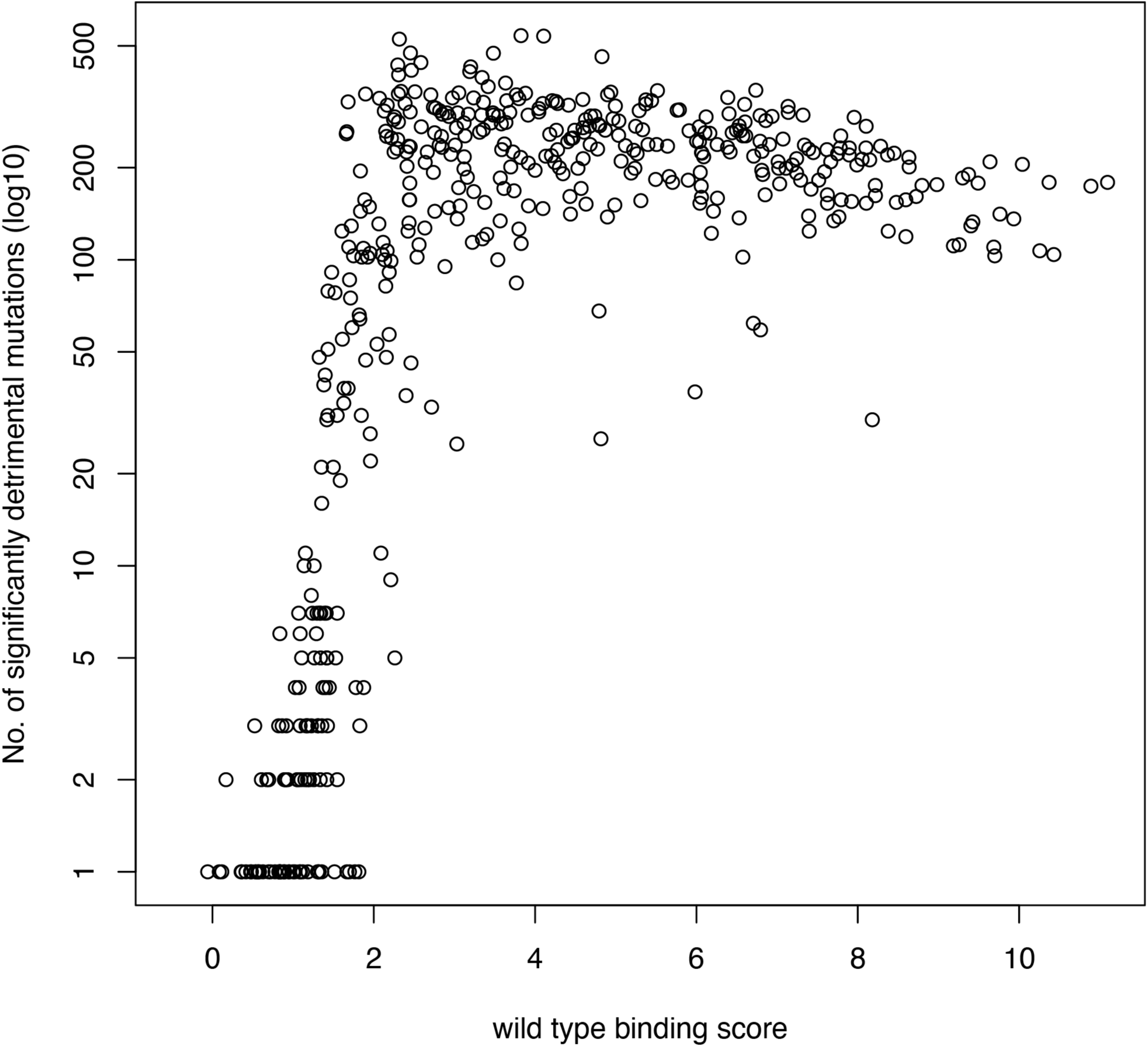
Relationship between the wild-type binding score and number of significantly negative mutation effects (mutation effect < -2 and FDR < 5%) for each bZIP pair, showing that mutations cannot detectably decrease interactions that are already weak or undetectable.

**Fig. S5.**
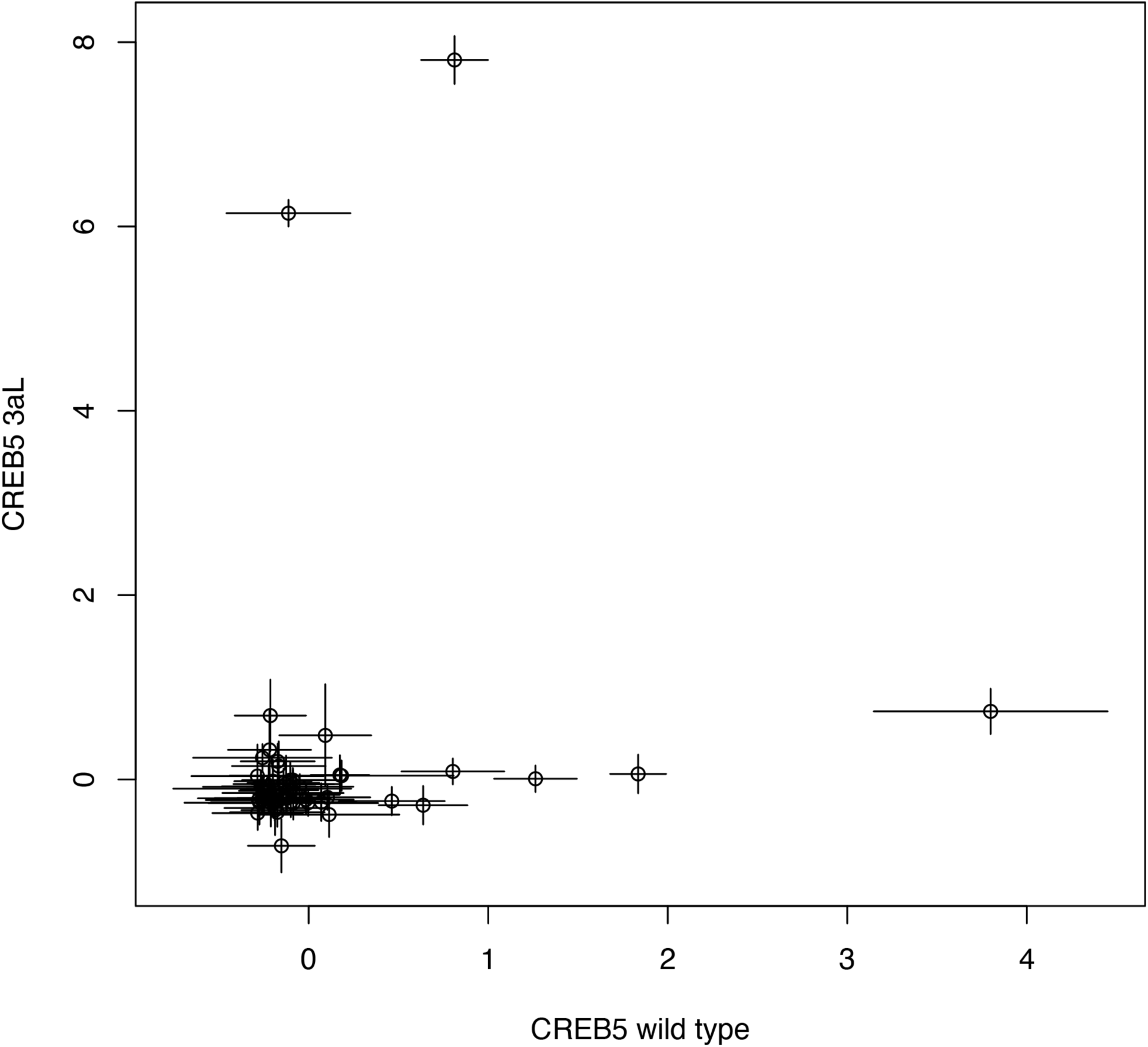
Comparison of binding scores profiles between wild-type CREB5 and substitution towards leucine at position *a* in heptad 3. Error bars represent the precision-weighted standard error across three replicate measurements.

**Fig. S6.**
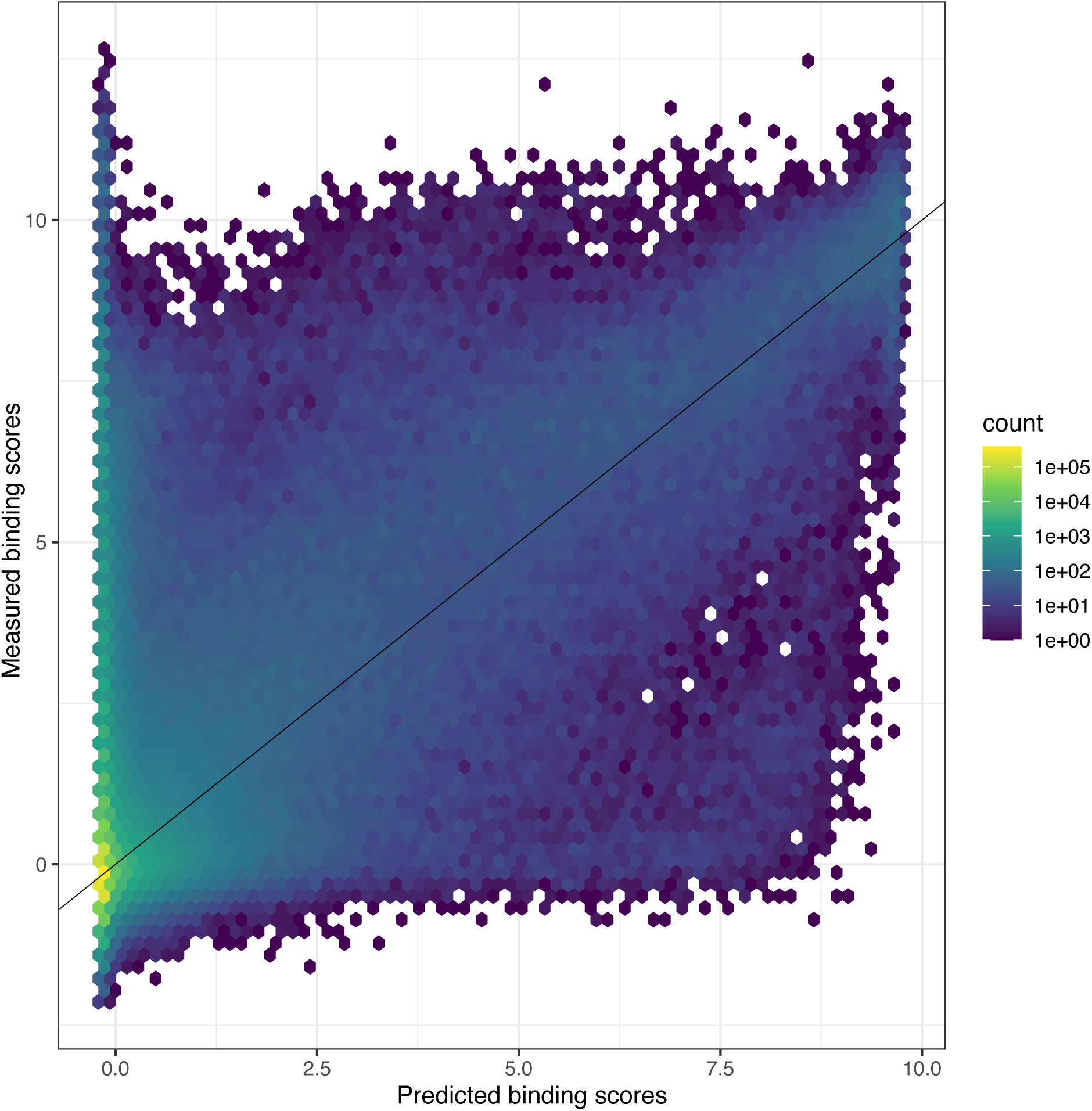
Comparison of measured and predicted binding scores for the global thermodynamic model.

**Fig. S7.**
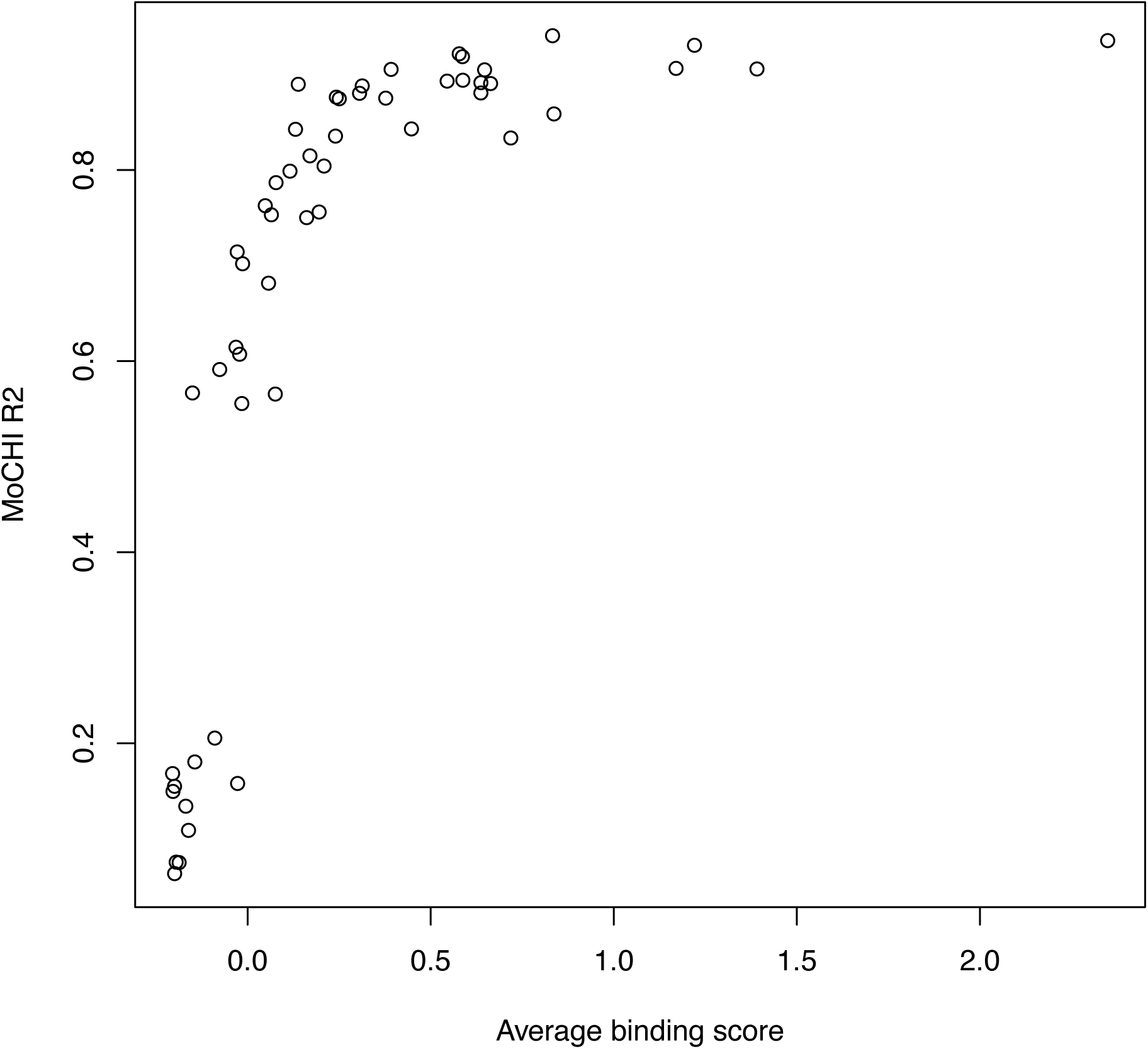
Comparison average binding score and indivdual thermodynamic model performance for each of the 53 bZIPs (ATF7 has been excluded due to large amount of missing data). The relationship shows that low performance is due to biases in the distribution of binding scores, since low binding scores are inherently noisier.

**Fig. S8.**
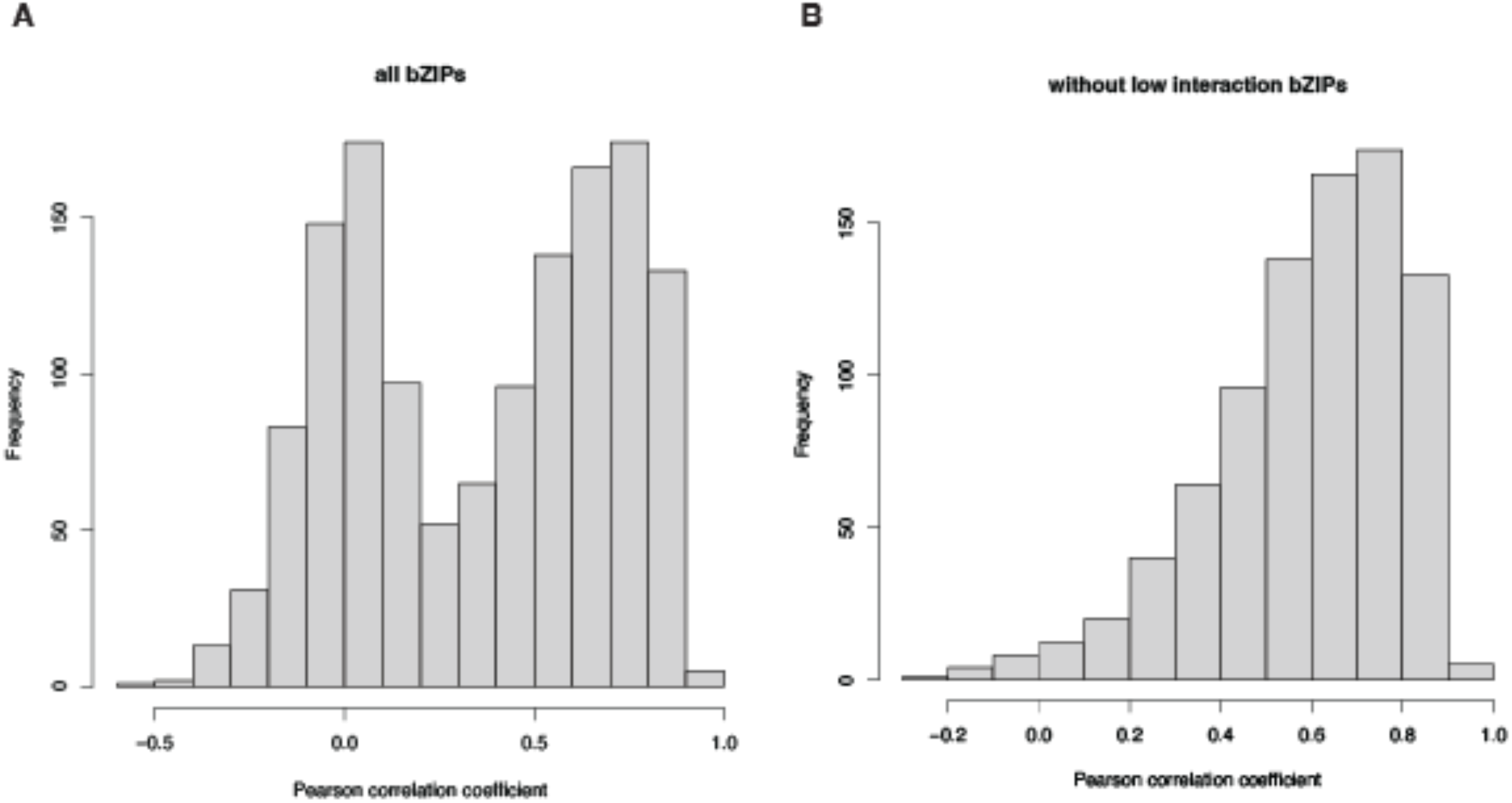
Distribution of Pearson correlation coefficients between fitted ΔΔG_mut values for the same substitution across all bZIP pairs, including (**A**) or excluding (**B**) information-poor bZIPs (individual thermodynamic model R^2^ < 0.4).

**Fig. S9.**
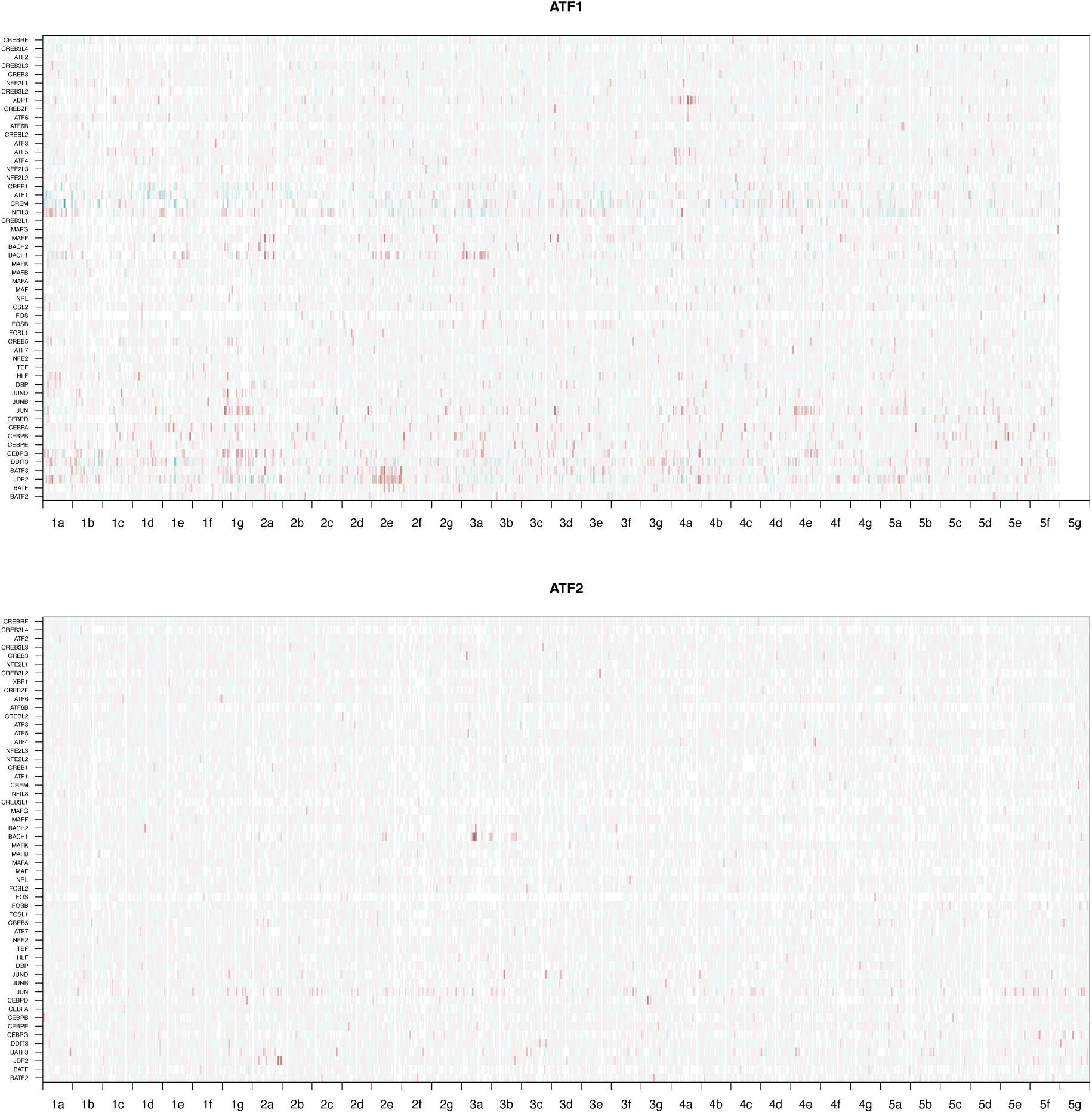
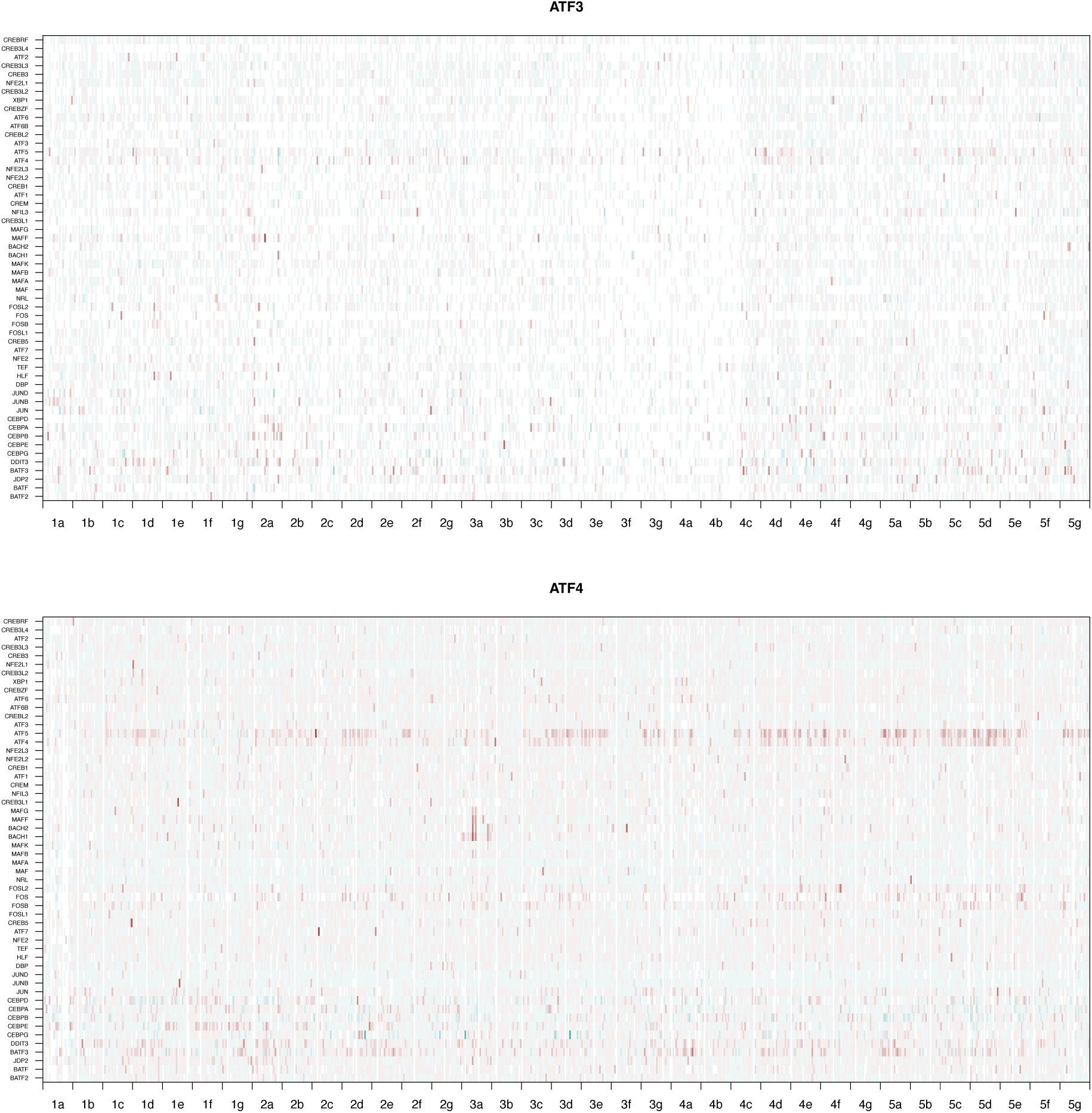
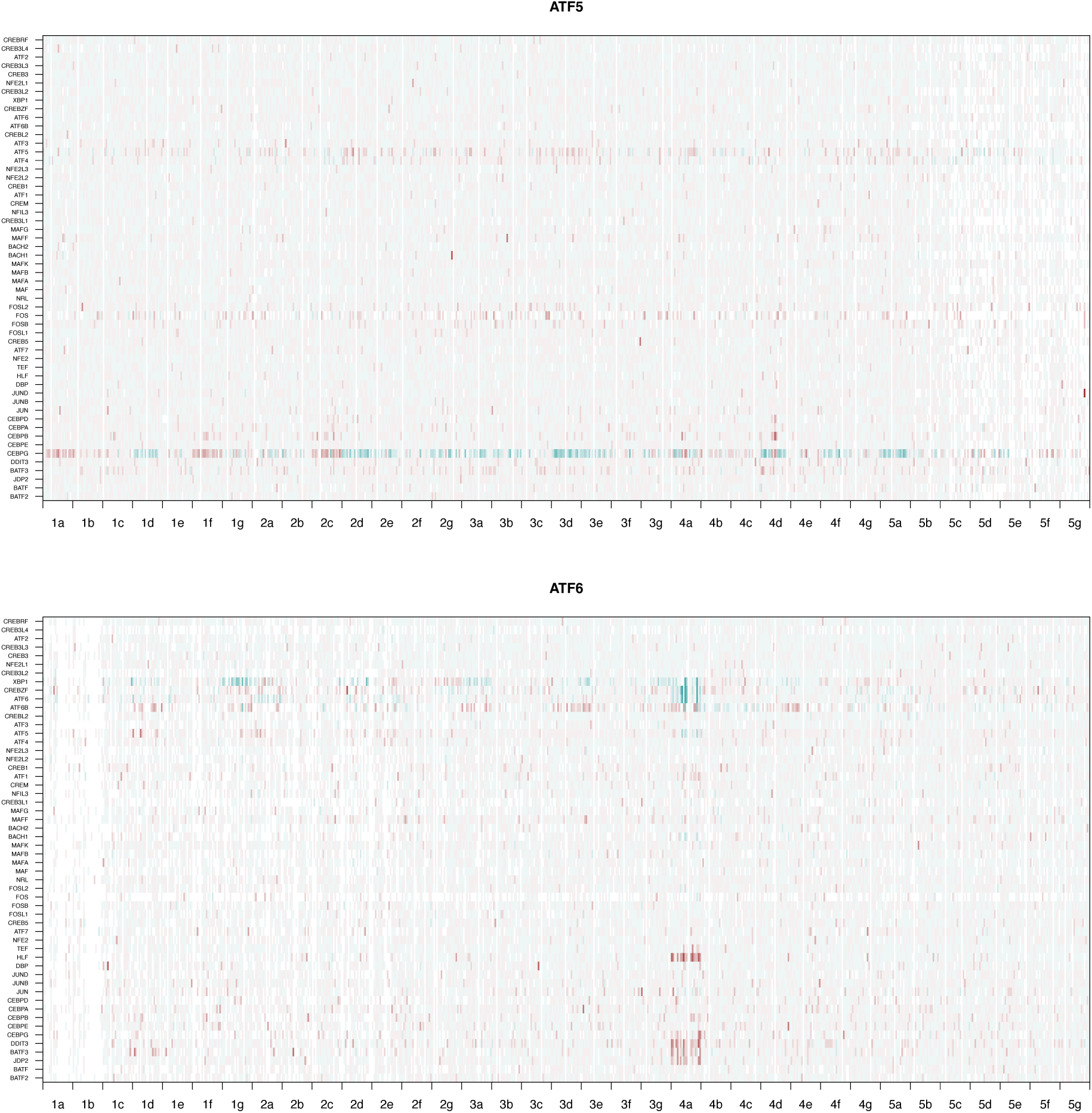
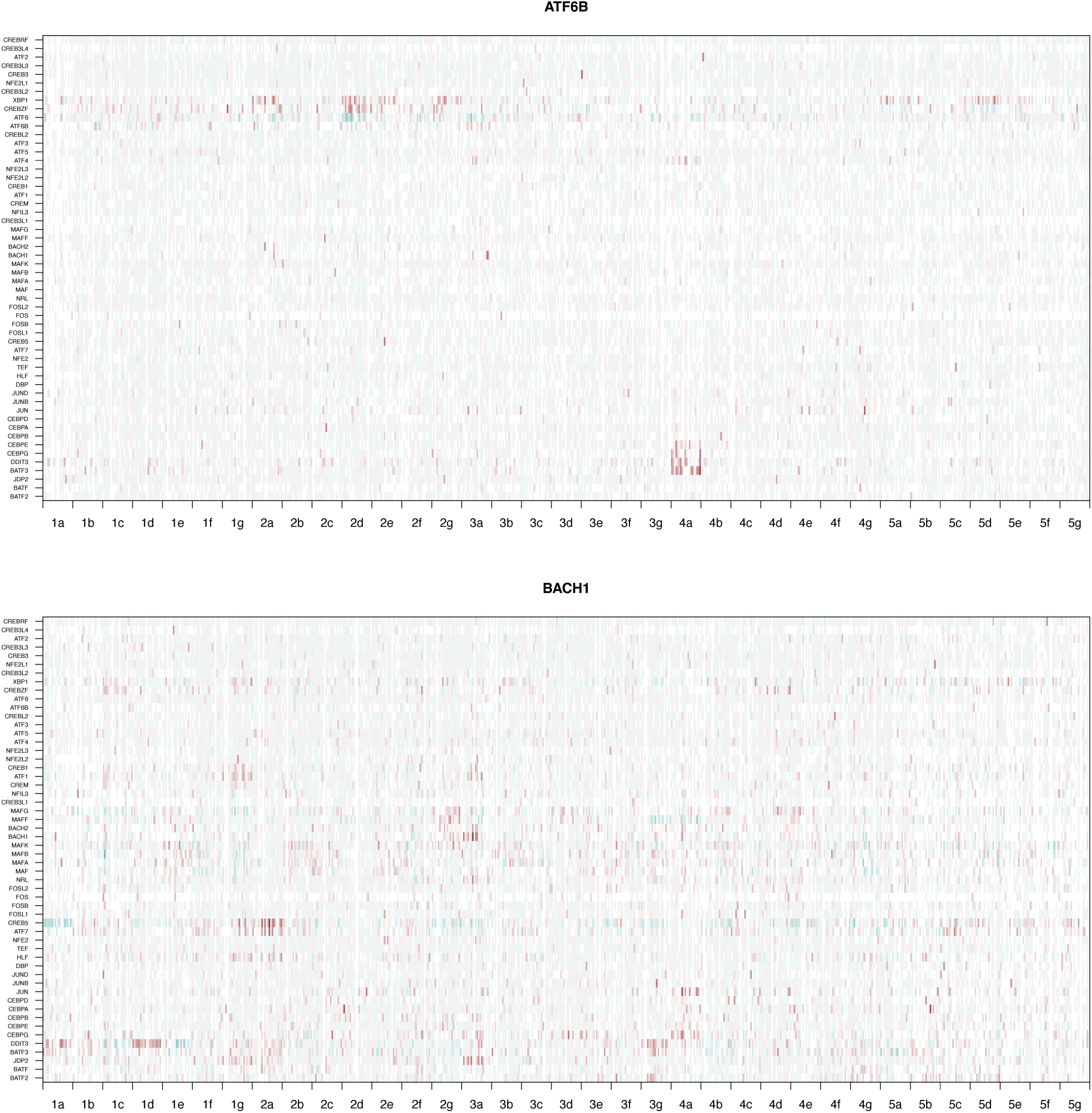
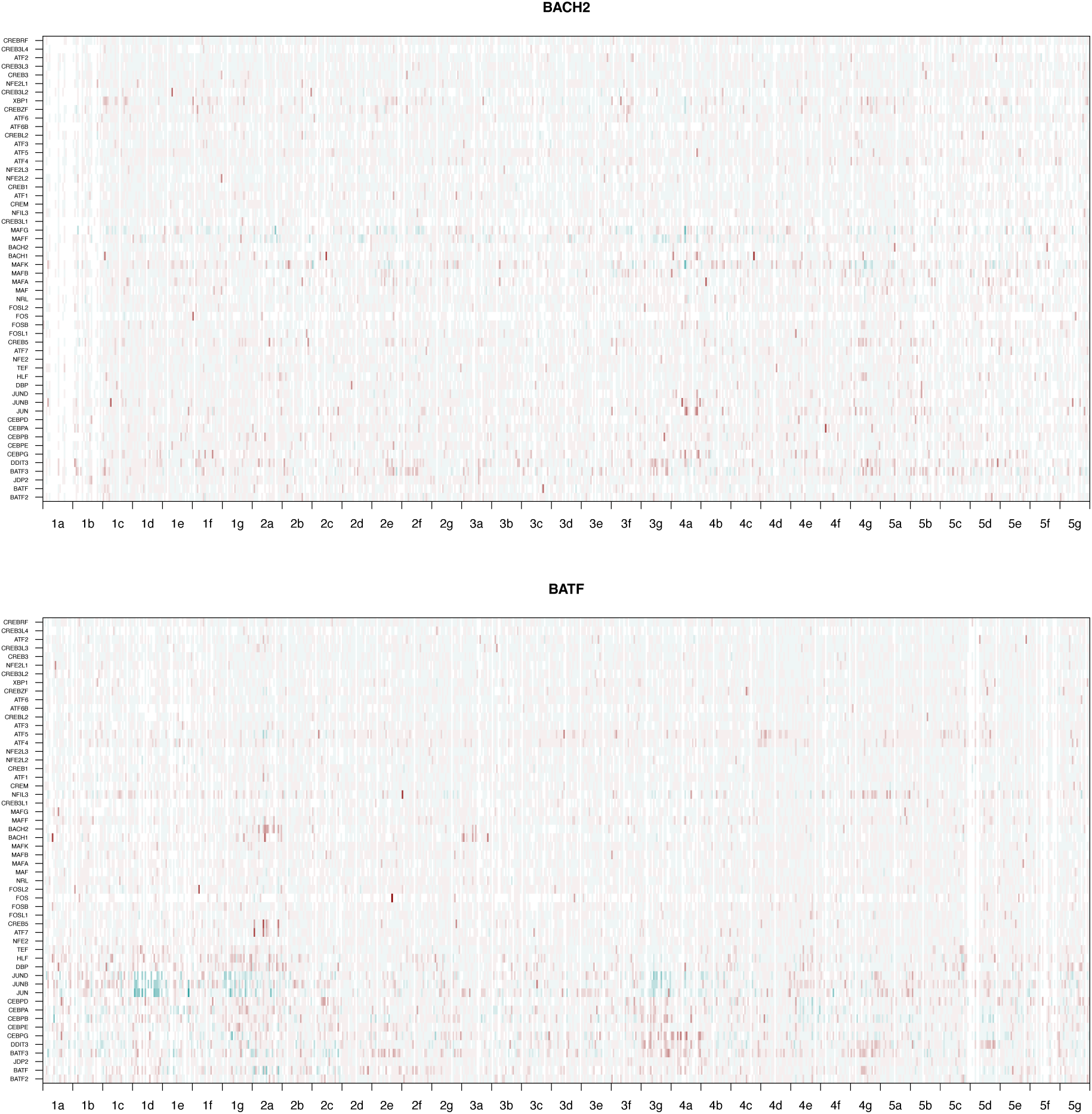
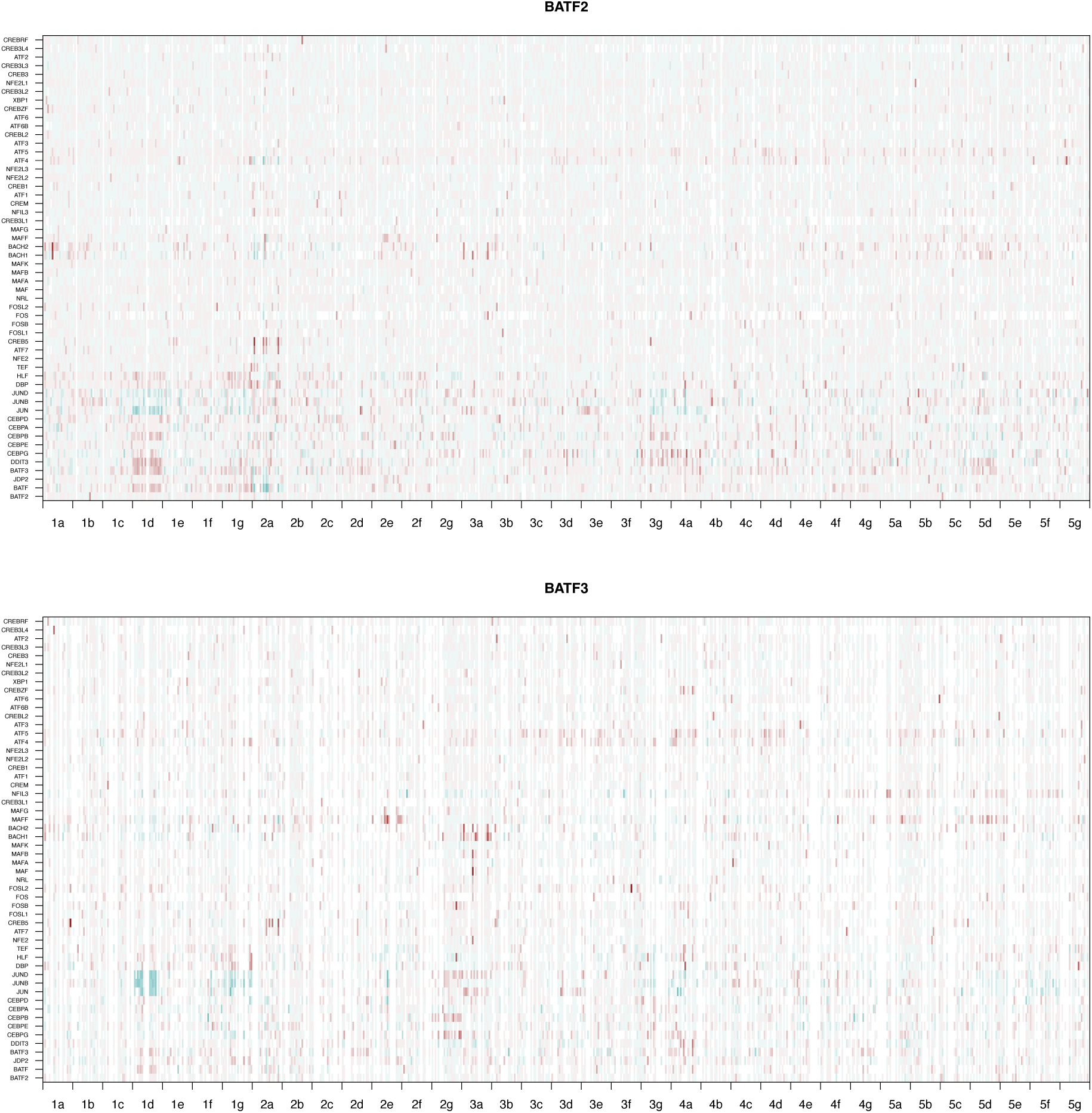
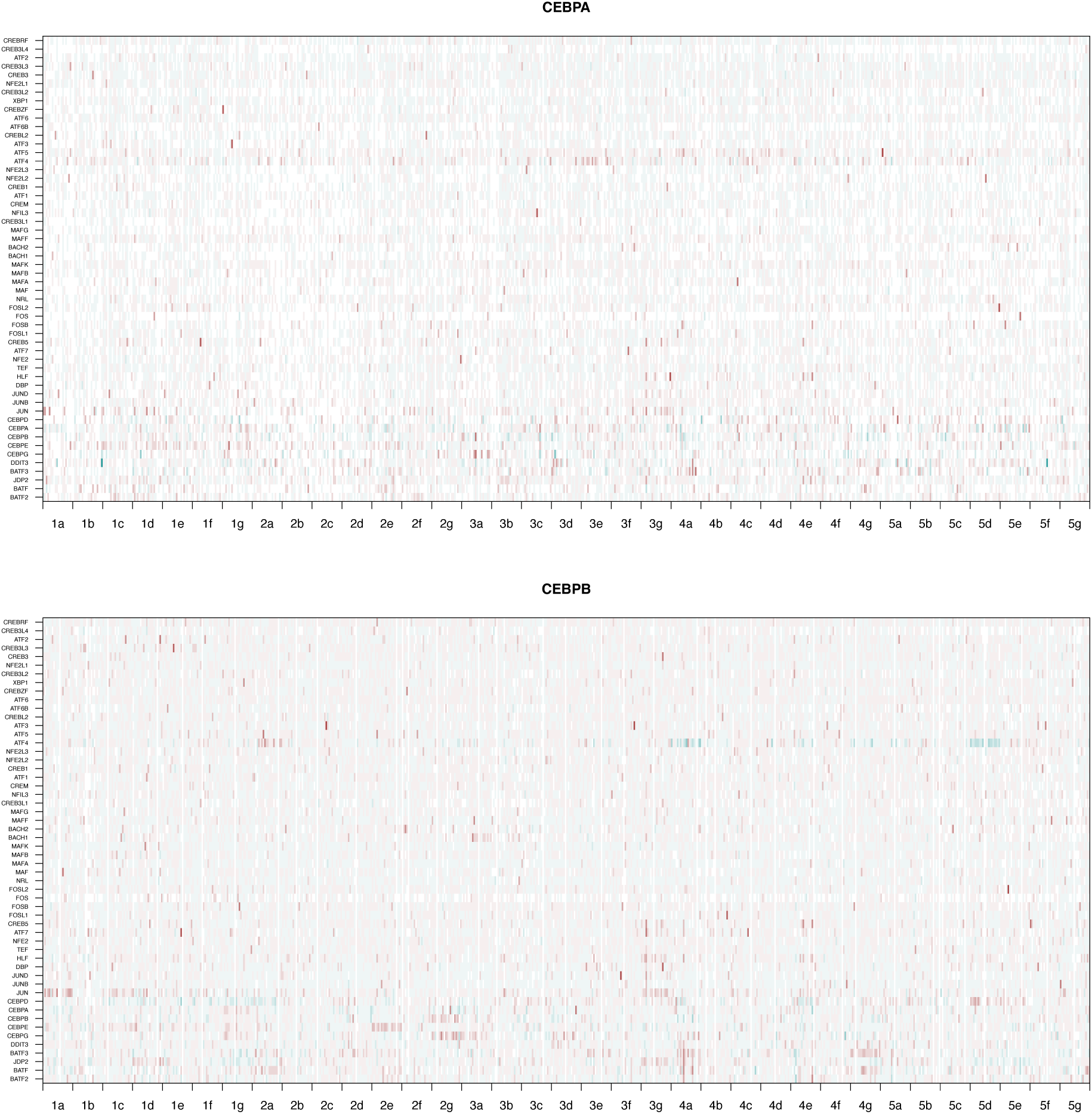
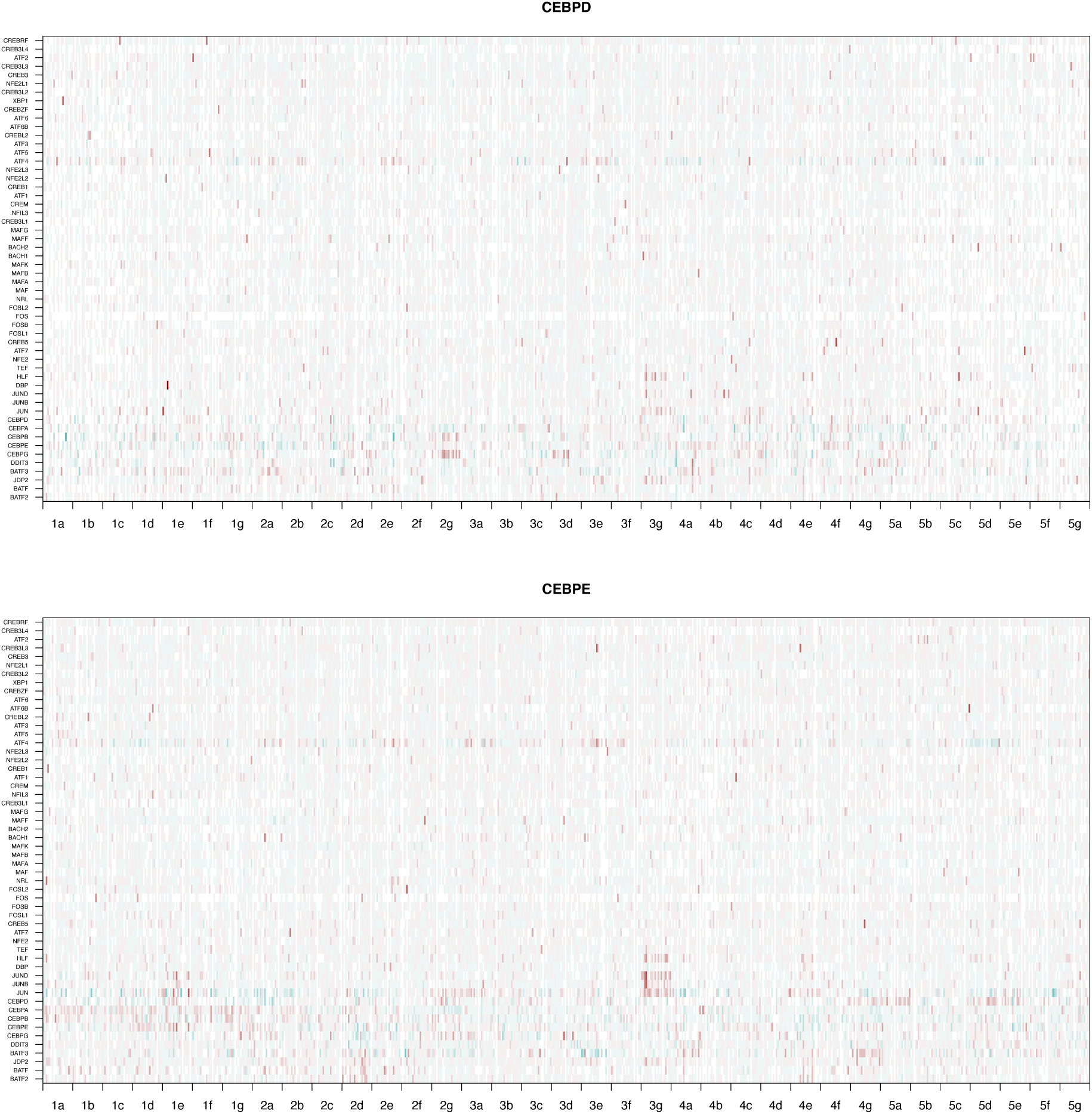
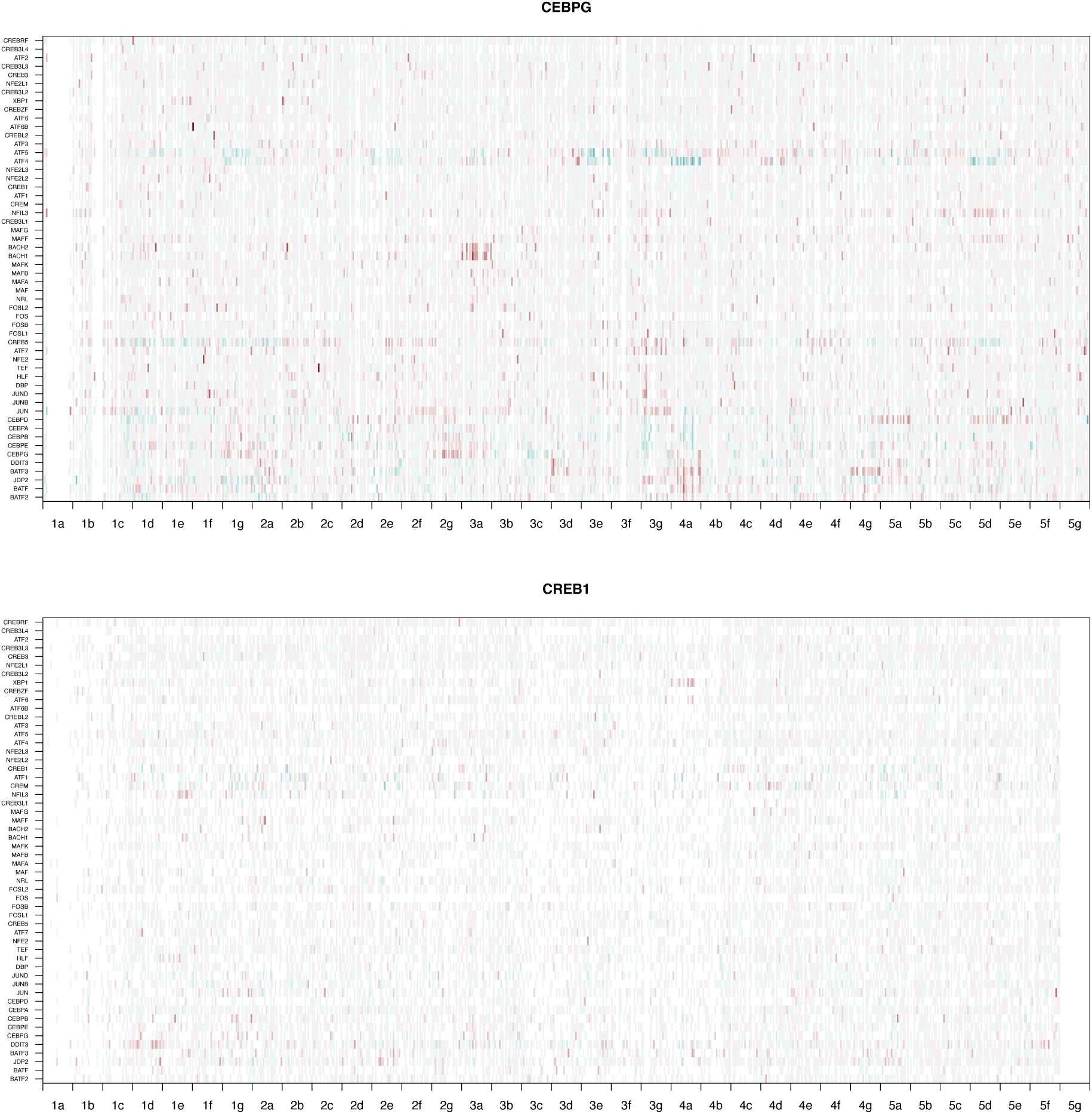
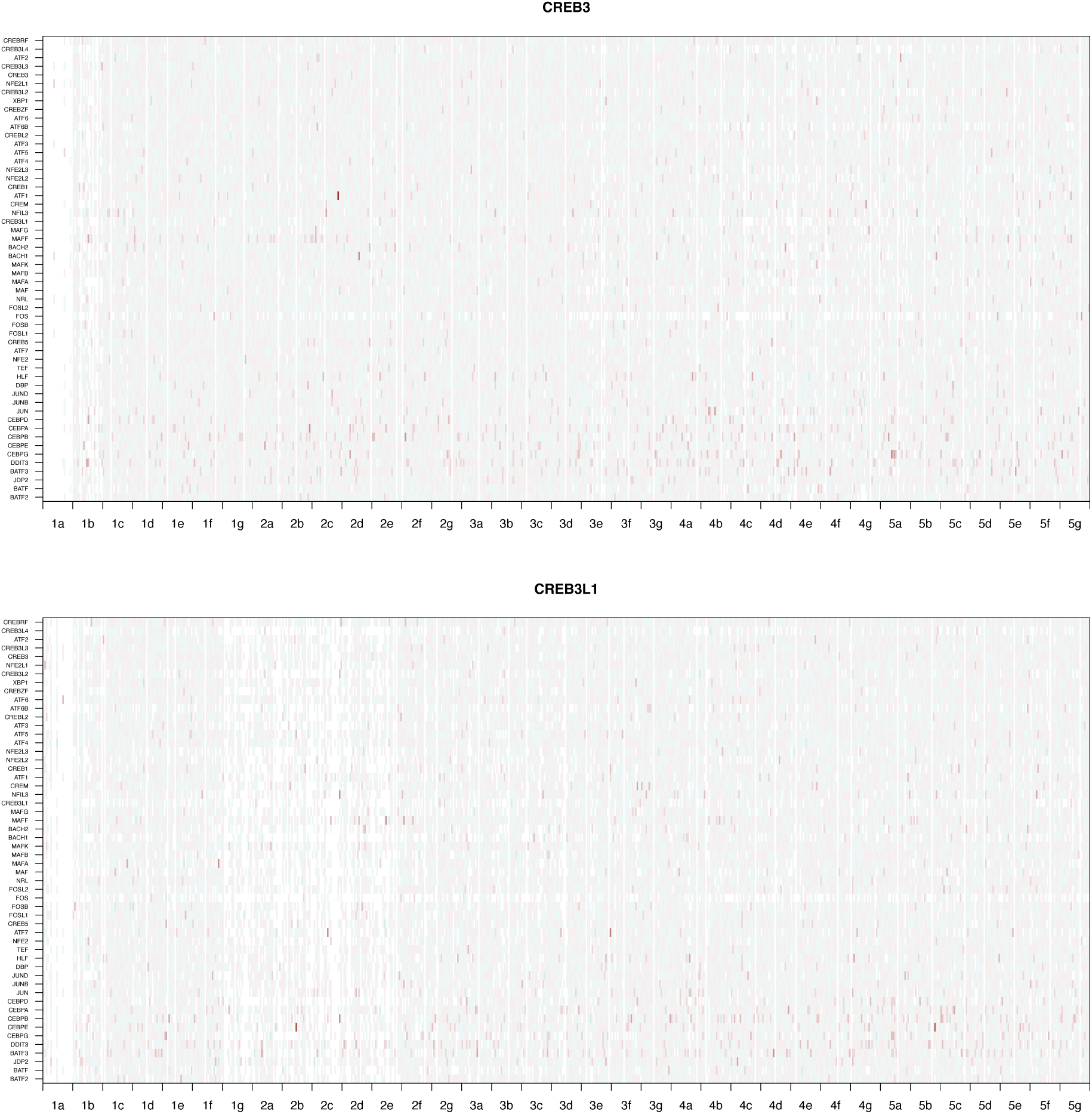
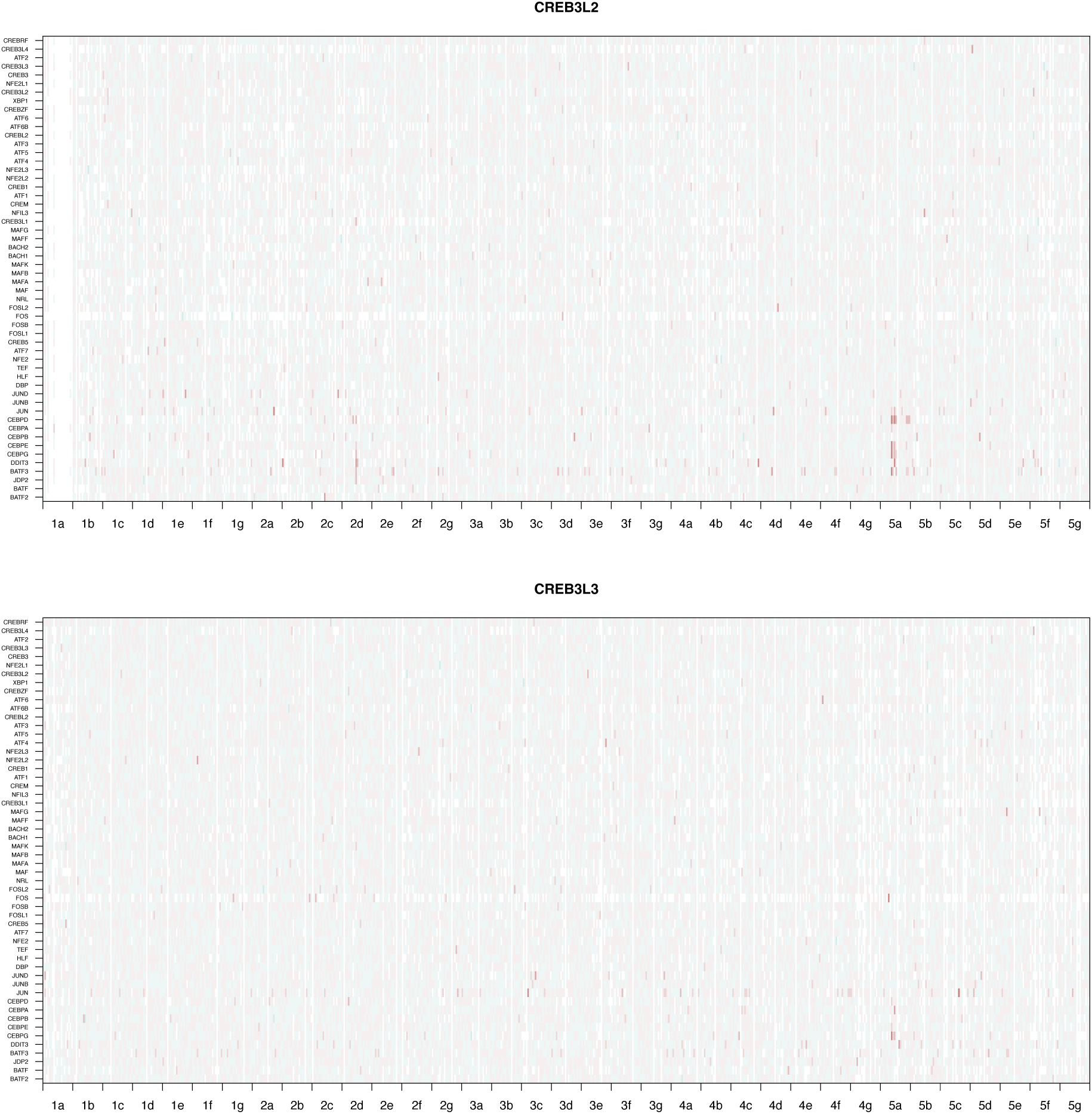
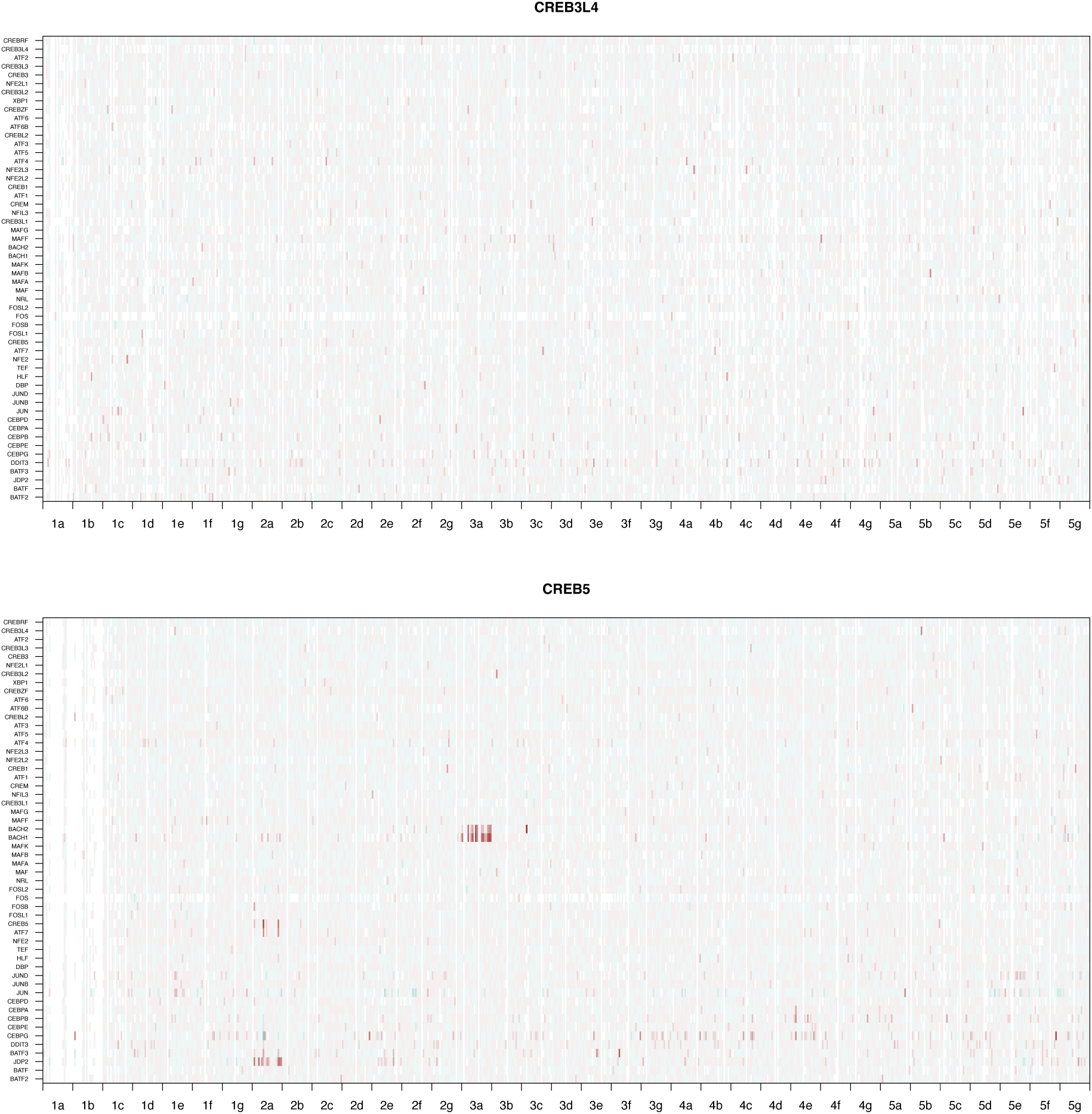
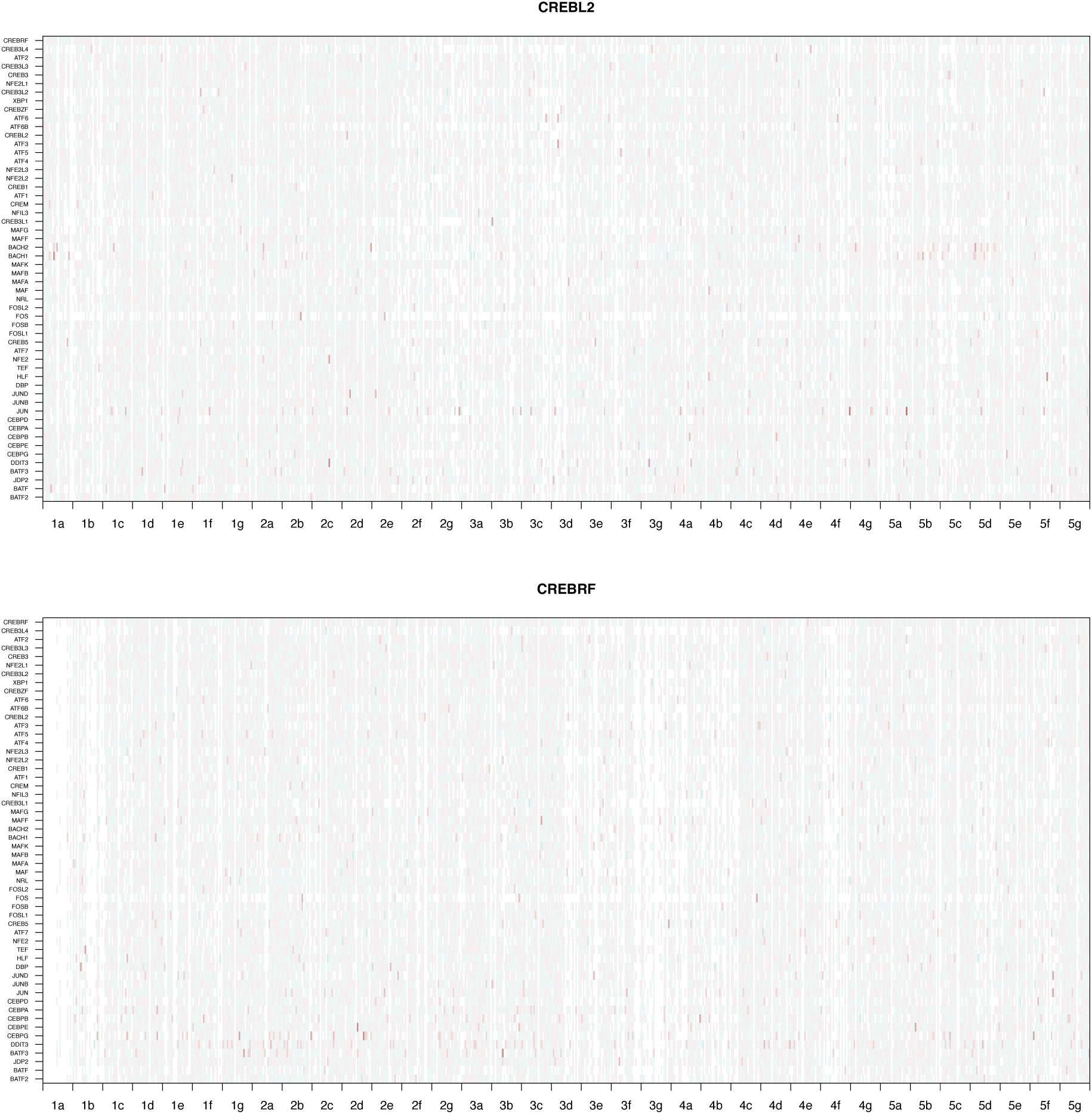
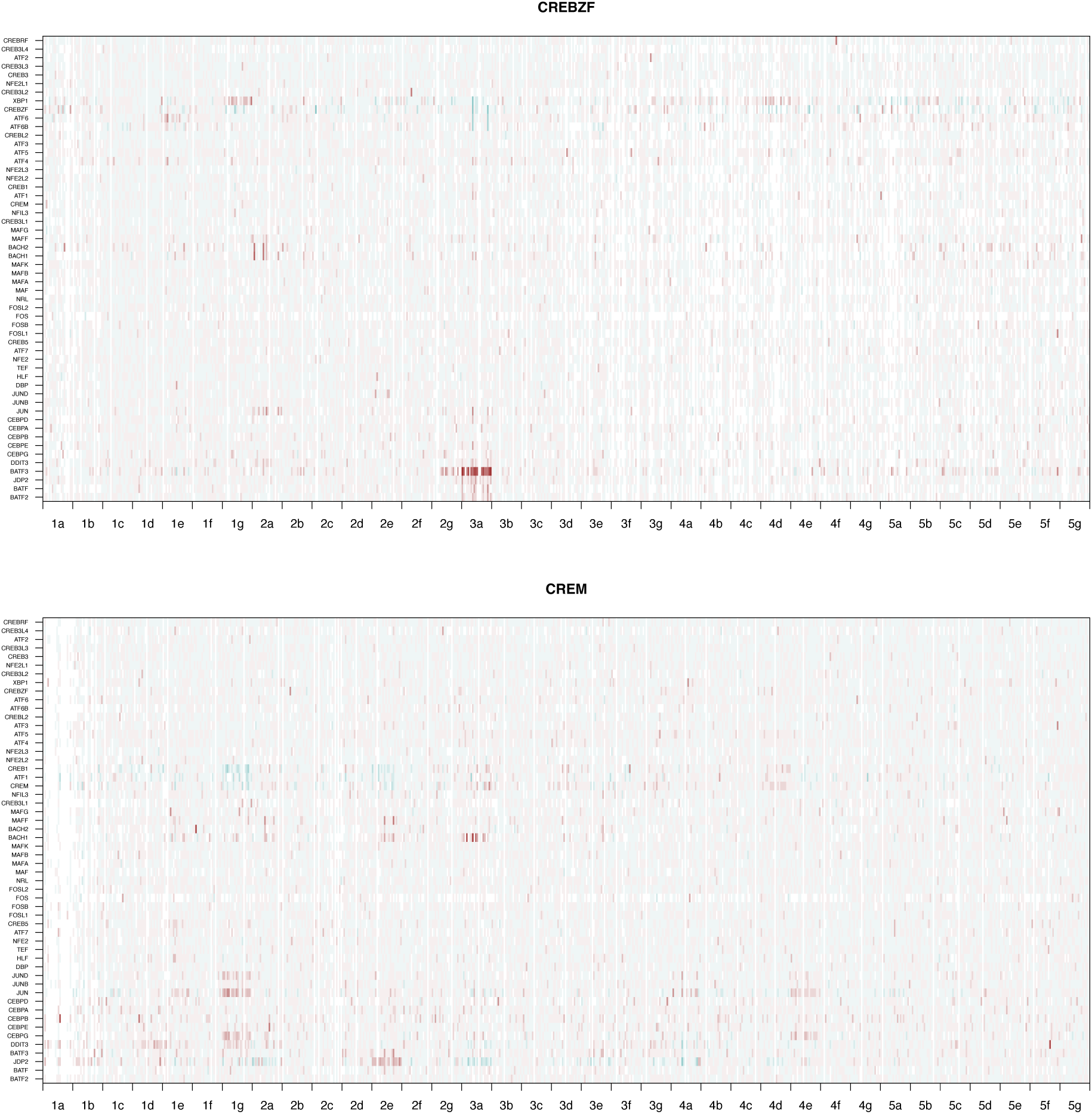
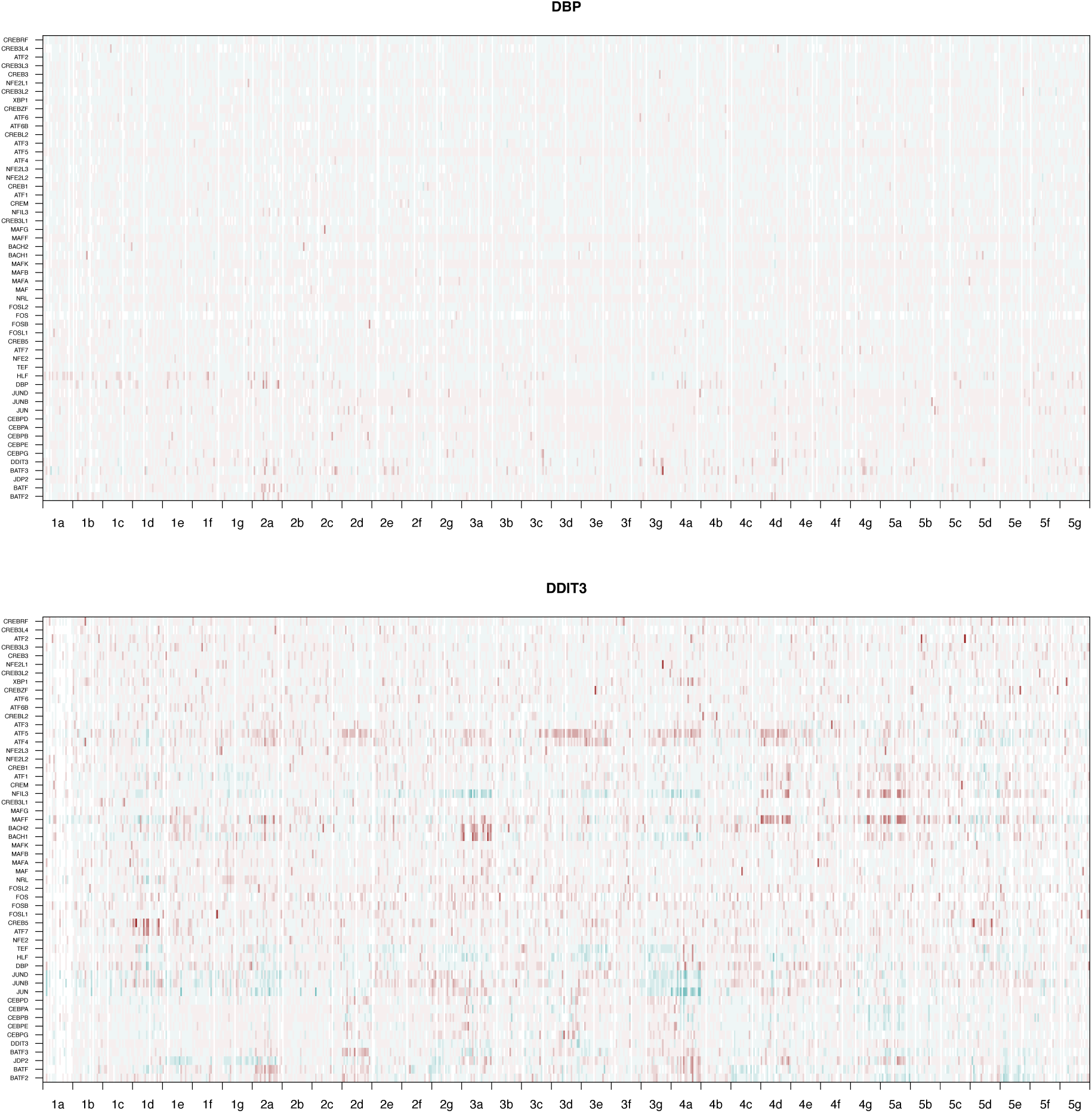
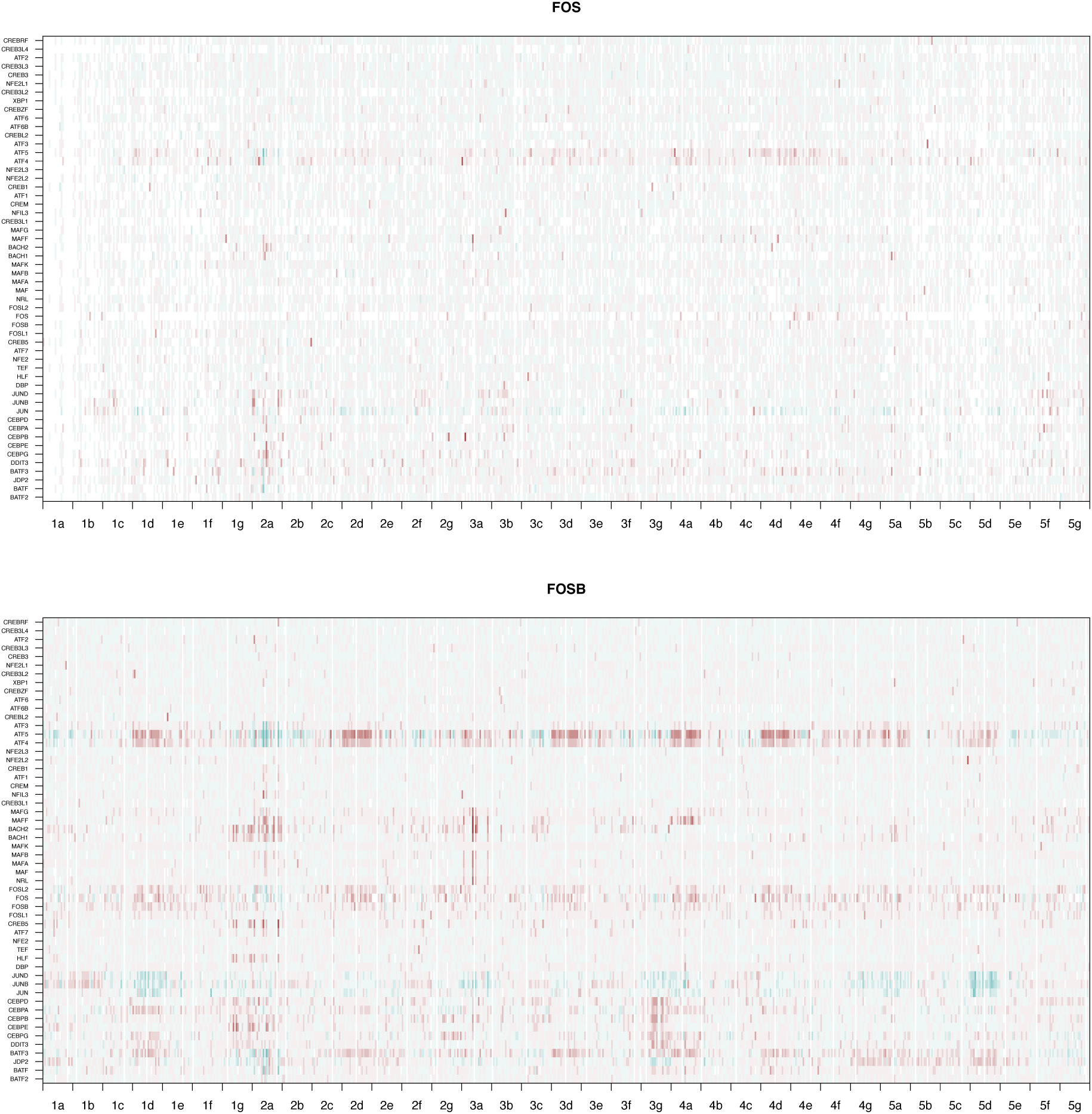
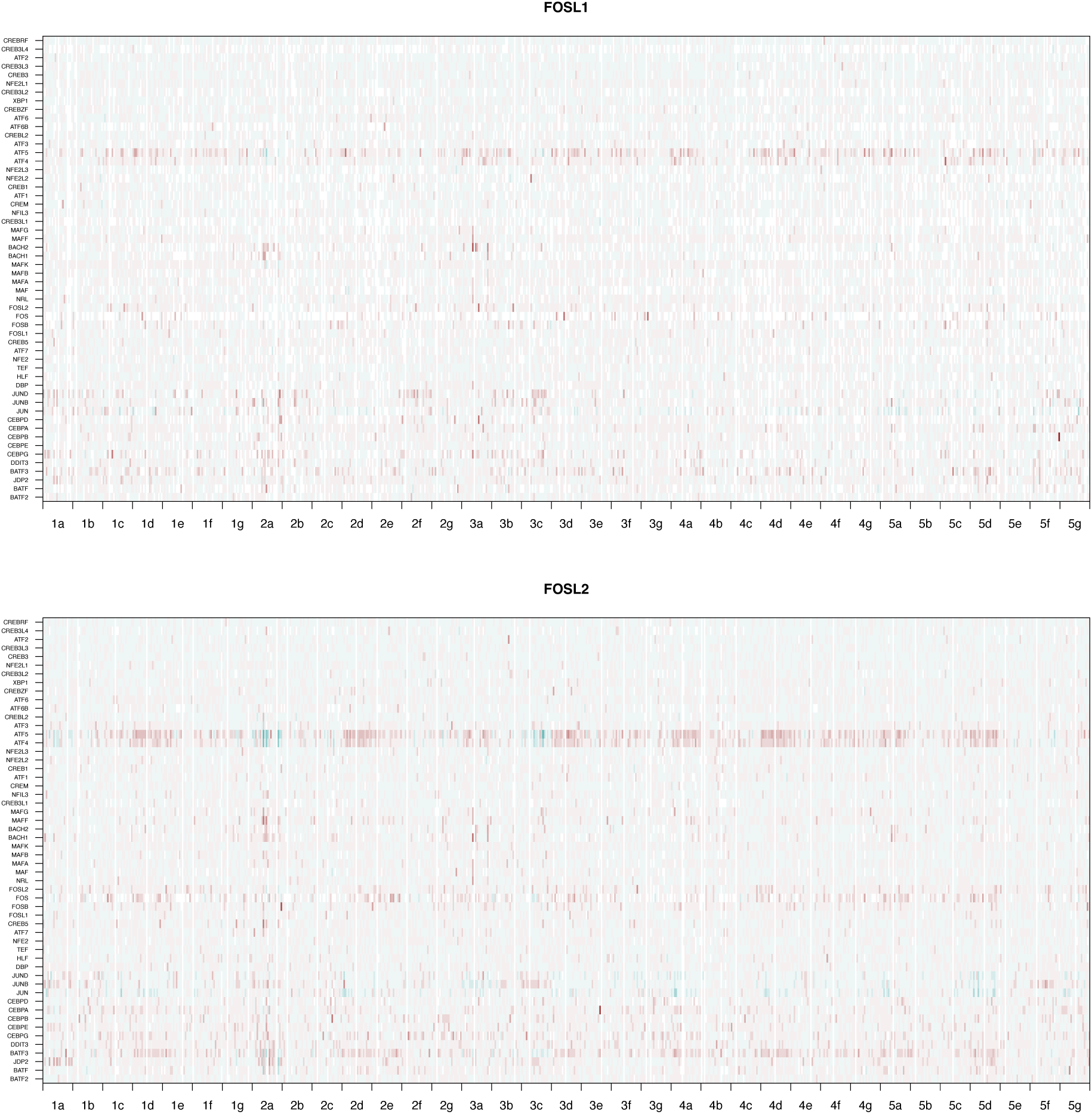
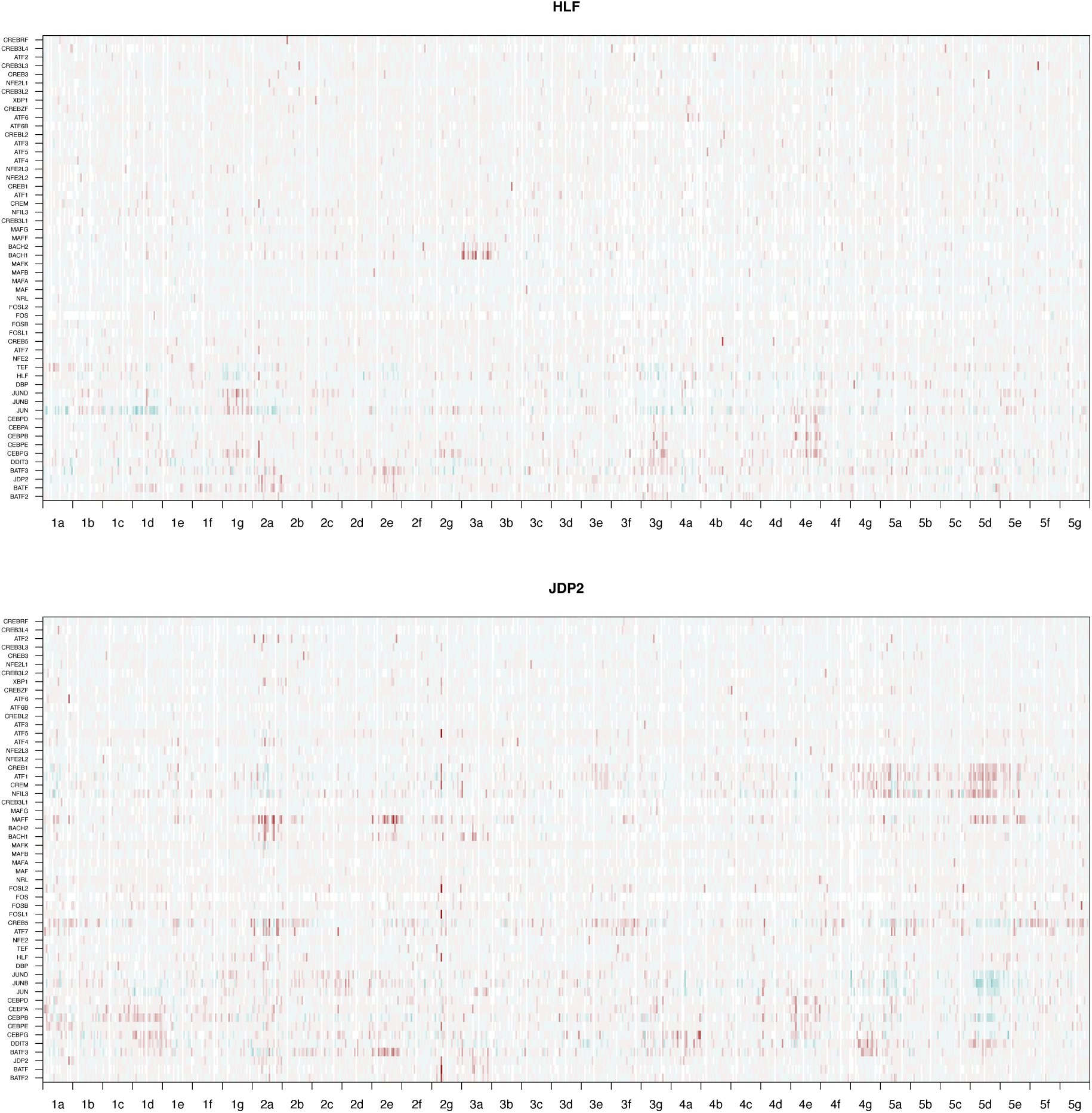
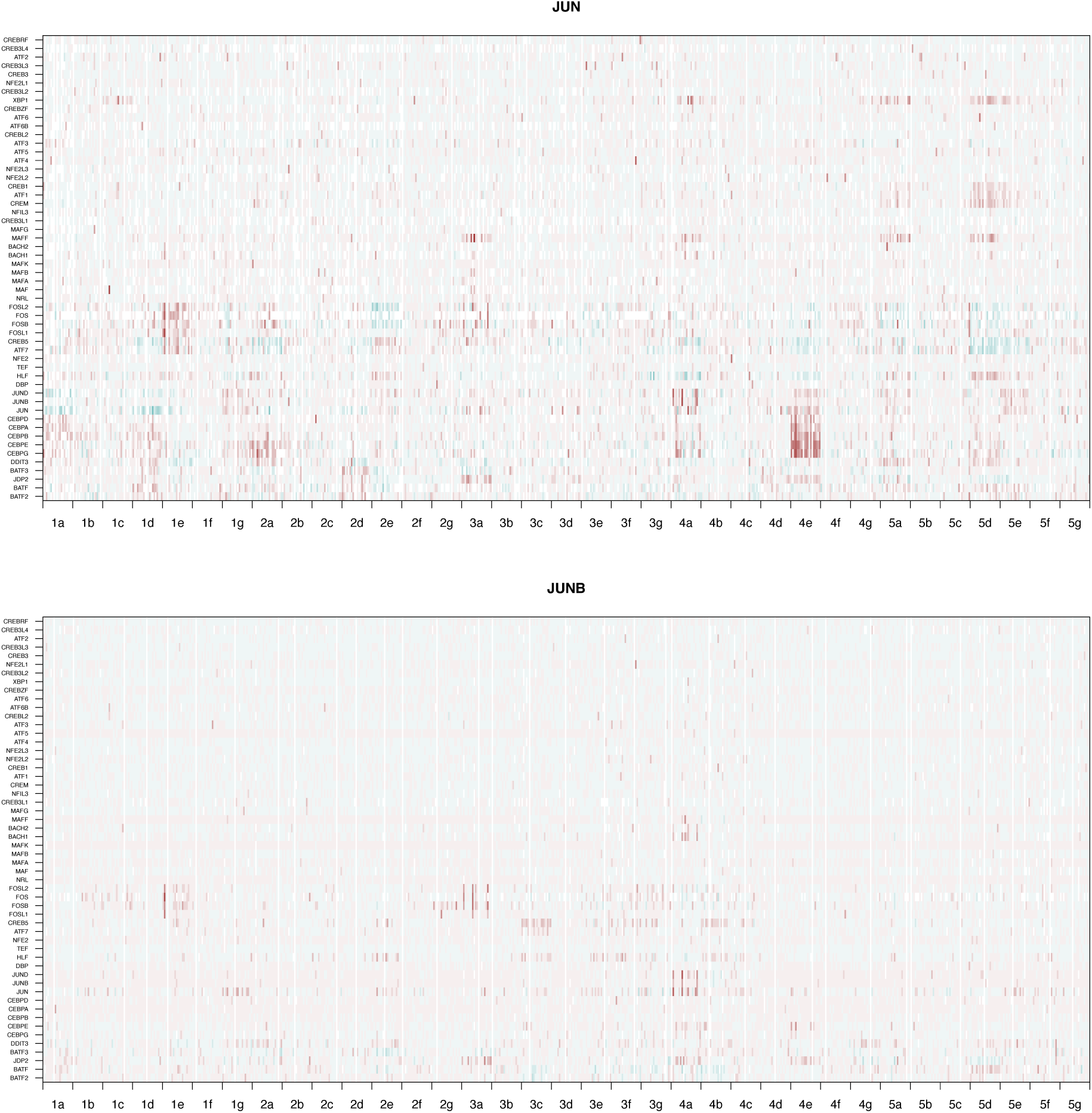
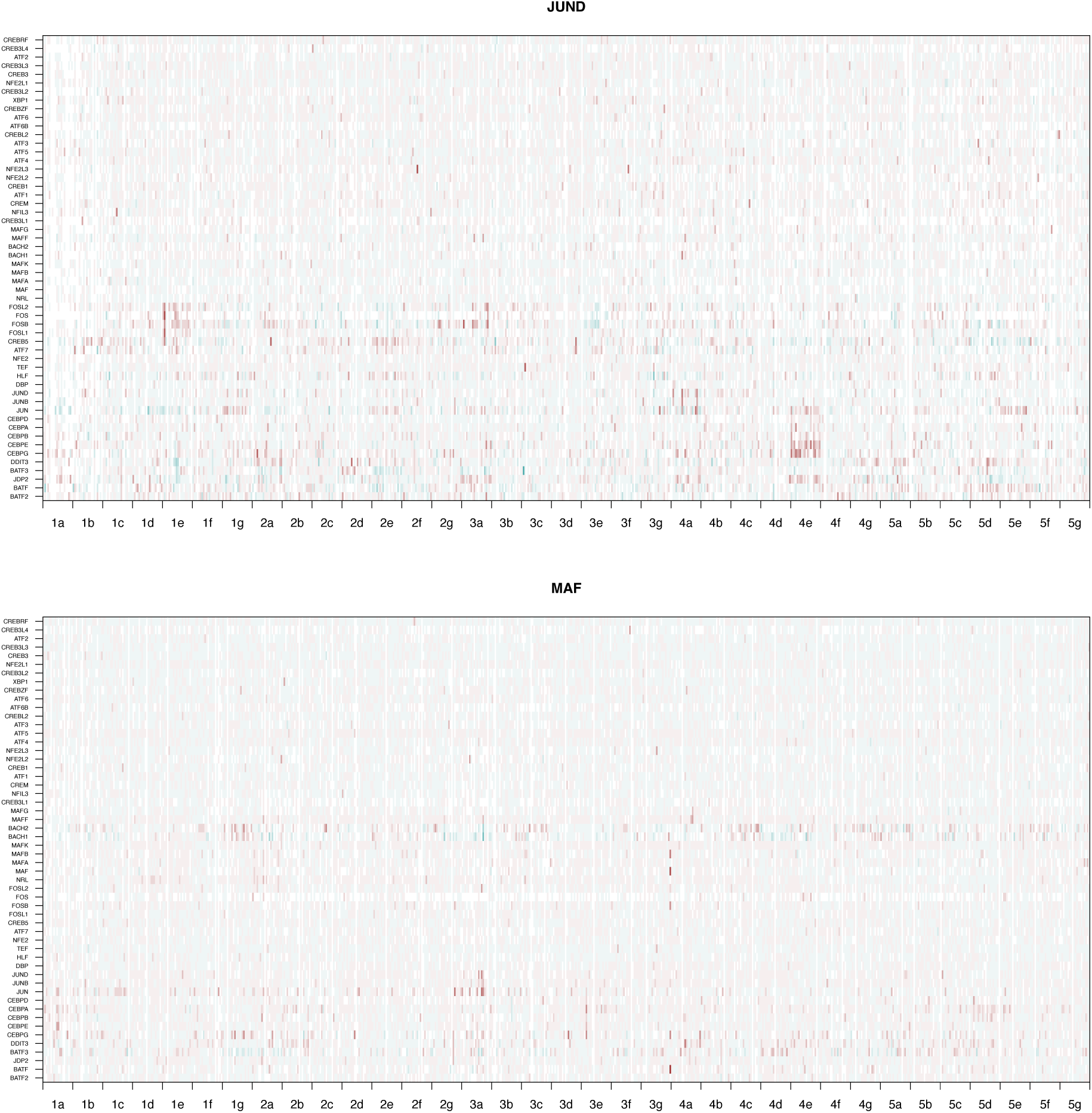
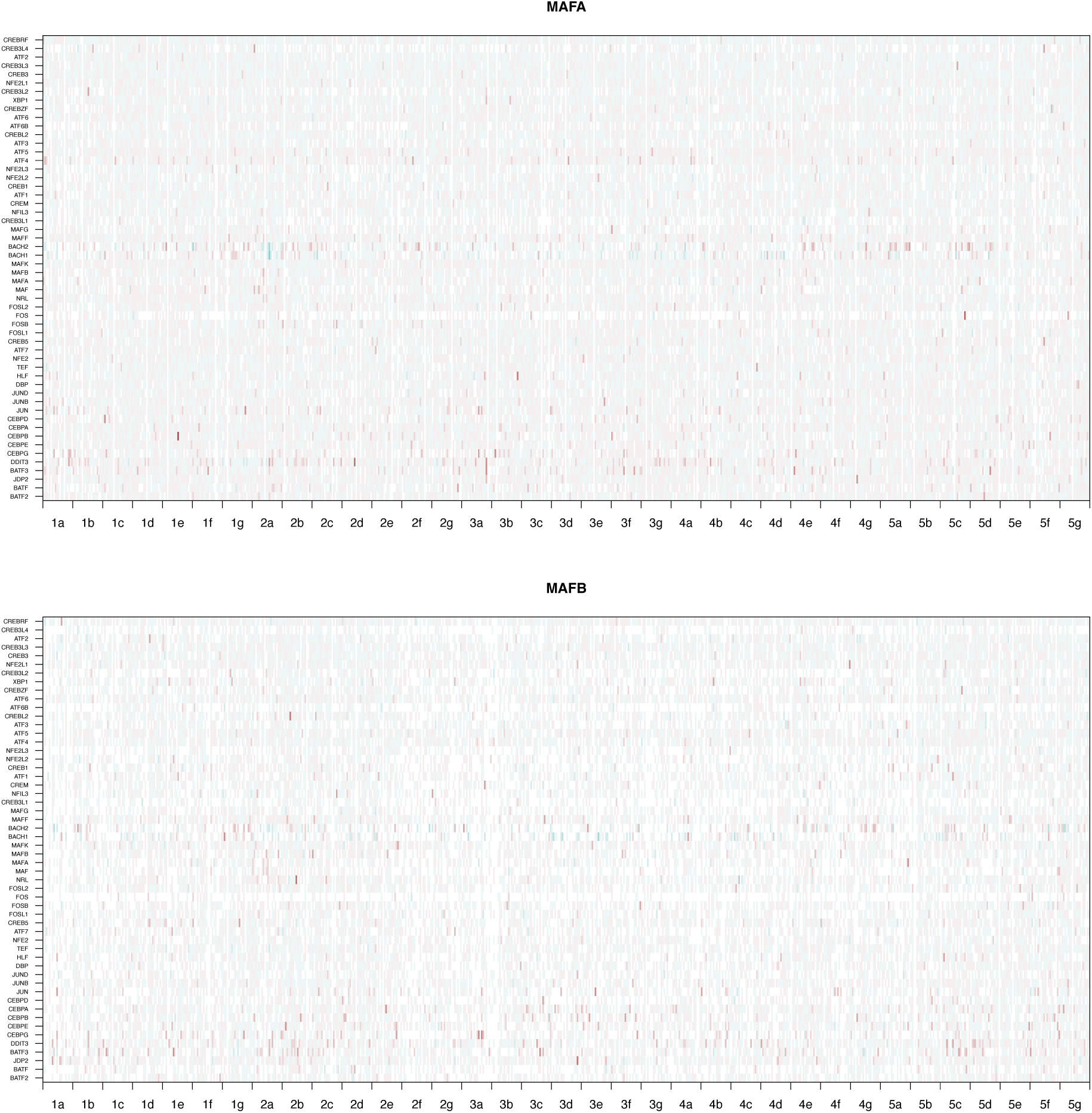
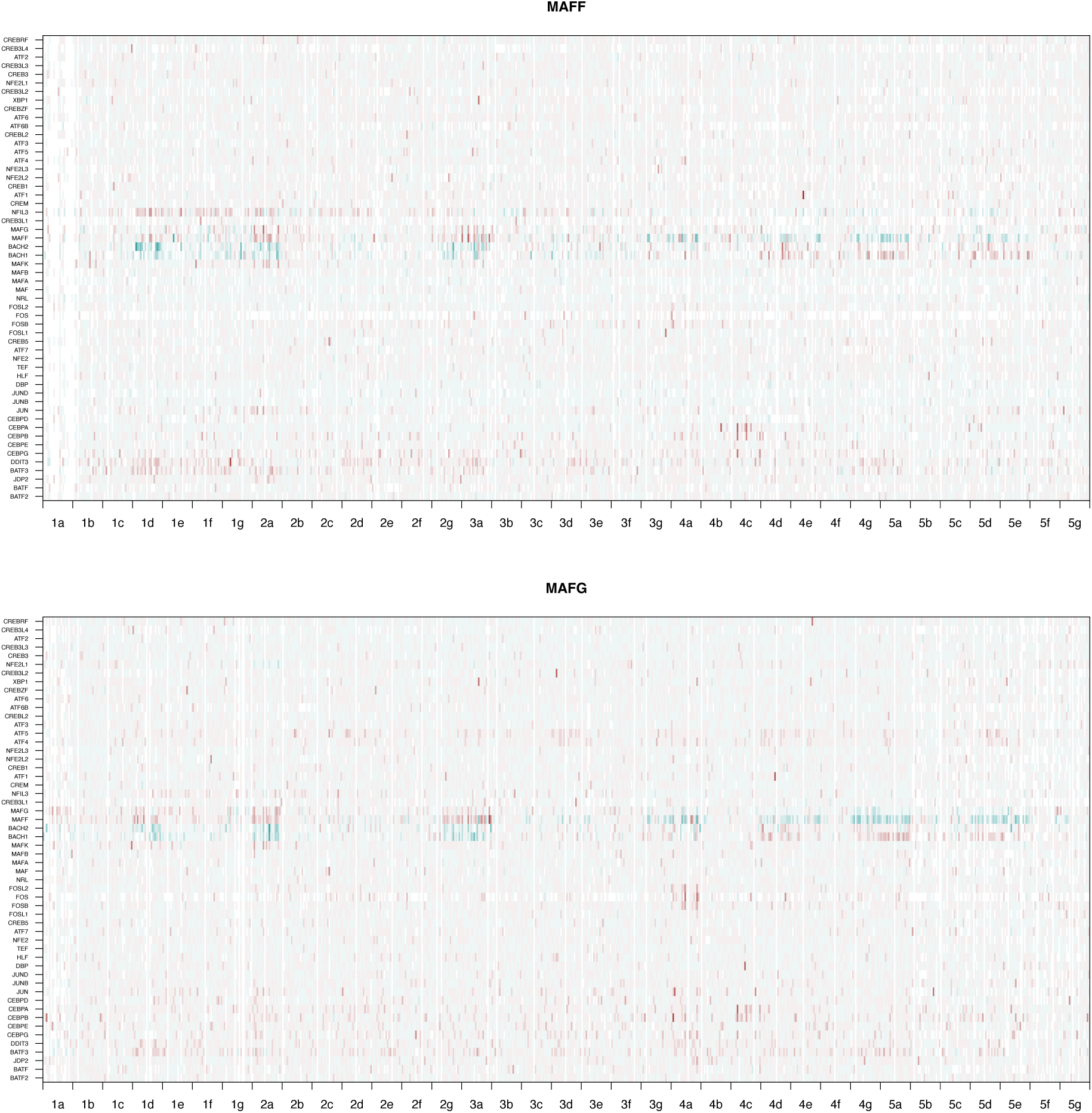
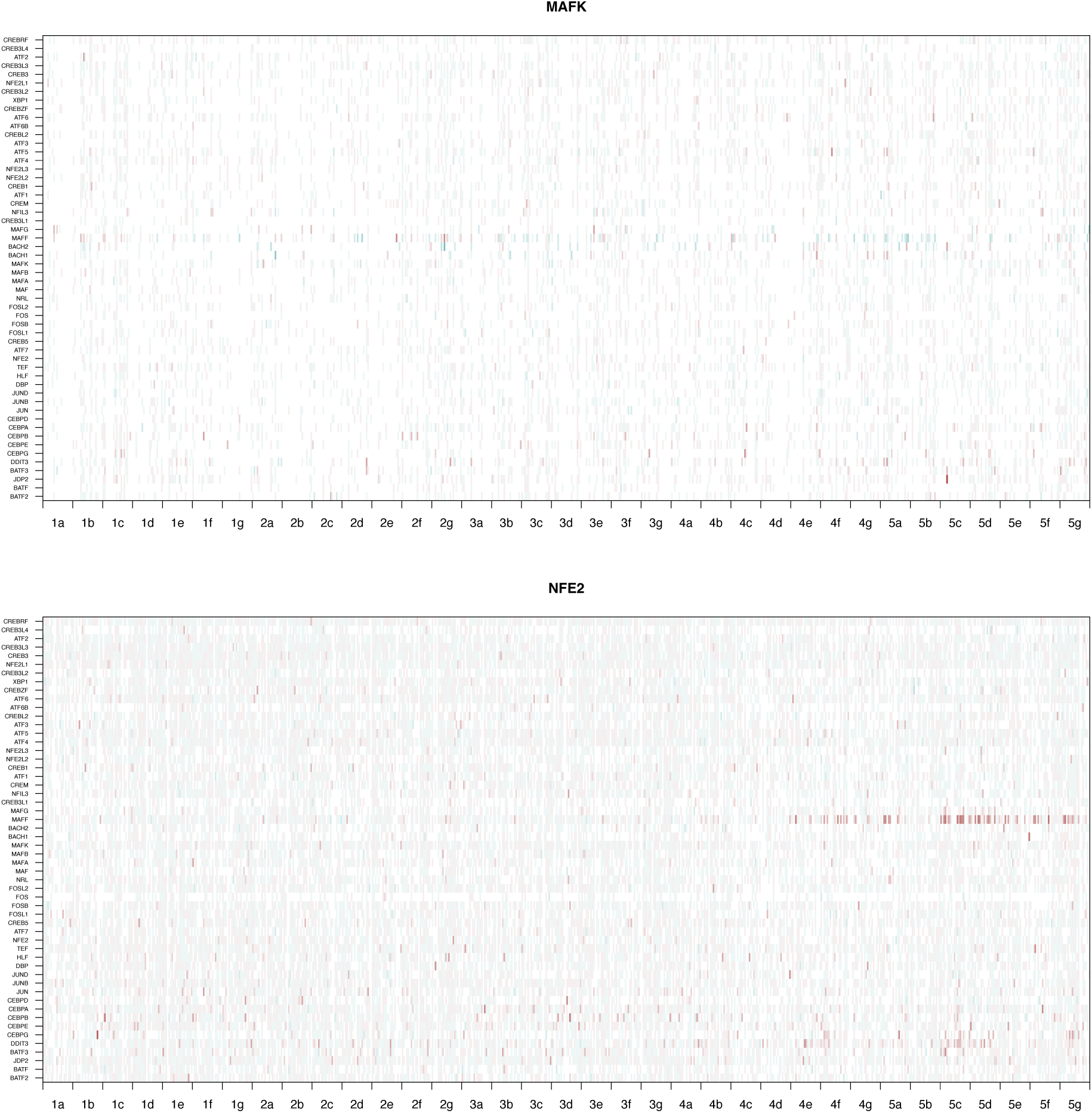
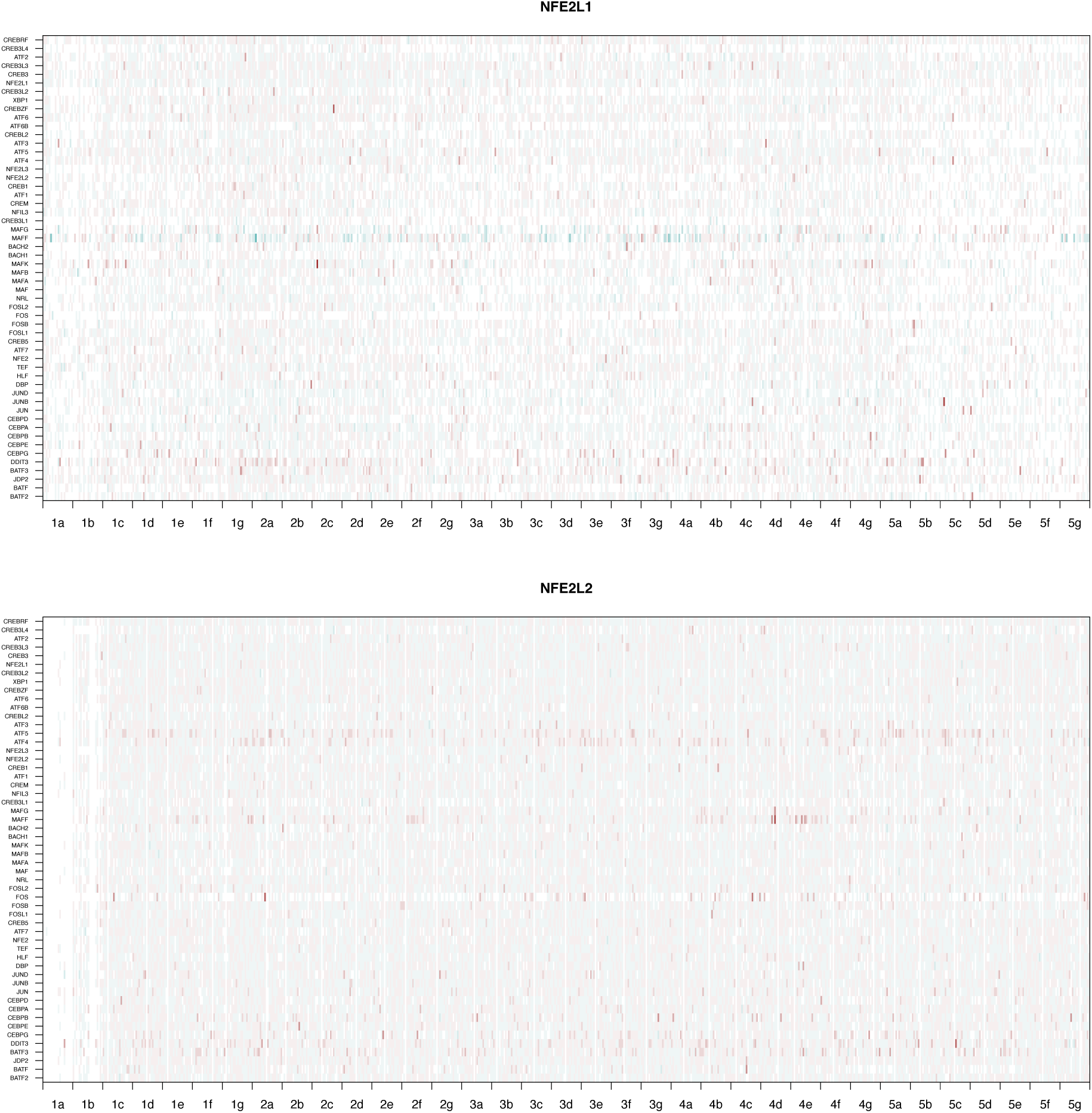
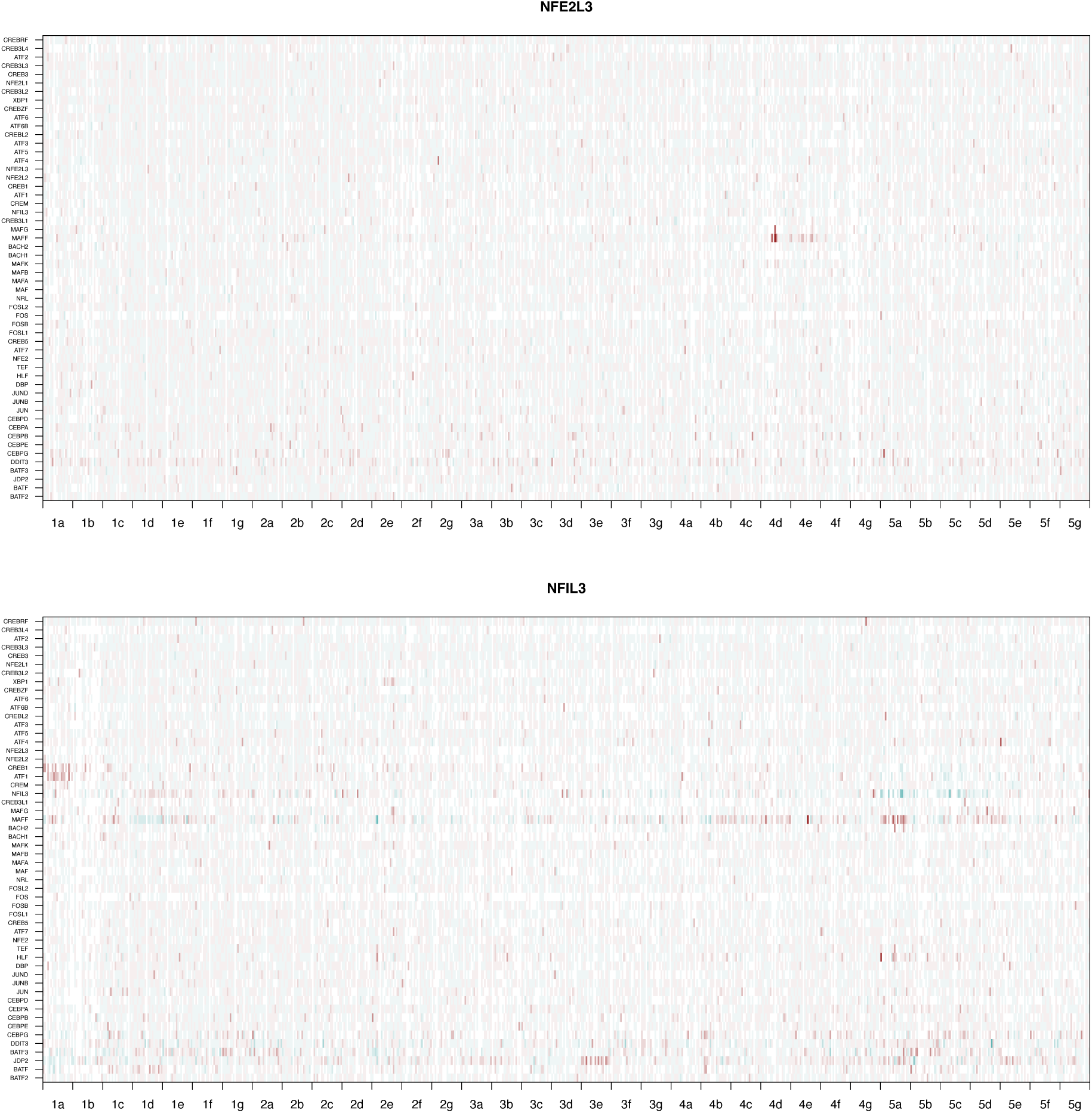
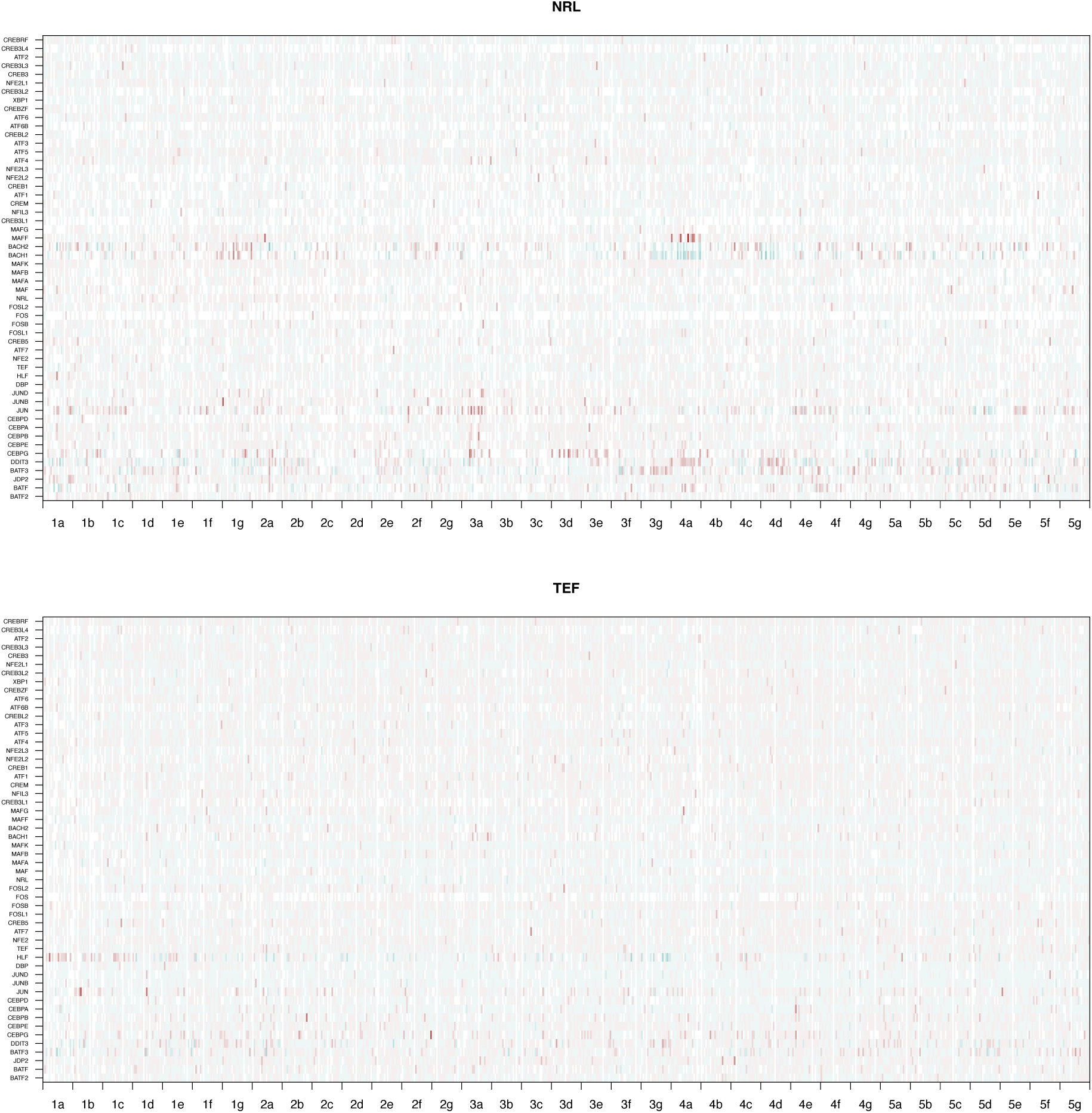
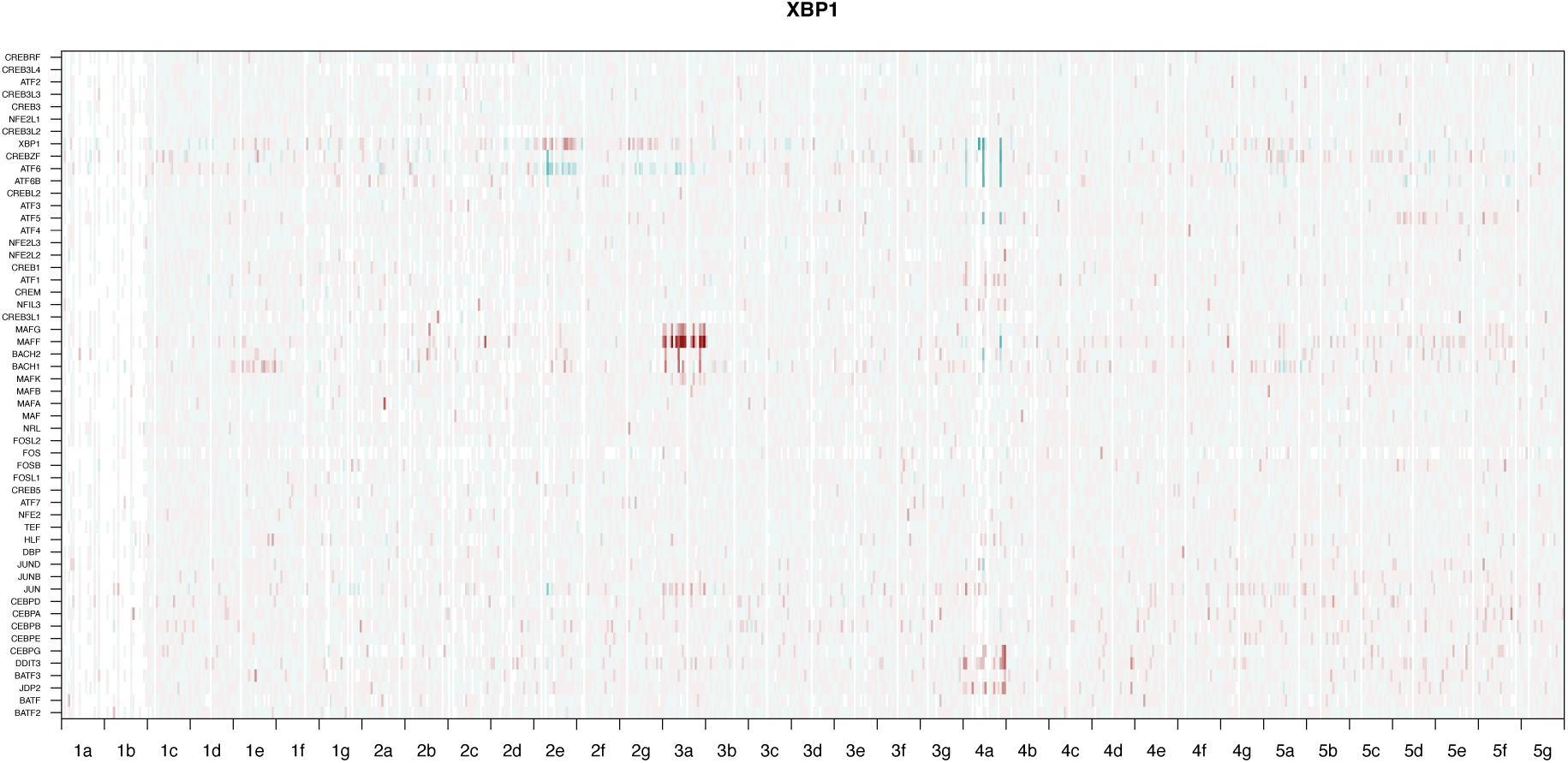
Heatmap of specific mutation effects (i.e. residuals from thermodynamic model) for all bZIPs. Row represent the 54 wild-type partners. Columns represent each of the 20 amino acids at each of the 35 positions. Negative specific effects (turquoise) indicate that the wild-type residue serves as a negative determinant of specificity. Positive specific effects (red) indicates that the wild-type residue serves as a negative determinant of specificity.

